# Morphology-Aware Profiling of Highly Multiplexed Tissue Images using Variational Autoencoders

**DOI:** 10.1101/2025.06.23.661064

**Authors:** Gregory J. Baker, Edward Novikov, Shannon Coy, Yu-An Chen, Clemens B. Hug, Zergham Ahmed, Sebastián A. Cajas Ordóñez, Siyu Huang, Clarence Yapp, Gaurav N. Joshi, Fumiki Yanagawa, Artem Sokolov, Hanspeter Pfister, Sandro Santagata, Peter K. Sorger

**Affiliations:** Laboratory of Systems Pharmacology, Harvard Medical School, Boston, MA; Ludwig Center for Cancer Research at Harvard, Harvard Medical School, Boston, MA; Department of Systems Biology, Harvard Medical School, Boston, MA; Harvard John A. Paulson School of Engineering and Applied Sciences, Harvard University, Cambridge, MA; Department of Pathology, Brigham and Women’s Hospital, Harvard Medical School, Boston, MA; Nikon Instruments, Lexington, MA

**Author notes:** Corresponding author **Corresponding Authors:** Gregory J. Baker, Ph.D., Pharm.D. Laboratory of Systems Pharmacology Harvard Medical School, Warren Alpert Building, Room 444, 200 Longwood Avenue, Boston, MA 02115, Peter K. Sorger, Ph.D., Laboratory of Systems Pharmacology Harvard Medical School, Armenise Building, Room 137F, 200 Longwood Avenue, Boston, MA 02115. Division of Oncological Sciences, Knight Cancer Institute, Oregon Health & Science University, Portland, OR. Visual Computing Division, School of Computing, Clemson University, Clemson, SC. Co-first author (Gregory J. Baker, Edward Novikov).

## Abstract

Spatial proteomics (highly multiplexed tissue imaging) provides unprecedented insight into the types, states, and spatial organization of cells within preserved tissue environments. To enable single-cell analysis, high-plex images are typically segmented using algorithms that assign marker signals to individual cells. However, conventional segmentation is often imprecise and susceptible to signal spillover between adjacent cells, interfering with accurate cell type identification. Segmentation-based methods also fail to capture the morphological detail that histopathologists rely on for disease diagnosis and staging. Here, we present a method that combines unsupervised, pixel-level machine learning using autoencoders with traditional segmentation to generate single-cell data that captures information on protein abundance, morphology, and local neighborhood in a manner analogous to human experts while overcoming signal spillover. We demonstrate the generality of this approach by applying it to CyCIF, Lunaphore COMET, and CODEX data, and show that it can learn histological features across multiple spatial scales.

## INTRODUCTION

High-plex tissue imaging (spatial proteomics) has emerged to study the biology of intact tissues and tumors by measuring the abundances and spatial distributions of tens to hundreds of biomolecules using fluorescently or metal-tagged antibodies^1^. Single-cell data are most commonly derived from these images by segmentation. This involves computing masks centered on nuclei that divide an image into individual cells. Staining intensities are then computed for each antibody to generate a *cell feature table*^2^ describing single-cell protein expression levels. A strength of this approach is that it facilitates multi-omic data integration by generating tabular data similar to flow cytometry data or count tables in single-cell transcriptional profiling^2^ .

However, segmentation of high-plex tissue images faces three fundamental limitations: spatial crosstalk (or lateral spillover), under and over segmentation, and insensitivity to morphology. The former arises when signals from neighboring cells cross-contaminate each other because they cross segmentation boundaries^3^, which confounds cell type identification and other analyses.^3,4^ Recent high-plex, high-resolution, 3D imaging of tissues^5^ has shown that cells have membrane-membrane distances <1 μm, their processes are often intertwined^6^, and two or more cells commonly overlap along the imaging (Z) axis. These observations explain the inability of even state-of-the-art algorithms such as CellPose^7^ (and its recent updates)^8^ to fully overcome spatial crosstalk or related problems such as over-segmentation (subdividing cells into multiple objects) or under-segmentation (combining different cells into a single object).

Differences in cell morphology play a major role in classical histopathology, most commonly based on human inspection of hematoxylin and eosin (H&E) stained slides, but this is another area in which single cell analysis based on tissue segmentation falls short. Differences in cell morphology can be detected using supervised and unsupervised machine learning (ML) from pixel-level image data, for example using computational pathology (CPath) AI/ML approaches.^9^ CPath methods use convolution neural networks, transformers, and other deep learning approaches to classify histology images by tissue type, disease state, or treatment response.^10^ Pixel-level learning has also been applied to multiplexed tissue images using convolutional and transformer-based neural networks, CNN-based classifiers^11–14^, generative models^15,16^, and vision transformers^17,18^, and unsupervised clustering of image features^19^. Supervised learning on multiplexed image data shows promise^18^ but it is unclear how model outputs compare to single cell data generated using traditional segmentation methods.

In this paper we describe an approach to spatial profiling that combines image segmentation and pixel-level ML using light-weight generative models known as variational autoencoders (VAEs)^20,21^; we then compare the results to what can be obtained using optimized segmentation approaches alone. We compile these complementary methods in the MORPHAEUS toolbox to facilitate reuse and further development. We find that VAEs can be trained to analyze high-plex images in a fully unsupervised manner using single-cell or multicellular image patches. Clustering of VAE latent space manifolds then enables the identification and cataloging of phenotypes at both the single-cell and tissue levels, allowing the model to learn histological features across spatial scales, from single cells to recurrent cell assemblies.

## RESULTS

### Spatial crosstalk limits accuracy of segmentation-based cell phenotyping

Image segmentation and pixel-level learning are agnostic to how images are acquired and both methods have been applied to a wide variety of fluorescence-based spatial profiling technologies including CODEX^23^, COMET (Lunaphore)^24^, cyclic immunofluorescence (CyCIF)^25^ and other spatial proteomic methods^1^. In this paper we focus on CyCIF datasets collected as reference data^26^ for the Human Tumor Atlas Network (HTAN)^27^ and demonstrate that the approach generalizes to COMET and CODEX datasets.

Analysis of CyCIF images started with a 22-plex whole-slide CRC dataset previously released by HTAN (HTAN ID: HTA13_1_101) and referred to here as CyCIF-1A (**Fig. 1a** and **Supplementary Table 1**). Algorithms in MCMICRO^28^ were used to perform image assembly and segmentation. An H&E image from a tissue section adjacent to CyCIF-1A (HTAN ID: HTA13_1_100) was annotated by two board-certified pathologists and found to comprise 11 distinct histological regions (HRs) of diagnostic or biological significance such as normal mucosa (the crypt-containing epithelial layer of the colon), smooth muscle and stromal tissue (**Fig. 1b,c**) as well as multiple adenocarcinoma (AC) morphologies including glandular, mucinous, and solid.^29^

**Fig. 1.**
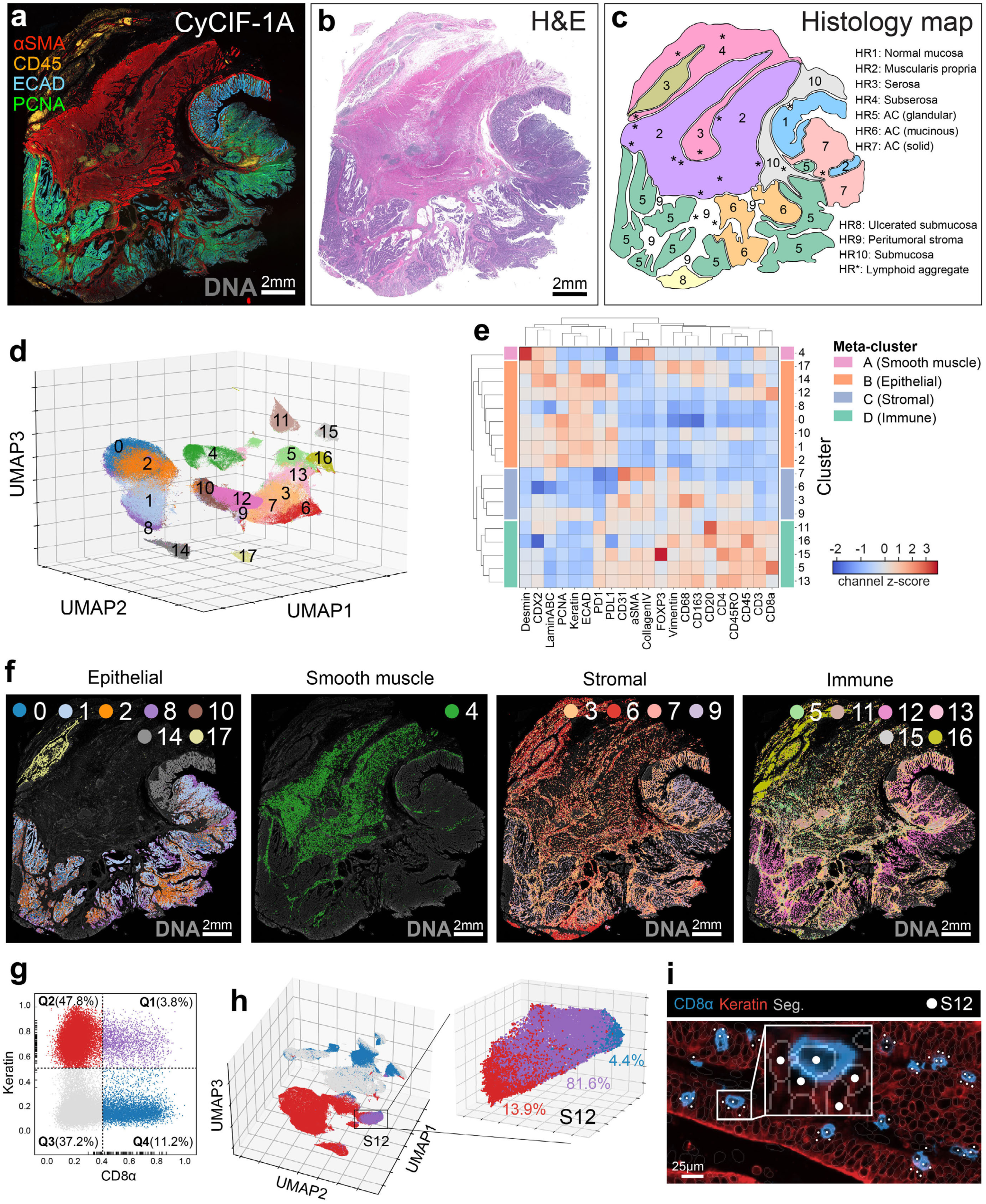
Spatial crosstalk limits accuracy of segmentation-based cell phenotyping. **a**, CyCIF-1A: a 22-plex, whole-slide CyCIF image of primary human CRC (HTAN ID: HTA13_1_101) showing five channels: αSMA (smooth muscle, red), CD45 (leukocytes, orange), ECAD (epithelium, blue), PCNA (proliferating cells, green), and DNA (Hoechst-stained nuclei, gray). See **Supplementary Table 2** for a complete list of antibodies used to label this tissue. **b**, Adjacent tissue section to CyCIF-1A (∼4µm away in Z-axis) stained with routine histology dyes Hematoxylin & Eosin (H&E). **c**, Histology map showing the locations of 11 histological regions (HRs) observed by two board-certified pathologists in the H&E image shown in panel b. **d**, UMAP embedding of segmented cells from CyCIF-1A colored by Leiden cluster; see **Supplementary Video 1** for 360° plot rotation. **e**, Hierarchical clustering of Leiden cluster z-score-normalized mean marker intensities revealing four primary meta-clusters (A-D). **f**, Location of Leiden clusters in CyCIF-1A grouped by major cell lineages: Epithelial, Smooth muscle, Stromal, and Immune. **g**, Gated bivariate plot showing the percentage of CD8α^+^ keratin^+^ (Q1, purple), keratin^+^ (Q2, red), CD8α^-^ keratin^-^ (Q3, gray), and CD8α^+^ cells (Q4, blue) in CyCIF-1A. Rugplots (small black ticks along x- and y-axes) indicate marker intensity values for 100 cells drawn at random from S12 showing that the majority of cells in this cluster are positive for both markers. **h**, UMAP embedding of segmented cells from CyCIF-1A colored by the gating strategy shown in panel g; cluster S12 is highlighted by the black rectangle. Inset shows S12 in further detail highlighting its percentages of keratin^+^ (red), CD8α^+^ (blue), and CD8α^+^ keratin^+^ cells (purple). **i**, Field-of-view from CyCIF-1A showing that cells in S12 (white scatter points) consist of both CD8α^+^ T cells (blue) and neighboring keratin^+^ epithelial cells (red). Inset shows spillover of marker signals across segmentation boundaries (translucent perimeters) between the two different cell types, which we refer to as “spatial crosstalk”.

To generate single-cell data from these images, per-cell mean fluorescence intensities were computed on segmented cells using an object mask appropriate for each protein and corresponding to the cell nucleus, cytoplasm, or plasma membrane (e.g. a plasma membrane mask for CD4 and CD8, and nuclear mask for PCNA; **Supplementary Table 2**). The resulting single-cell feature table underwent quality control (QC) with CyLinter^30^ to remove visual and image processing artifacts (affecting ∼21% of total cells; **Extended Data Fig. 1a**). For an extended discussion of segmentation-based tissue analysis see (**Supplementary Note 1** and **Supplementary Fig. 1**).

Clustering of post-QC segmented data using Leiden community detection^31^ yielded 18 clusters (S0-17, with “S” denoting segmentation-based). Clusters ranged in size from 0.8% to 12.6% of total cells (**Supplementary Table 3**). Horn’s parallel analysis^32^ showed that the first three principal components (PCs) captured the majority (68%) of dataset variance; single cell data were therefore visualized in 3D UMAP^33^ embeddings (**Fig. 1d**, **Extended Data Fig. 1b** and **Supplementary Video 1**). Hierarchical clustering (using mean, z-score-normalized, signal intensities) revealed four major cell types occupying characteristic regions of tissue: keratin^+^ epithelium, ⍺SMA^+^ smooth muscle, vimentin^+^ stroma, and CD45^+^ immune cells (**Fig. 1e,f**, **Extended Data Fig. 1c-e** and **Supplementary Fig. 2**). Clusters S0, 1, 2, 8, and 10 represented four tumor cell subsets that differed in PCNA expression (**Extended Data Fig. 1f**) but did not correspond directly to glandular, mucinous, and solid AC histologies (**Fig. 1b,c and f**). Normal epithelial cells comprised the tumor-adjacent colonic mucosa (S14, HR1) and benign reactive mesothelium in the serosa (S17, R3), the latter likely arising from tumor-associated inflammation. Smooth muscle occupied the muscularis propria (S4, HR2), while fibroblasts (S6) were identified in the subserosa (HR4), peritumoral stroma (HR9), and submucosa (HR10). Endothelial cells (S7) were present throughout the tissue, as were immune cells, including populations of macrophages (S3), CD8^+^ T cells (S5), and both conventional (S13) and regulatory (S15) CD4^+^ T cells (**Extended Data Fig. 2** and **Supplementary Fig. 3**).

Despite the biological plausibility of these cell phenotypes, closer examination revealed multiple clusters with unlikely immunomarker combinations. For example, cells in S12 appeared to express markers of proliferating epithelial cells (keratin, ECAD, PCNA) as well as CD8^+^ T cells (CD45, CD3ε, CD8α, CD45RO; **Fig. 1e**). This was not a consequence of suboptimal clustering, since manual gating on keratin and CD8α channels showed that >80% of segmented cells in S12 were positive for both markers (**Fig. 1g,h**) and inspection of the underlying image showed that they corresponded to CD8^+^ T cells immediately adjacent to keratin^+^ tumor cells (**Fig. 1i**). We conclude that S12 cells represent CD8^+^ T cells affected by spatial crosstalk from tumor cells (**Supplementary Fig. 4**).

Clusters S11 and S16 were also affected by spatial crosstalk. Both clusters represented tightly packed B and T cells in lymphoid aggregates (HR*), with S16 cells found exclusively in the serosa (HR3) and subserosa (HR4). Non-tumor infiltrating CD8^+^ T cells in S5 were impacted by spatial crosstalk as well, but this cell type was distributed across multiple additional subclusters in UMAP space. A detailed investigation of the relationship between spatial crosstalk and dimensionally reduced embeddings (UMAP, tSNE) showed that spatial crosstalk: (i) was pervasive, affecting up to 30% of the ∼1M cells in CyCIF-1A; (ii) promoted formation of non-biological clusters due to distinctive (yet artifactual) marker combinations and (iii) ii) represented a significant barrier to accurate cell phenotyping (**Supplementary Note 2** and **Supplementary Figs. 5 & 6**)

### VAE classification of human CRC at single-cell resolution

To analyze the same CyCIF data at the pixel level we developed a method for training VAE models on image patches derived from whole-slide data. This approach used Cellcutter^34^ to generate 9x9µm (14x14 pixel) multi-channel patches centered on individual nuclei (as determined by image segmentation) and store them in a memory-efficient Zarr file (**Fig. 2a,b** and **Methods**). After subsampling, the final dataset for CyCIF-1A consisted of 479,909 patches, 86% with one and 13% with two nuclei (**Fig. 2c**). Image patch pre-processing involved: (i) log transformation, (ii) clipping to remove bright and dim outlier pixels, (iii) per-channel normalization (0-1) based on fluorescence intensities from the whole-slide image, and (iv) rotational data augmentation to reduce sensitivity to cell orientation (**Extended Data Fig. 3a,b**). Patches were divided into training, validation, and test sets at a ratio of 80:10:10 and the first set used to train a standard VAE network consisting of four convolutional encoding layers followed by a dense layer (**Fig. 2d**, **Extended Data Fig. 3c**, and **Methods**). To focus the attention of the network on the cell at the center of the patch, each channel was convolved with a 2D Gaussian kernel having a standard deviation of 7 pixels. This vignetting technique reduced the intensity of features towards patch borders, causing the network to perceive them with less acuity (42% of maximum intensity at patch corners; **Fig. 2e**).

**Fig. 2.**
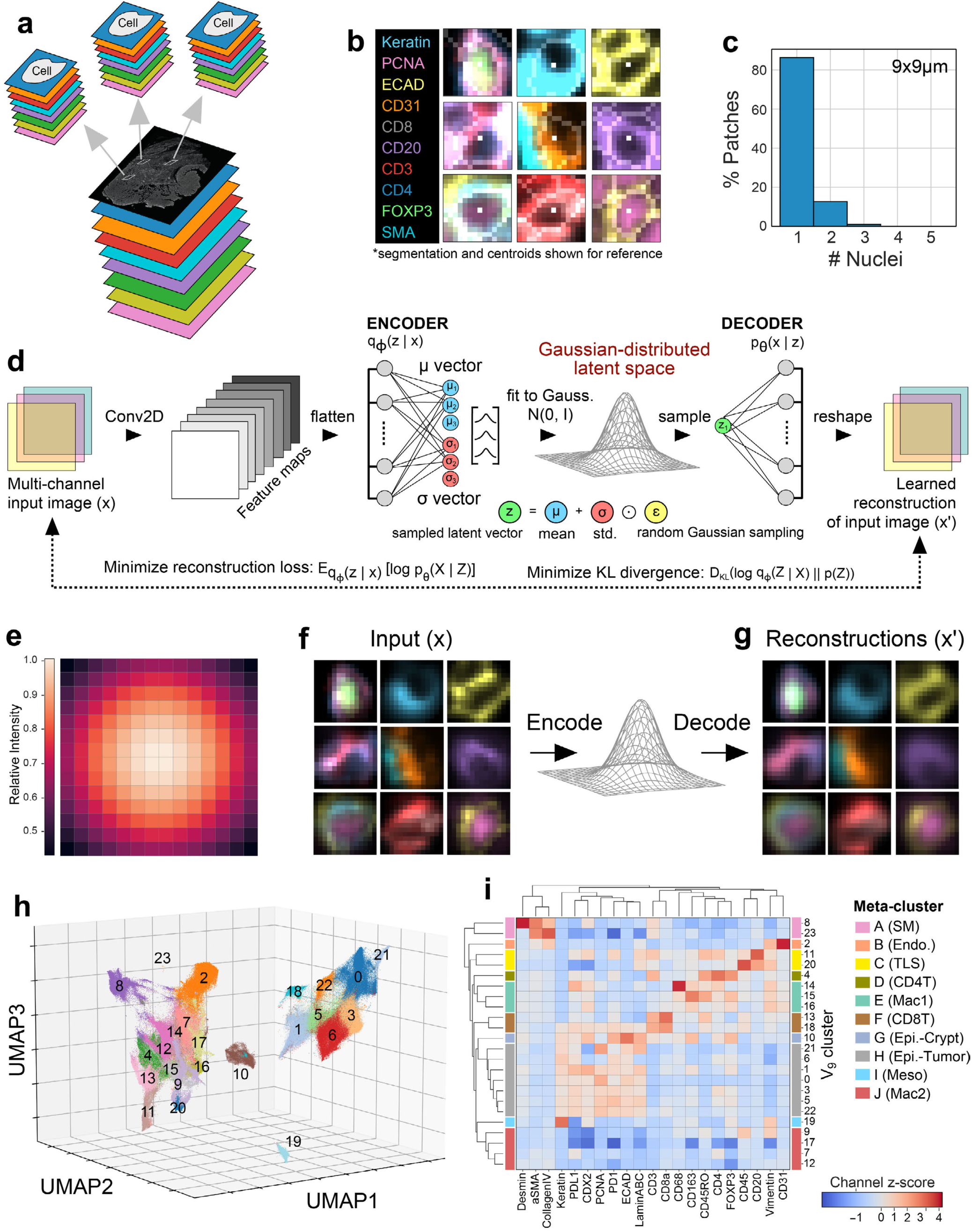
VAE-based classification of human CRC at single-cell resolution. **a**, Schematic showing the process of extracting multi-channel, single-cell image patches from whole-slide multiplex tissue images using Cellcutter. **b**, Example 9x9µm multi-channel image patches extracted from CyCIF-1A. Nuclear centroids of reference cells at the center of each patch (white pixels) and segmentation boundaries (translucent perimeters) shown for reference only and not used during VAE model training or inference. **c**, Histogram showing the percentage of 9x9µm image patches from CyCIF-1A containing different numbers of cell nuclei (median = 1, Q1 = 1, Q3 = 1, IQR = 0). **d**, Schematic of a standard VAE network trained on multi-channel image patches. The encoding network is trained to map vector representations of input images onto a lower-dimensional, Gaussian-distributed latent space manifold. Image patch encodings are passed through the decoding network to visualize learned reconstructions whose similarity to corresponding input images is assessed using the average of two metrics: the mean squared error (MSE) and Kullback–Leibler (KL) divergence from a multi-dimensional Gaussian distribution. **e**, Heatmap showing relative pixel intensities of a 2D Gaussian-distributed vignette mask with a standard deviation of 7-pixels applied per-channel and used to focus the attention of the VAE network on the reference cell at the center of each 9x9µm image patch. **f**, Image patches shown in panel b after vignette masking, which are input into the VAE encoding network. **g**, Learned reconstructions of vignetted input images shown in panel f showing that information on protein expression and morphology have been learned by the network. **h**, 3D UMAP embedding of 9x9µm VAE image patch encodings from CyCIF-1A colored by Leiden cluster; see **Supplementary Video 2** for 360° plot rotation. **i**, Hierarchical clustering of z-score-normalized mean pixel-level marker intensities for 9x9µm VAE image patch Leiden clusters revealing 10 meta-clusters (A-J).

A VAE treats individual image pixels as independent features (or dimensions). Thus, 9x9µm image patches from our 21-plex CRC image comprised 4,116 total dimensions (14 pixels x 14 pixels x 21 channels). The VAE latent space dimension (an adjustable parameter influencing model accuracy) was set to 10% of total image patch dimensionality, resulting in a 412-dimensional latent space. Visual examination of learned VAE reconstructions confirmed accurate capture of marker location and intensity and exclusion of idiosyncratic differences among cells of the same type (**Fig. 2 f, g**). At convergence, the model achieved a validation loss of 8.3x10^-4^.

Leiden clustering of VAE image patch encodings generated 24 clusters (V_9_0-23, with “V_9_” denoting a VAE trained on 9x9µm patches (**Fig. 2h**, **Supplementary Table 4**, **Supplementary Video 2**, and **Supplementary Image Gallery 1**) and these corresponded to the primary lineages identified by segmentation-based analysis: keratin^+^ epithelium, ⍺SMA^+^ smooth muscle, vimentin^+^ stroma, and CD45^+^ immune cells (**Extended Data Fig. 4a,b** and **Supplementary Fig. 7**). Primary clusters could be grouped into 10 meta-clusters (A-J; **Fig. 2i**) by hierarchical clustering based on median, z-score normalized, pixel-level marker intensities. To evaluate the agreement between VAE and segmentation-based cluster pairs, we used the Jaccard index (JI). Six clusters (29% of total cells) exhibited substantial overlap (JI ≥ 0.44). These comprised Tregs (V_9_4/S15), smooth muscle cells (V_9_8/S4), normal crypt-forming epithelial cells (V_9_10/S14), lymphoid aggregates (V_9_11/S11), CD8^+^ T cells (V_9_13/S5), and mesothelial cells (V_9_19/S17; **Extended Data Fig. 4c,d**, **Extended Data Fig. 5** and **Supplementary Fig. 8**). Since segmentation-based clustering is based on marker intensity, we surmised that formation of these VAE clusters was also driven primarily by differences in marker expression. To demonstrate that VAE-based cell state inference generalizes across spatial proteomics platforms despite differences in image bit depth, resolution, and tissue type, we performed similar analysis of images of non-small cell lung cancer collected using COMET and head & neck squamous cell carcinoma collected using CODEX (**Supplementary Note 3**, **Extended Data Fig. 6**, and **Supplementary Figs. 9 & 10**).

### VAE image patch classification is morphology-aware and overcomes spatial crosstalk

VAE-based learning identified seven tumor cell clusters in CyCIF CRC data (meta-cluster H; **Fig. 2i**), compared to five with segmentation. While marker expression profiles for tumor-associated segmentation and VAE clusters were similar (**Fig. 3a**), only VAE-based clusters mapped to distinct physical regions of the tumor (**Fig. 3b,c**, **Extended Data Fig. 7a** and **Fig. 1b,c**). To quantify this correspondence, we calculated normalized mean posterior probabilities for VAE clusters relative to four previously developed kNN classifiers trained to predict normal, glandular, mucinous, or solid AC histologies from CyCIF data^35^. Cells in clusters V_9_0, 3, 5, 21, and 22 showed high mean probabilities of glandular AC (≥71%), while those in clusters V_9_6 and V_9_1 had the highest probabilities of mucinous (54%) and solid AC (47%), respectively. These results suggested that the VAE learned to distinguish tumor cells with different morphologies, and that mucinous and solid AC subtypes exhibit greater morphological variability than glandular AC (**Fig. 3d** and **Extended Data Fig. 7b**). To quantify the degree of uncertainty in AC subtype assignment, we calculated the information entropy of VAE clusters relative to their kNN class probabilities (**Fig. 3d**, red dotted line). Coloring clusters by entropy scores and mapping them back onto CyCIF-1A, we observed a distinctive pattern, with lower entropy glandular AC appearing at the left of the tissue and higher entropy mucinous and solid AC at the right (**Fig. 3e**). This result revealed a spatial gradient of tumor dedifferentiation across the specimen. Inspection by pathologists confirmed that the VAE had correctly learned differences in tumor phenotypes in an unsupervised manner and was superior to kNN classifiers generated by supervised training.

**Fig. 3.**
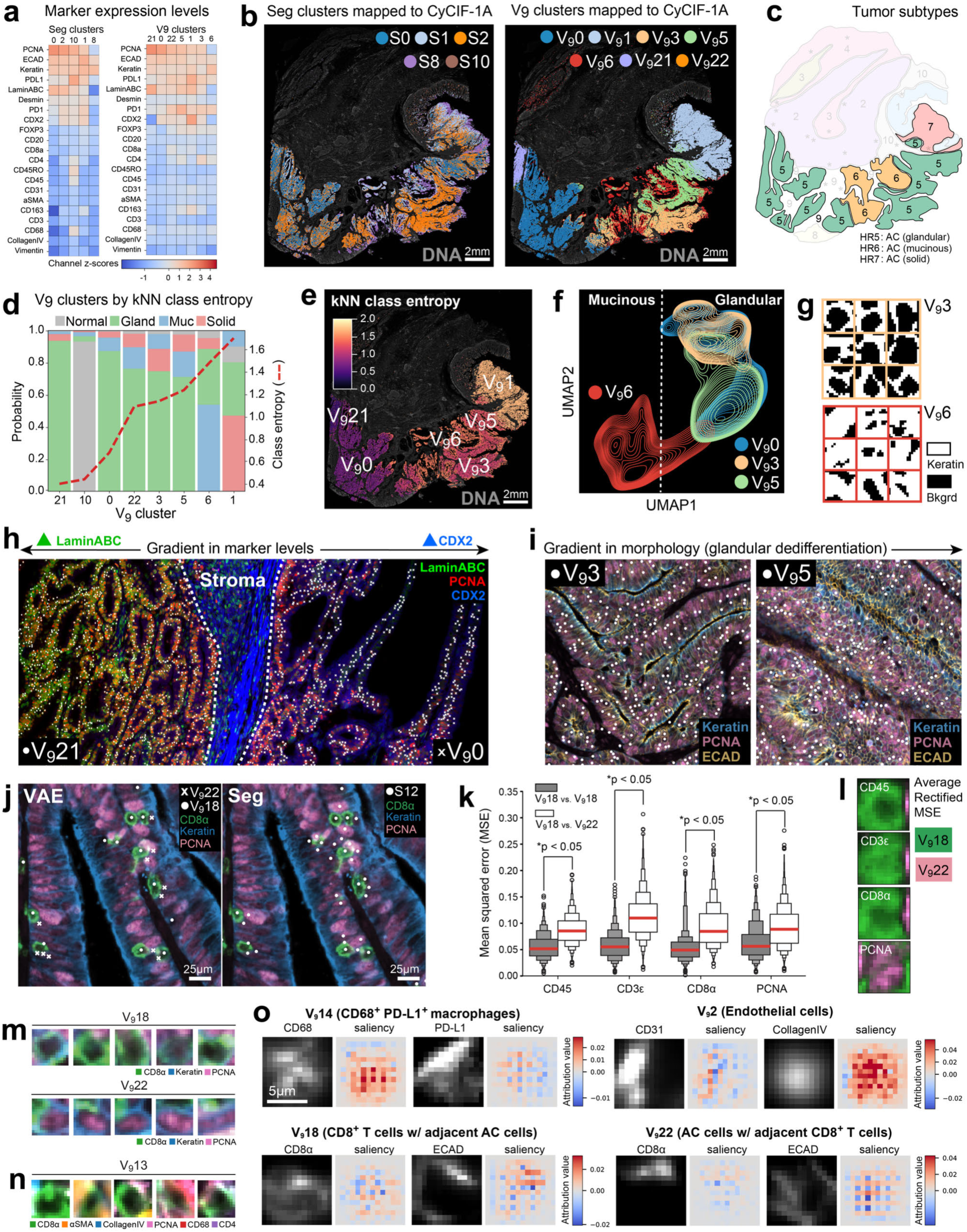
VAE image patch classification is morphology-aware and overcomes spatial crosstalk. **a**, z-score normalized protein expression profiles for tumor-associated segmentation (left) and VAE (right) Leiden clusters. **b**, Location of tumor-associated segmentation (left) and VAE (right) Leiden clusters in CyCIF-1A showing that V_9_ clusters are organized into spatially defined regions throughout the tumor domain. **c**, Histology map highlighting the locations of different AC subtypes observed in the H&E image: glandular (green), mucinous (orange), and solid (red; Fig. 1b). **d**, Stacked bar chart showing normalized mean posterior probability scores (left y-axis) for kNN classes corresponding to normal epithelium (gray) and glandular (green), mucinous (blue), and solid AC (red) among epithelial V_9_ clusters sorted from left to right in ascending rank order based on class entropy (right y-axis). **e**, Location of cells from tumor-associated VAE clusters from meta-cluster H in CyCIF-1A colored by kNN class entropy revealing a spatial gradient from lower entropy glandular AC at the left of the image to higher entropy mucinous and solid AC at the right. **f**, UMAP embedding showing kernel density estimates for binarized VAE image patch encodings of the keratin channel from tumor-associated V_9_ clusters 0, 3, 5, and 6. **g**, Example binary keratin image patches from V_9_ clusters annotated as glandular (V_9_3) and mucinous AC (V_9_6) showing their morphological differences. **h**, Field-of-view in CyCIF-1A at the interface between V_9_21 (left of stroma region) and V_9_0 (right of stroma region) showing that these two clusters differ in their expression of LaminABC (green), PCNA (red), and CDX2 (blue). **i**, Fields-of-view in CyCIF-1A of glandular AC clusters V_9_3 (left) and V_9_5 (right) as seen in channels keratin (blue), PCNA (pink), and ECAD (orange) revealing morphological difference in the density and organization of the glandular structures formed by these cells. **j**, Matched fields-of-view in CyCIF-1A as seen in the CD8⍺ (green), keratin (blue), and PCNA (pink) channels showing that VAE classification correctly distinguishes between tumor-infiltrating CD8^+^ T cells (V_9_18) and neighboring keratin^+^ epithelial cells (V_9_22; left image) that co-cluster in segmentation cluster S12 due to spatial crosstalk (right image). Note: due to random data subsampling, not all data points match between left and right images. **k**, Boxen plots showing mean squared error (MSE) distributions among pairs of V_9_18 (gray) image patches and between V_9_18 and V_9_22 (white) image patches for channels CD3ε, CD45, CD8α, and PCNA. Horizontal red lines indicate the median MSE at the center quantile of each distribution (p<0.05, two-sided Mann-Whitney U-test). **l**, Average rectified pixel-wise MSE for channels CD3ε, CD45, CD8α, and PCNA in V_9_18 (green) and V_9_22 (pink) image patches revealing CD45, CD3ε, and CD8⍺ signals appear at the center of V_9_18 patches and towards the periphery of V_9_22 patches; the opposite is true for PCNA. This indicates that reference cells in V_9_18 patches represent CD8^+^ T cells and those in V_9_22 patches represent PCNA^+^ tumor cells. **m**, Image patch galleries confirming that cluster V_9_18 cells represent CD8⍺^+^ (green) T cells (top row) with surrounding PCNA^+^ (pink) keratin^+^ (blue) tumor cells while V_9_22 represents the opposite scenario (bottom row). **n**, Image patch gallery showing that cells in V_9_13 represent non-tumor infiltrating CD8^+^ T cells neighbored by diverse cell types as evidenced by variable immunomarker expression at patch borders. **o**, Grayscale channels (left images) and corresponding concept saliency maps (right images) for image patches within the 95^th^ percentile of positive concept score distributions from four V_9_ clusters showing characteristic channels for each.

To determine the contribution of morphology to tumor-associated VAE clusters, we isolated the keratin channel of patches from meta-cluster H to and binarized them to remove intensity variation; we then trained a new VAE model. We found that cells with mucinous (V_9_6) and glandular (V_9_0, 3, and 5) morphologies occupied distinct regions in feature space and exhibited differences in keratin organization (**Fig. 3f,g**), although this was not true for solid AC (V_9_1). Thus, the shape of the keratin signal alone was sufficient to distinguish glandular from mucinous cells. This finding, combined with the marker intensity map for V_9_ clusters (**Fig. 3a**, right panel) as well as spatial gradients observed in cluster marker intensity (**Fig. 3h**) and tissue morphology (**Fig. 3i**), strongly suggest that the VAE network simultaneously captures intensity and morphology information when distinguishing tumor cell subtypes.

### Identifying factors driving VAE cluster membership

V_9_22 cancer cells were found throughout the tumor and were consistently surrounded by CD8^+^ T cells from cluster V_9_18 (**Fig. 3j**). The Jaccard index plot (**Extended Data Fig. 4c**) showed that these VAE clusters were combined in segmentation cluster S12. Thus, VAE-based learning correctly resolved CD8^+^ T cells and tumor cells affected by spatial crosstalk. To investigate how this occurred, V_9_18 and V_9_22 image patches were compared at a pixel level using mean squared error (MSE) analysis (**Fig. 3k**). This showed that variance between V_9_18 and V_9_22 patches was significantly greater than variance within V_9_18 patches for channels CD3ε, CD45, CD8α, and PCNA. Visualizing average MSE representations for each cluster showed that CD3ε, CD45, and CD8α signals were concentrated at the center of V_9_18 patches but the periphery of V_9_22 patches (**Fig. 3l**), while PCNA signals exhibited the opposite pattern. This result was confirmed by inspection of image patch galleries for V_9_18 and V_9_22 (**Fig. 3m**). CD8^+^ T cells were also found in V_9_13, but these cells were adjacent to different stromal and immune cell types in the lamina propria of normal epithelium (HR1), muscularis propria (HR2), and peritumoral stroma (HR9; **Fig. 3n**). We conclude that a VAE can distinguish adjacent cell types affected by spatial crosstalk in segmentation-based analysis by correctly learning different spatial patterns of protein expression and can further differentiate similar cell types (e.g., CD8^+^ T cells) based on tissue context.

To further study how patch features influenced cluster membership, we performed concept saliency analysis^22^—a method in interpretable AI that seeks to explain the features driving differences in ML classes. Archetypal image patch encodings or “concept vectors” (denoted, *Zc*) were first selected for each cluster based on maximum cosine similarity to the average encoding for all patches bearing the same cluster label. Individual patches were then scored for similarity to cluster archetypes by computing the dot product between the VAE encoding for a particular image patch (*Zi*) and the previously identified concept vector (*Sc = Zi ⋅ Zc*). This showed that image patches with and without a specific cluster label formed discrete concept score distributions (**Extended Data Fig. 7c**) and that the VAE identified groups of image patches with unique spatial features. We then generated concept saliency maps using the rectified gradient method^36^, which uses backpropagation to calculate pixel-wise contributions to a given concept score while minimizing noise (**Fig. 3o**, **Extended Data Fig. 7d** and **Supplementary Image Gallery 2**). In these maps, pixels with high attribution values corresponded to signals consistently identified with specific clusters. For example, in the case of CD68^+^ CD163^-^ PD-L1^+^ macrophages (cluster V_9_14), the CD68 and PD-L1 channels showed positive attribution values in regions of the patch with high fluorescence signals, whereas biologically unrelated signals from CD163 and CD8α had negative values (**Fig. 3o**). These results provide further evidence that intensity signals and spatial distributions learned in an unsupervised manner by a VAE are interpretable and biologically meaningful.

### Patch-based learning achieves greater precision relative to image segmentation

The Jaccard index plot showed that segmentation-based clusters were often subdivided by the VAE into multiple subclusters, suggesting higher precision in distinguishing cell types (**Extended Data Fig. 4c**). For example, within segmentation cluster S3 (annotated as macrophages) the VAE identified three phenotypically and spatially distinct cell types: (i) CD163^+^ CD68^+^ tumor-reactive macrophages in the peritumoral stroma (V_9_7, HR9), (ii) CD163^+^ tissue-resident macrophages in the muscularis propria (V_9_12, HR2), and (iii) CD68^+^ PD-L1^+^ tumor-associated macrophages (TAMs) infiltrating the tumor mass (V_9_14, HR5-7; **Fig. 4a**). S3 also contained multiple non-macrophage cells populations, including CD31⁺ vascular endothelial cells (V_9_2, 23.1% of S3 cells) in the serosa and subserosa (HR3 & 4) and vimentin⁺ fibroblasts (V_9_16, 11.7% of S3 cells) in the lamina propria of healthy mucosa (HR1). Co-clustering of such dissimilar cell types confounds methods for cell-type identification. A second example of cell types incorrectly grouped together in segmentation-based analysis but separated by the VAE was S13 (conventional CD4^+^ T cells, Tcons). Twenty-seven (27) percent of S13 cells were identified as CD163^+^ CD4^+^ tissue-resident macrophages (V_9_15)^37^ in HR1 and HR2 **(Fig. 4b-d**) with the remaining cells distributed across fibroblast and macrophage clusters (V_9_7, 14, 16). We reasoned that shared expression of CD4 between Tcons and this macrophage population caused them to co-cluster in S13. Thus, pixel-level learning enabled by a VAE was able to discriminate cell populations occupying distinct spatial contexts, lacking definitive immunomarker profiles (e.g., fibroblasts), or expressing atypical combinations of markers (CD163^+^ CD4^+^ macrophages). The identification of these CD4^+^ macrophages in CyCIF-1A highlights how unsupervised learning can “discover” unexpected cell types.

**Fig. 4.**
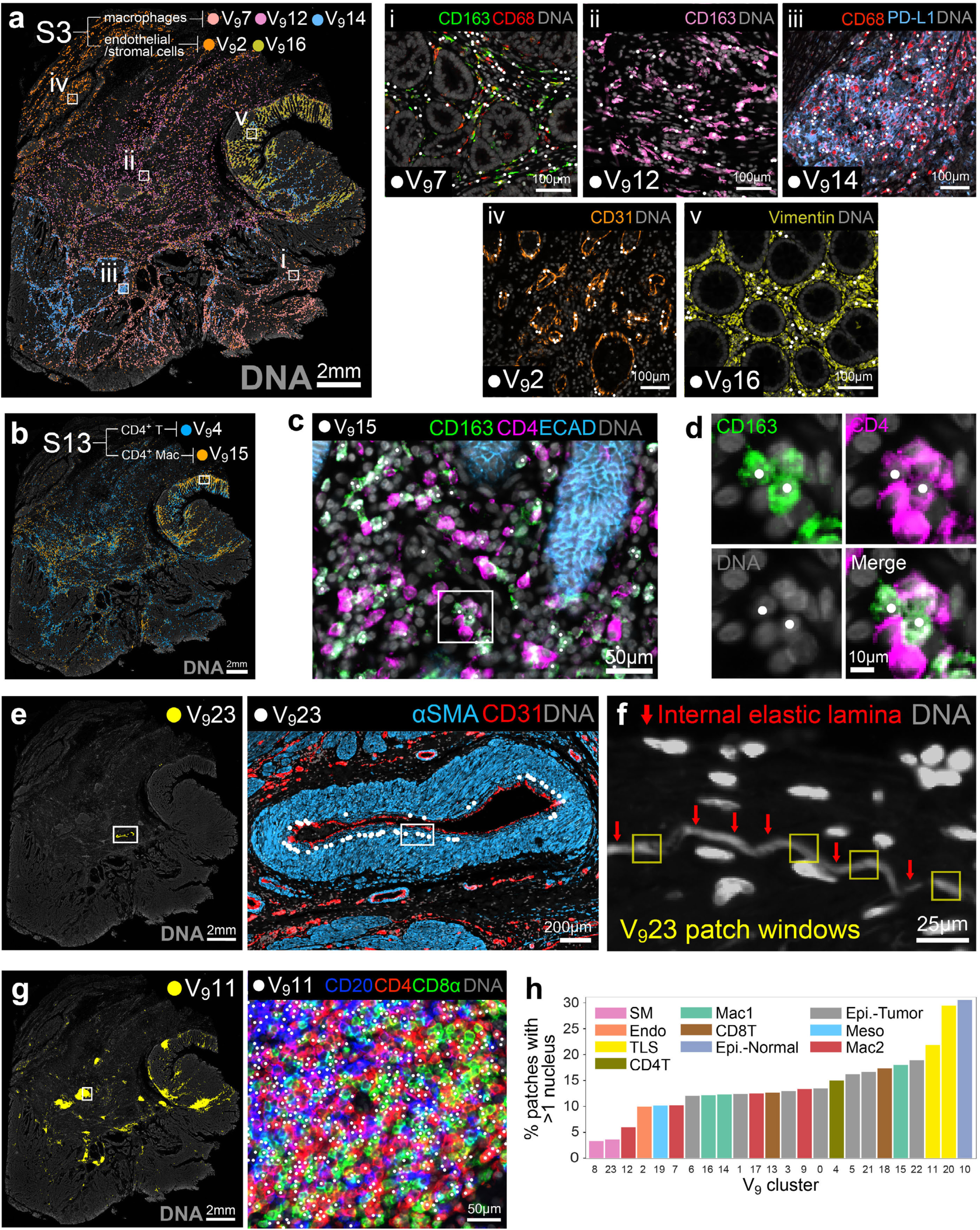
Strengths and limitations of patch-based learning for single-cell phenotyping. **a**, Left panel: Location of segmentation cluster S3 in CyCIF-1A colored by V_9_ cluster labels showing multiple subclusters identified by the VAE. Right panels: Fields-of-view corresponding to white boxes in left image showing example cells (white scatter points) identified by the VAE as macrophages (i-iii), fibroblasts (iv), and endothelial cells (v). **b**, Location of cells in segmentation cluster S13 in CyCIF-1A corresponding to V_9_4 (CD4^+^ T cells) and V_9_15 (CD4^+^ macrophages) in VAE analysis. **c**, Enlarged view of the region highlighted by the white box in panel b as seen in the CD163 (green), CD4 (magenta), and ECAD (blue) channels confirming that V_9_15 represents CD163^+^ CD4^+^ macrophages in the lamina propria of normal mucosal crypts (HR1); DNA (gray) channel shown for reference. **d**, Enlarged view of the region highlighted by the white box in panel c showing CD163^+^ CD4^+^ macrophages interacting with CD163^-^ CD4^+^ helper T cells; channels shown separately for clarity. **e**, Left panel: Location of V_9_23 cells in the muscularis propria (HR2) of CyCIF-1A. Right panel: Enlarged view of the region highlighted by the white box in left image as seen in the αSMA (blue) and CD31 (red) channels showing that V_9_23 represents smooth muscle cells proximal to the abluminal wall of an artery. **f**, Enlarged view of the region highlighted by the white box in the image at left as seen in the DNA (gray) channel highlighting the internal elastic lamina (red arrows), an autofluorescent acellular structure present in V_9_23 image patches (yellow boxes). **g**, Left image: Location of V_9_11 cells comprising lymphoid aggregates in CyCIF-1A. Right image: Enlarged view of the region highlighted by the white box in the image at left as seen in the CD20 (blue), CD4 (green), and CD8α (red) channels showing that V_9_11 comprises a mixed of CD20^+^ B cells, CD4^+^ T cells, and CD8^+^ T cells. **h**, Bar plot showing the percentage of V_9_ cluster cells with greater than one nucleus colored by meta-cluster labels from (Fig. 2i). Note that image patch clusters associated with lymphoid aggregates (V_9_11 and V_9_20, yellow bars) show a high proportion of cells with multiple nuclei, helping to explain why B and T cells comprising these structures do not form discrete clusters

V_9_23 presented another example of how unsupervised learning can identify recurrent spatial patterns not part of an *a priori* analysis plan. V_9_23 represented a rare population of ⍺SMA^+^ smooth muscle cells lining the basement membrane of an artery within the muscularis propria (0.1% of total cells, HR1; **Fig. 4e**). V_9_23 image patches were distinguished by the external elastic lamina—an acellular structure that provides mechanical support to the blood vessel wall and can be detected based on its strong and photostable autofluorescence^38^ (**Fig. 4f**). Acellular structures such as this are visible in images but lack nuclei and are therefore absent from segmentation-based spatial feature tables.

### Limitations of patch-based learning relative to image segmentation

In a few cases, VAE-based learning led to co-clustering of cell types that segmentation-based analysis correctly separated. For example, V_9_4 contained both Tcons (S13) and regulatory T cells (Tregs, S15; **Extended Data Fig. 8a,b** and **Supplementary Video 3**). In biplots of segmented cells, CD4 and FOXP3 formed discrete distributions corresponding to Tcons and Tregs, but this was not true in patch-based representations (**Extended Data Fig. 8c-e**). We speculate that this occurred because nuclear FOXP3 signals represent only a small fraction of total patch dimensionality. The localization of Tcons and Tregs to similar stromal niches failed to provide additional discriminatory information (**Extended Data Fig. 8f,g**). This highlights a limitation of unsupervised learning that could be addressed by increasing the weight of signals for specific lineage markers or cell compartments (e.g., the nucleus) during image patch encoding.

In both segmentation and VAE-based analysis, clusters corresponding to lymphoid aggregates (S11, V_9_11 and V_9_20, HR*; **Fig. 4g**) were heterogeneous with respect to cell type, containing both B and T cells. With segmentation, this arose from spatial crosstalk, but in VAE analysis it was due to image patches containing multiple nuclei due to close cell packing; more than 22% of patches from V_9_11 and V_9_20 contained >1 nucleus (**Fig. 4h**). Thus, while unsupervised VAE-based learning has greater sensitivity to morphological and contextual features it cannot separate whole cells that consistently co-appear in an image patch. A lack of prior knowledge about what is important but subtle differences (e.g., nuclear FOXP3) imposes additional limitations on unsupervised learning and underscores the value of performing pixel-level inference and traditional segmentation in parallel.

### VAE analysis identifies matched cell states within and across specimens

To explore the consistency of VAE-based learning across tissue sections and tumors, we trained a new network on 9x9µm image patches from three CyCIF images of CRC labeled with the same 21 antibodies (**Supplementary Table 2**). Tissues included (i) CyCIF-1A, to test technical reproducibility of the method; (ii) a serial section from the same tissue block located ∼20 µm along the Z-axis, to evaluate concordance across biological replicates (CyCIF-1B; **Fig. 5a**); and (iii) a tissue section from a different patient, to compare clusters across tumors (CyCIF-2; **Fig. 5b**). Leiden clustering of combined VAE image patch encodings from all three samples yielded 33 clusters (V_9M_0-32, with “V_9M_” denoting a VAE trained on 9x9µm patches from multiple tissues; **Fig. 5c**, **Supplementary Table 5** and **Supplementary Video 4**). Three clusters (V_9M_29, 31, and 32) were identified as microscopy artifacts (<0.3% of cells) that underwent no further analysis. The remaining clusters corresponded to the expected four primary cell lineages: keratin^+^ epithelial, ⍺SMA^+^ smooth muscle, vimentin^+^ stromal, and CD45^+^ immune cells (**Extended Data Fig. 9a,b** and **Supplementary Fig. 11**). V_9M_ clusters mapping to CyCIF-1A and 1B revealed strong concordance, with matched clusters occupying similar regions of tissue (**Fig. 5d** and **Supplementary Fig. 12**). Hierarchical clustering of V_9M_ clusters by protein expression profiles showed that 60% (18 of 30) of two-way dendrogram merges involved matched clusters from CyCIF-1A and 1B, and an additional 37% (11 clusters) involved three-way dendrogram merges between matched clusters from all three samples (**Fig. 5e**). Thus, 97% (60% + 37%) of V_9M_ clusters were matched between serial sections of the same tumor.

**Fig. 5.**
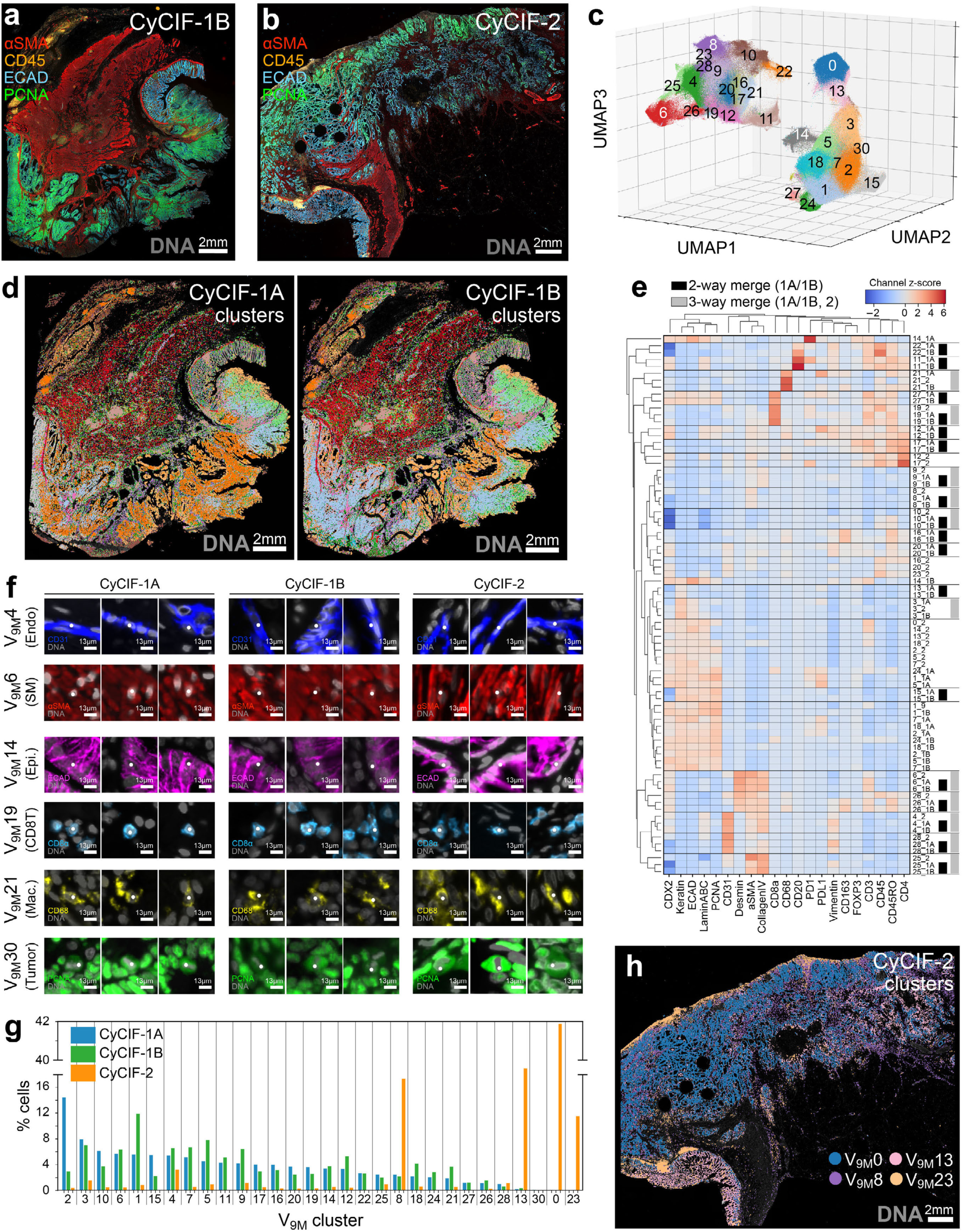
VAE analysis identifies matched cell states within and across patient specimens. **a**, CyCIF-1B: a second section from the same tissue block as CyCIF-1A (∼20µm away in the Z-axis) imaged for the same CyCIF antibodies listed in (**Supplementary Table 2**). **b**, CyCIF-2: a tissue section from a second CRC patient imaged for the same CyCIF antibodies as CyCIF-1A and 1B. **c**, 3D UMAP embedding of 9x9µm VAE image patch encodings from CyCIF-1A, CyCIF-1B, and CyCIF-2 colored by Leiden cluster. Clusters 29, 31, and 32 were identified as microscopy artifacts and are not labeled; see **Supplementary Video 4** for 360° plot rotation. **d**, Location of V_9M_ clusters in CyCIF-1A (left) and 1B (right) revealing color-matched clusters in overlapping locations in serial sections of this tissue; DNA channel (gray) shown for reference. **e**, Hierarchical clustering of z-score-normalized median marker intensities of tissue-specific V_9M_ clusters showing that many share similar protein expression profiles. Black vertical bars indicate two-way dendrogram merges between CyCIF-1A and 1B. Gray vertical bars indicate three-way merges between all three specimens. Data are filtered to remove clusters associated with microscopy artifacts (V_9M_29, 31, and 32) and clusters representing <0.1% of their respective tissue. **f**, V_9M_ clusters identified in all three specimens as seen in their characteristic marker channels with reference cells highlighted by white scatter points; DNA (gray) shown for reference. **g**, Grouped bar chart showing the percentage of cells in each specimen accounted for by each V_9M_ cluster sorted in descending rank-order according to cluster frequency in CyCIF-1A. Clusters identified as microscopy artifacts (V_9M_29, 31, and 32) are not shown. **h**, Location of the four most abundant V_9M_ clusters in CyCIF-2; DNA channel (gray) shown for reference.

Clusters common to all three images included: endothelial cells (V_9M_4), smooth muscle cells (V_9M_6), crypt-forming epithelial cells (V_9M_14), CD8^+^ T cells (V_9M_19), CD68^+^ macrophages (V_9M_21) and mitotic tumor cells (V_9M_30; **Fig. 5f**). The latter were identified based on their condensed nuclear chromatin (a feature of pro-metaphase and metaphase cells), which was visible in the DNA channel and as a silhouette in the PCNA channel (**Extended Data Fig. 9c**). Due to their scarcity (<0.08% of cells in each specimen), these cells were missed in single-tissue analysis but were detected in multi-tissue analysis simply because there was more data. Four clusters (V_9M_0, 8, 13, and 23) were unique to CyCIF-2, together accounting for 85.5% of cells (**Fig. 5g,h** and **Extended Data Fig. 9d** and **Supplementary Video 5**). The largest clusters, V_9M_0 (41.2%) and V_9M_13 (17.6%), corresponded to PCNA^high^ and PCNA^low/neg^ tumor cells, respectively (**Extended Data Fig. 9e**). Both exhibited abnormal CD3ε labeling, likely due to expression of cross-reactive epitopes, and explaining why CyCIF-2 tumor cells did not co-cluster with those in CyCIF-1 (**Extended Data Fig. 9f,g**). While V_9M_8 cells (fibroblasts) were found in all three samples, they were 7-fold more abundant in CyCIF-2 due to a higher stroma-to-tumor ratio (**Fig. 5g** and **Extended Data Fig. 9h**). V_9M_23 represented CD45^+^ cells specific to CyCIF-2 that lacked expression of other immune lineage markers (**Extended Data Fig. 9i**). However, the same tumor-infiltrating CD8^+^ T cells (V_9_18) and surrounding keratin^+^ tumor cells (V_9_22) in CyCIF-1A were also identified in the other two images (V_9M_24 and V_9M_27; **Extended Data Fig. 9j**), emphasizing the ability of a VAE to reproducibly overcome spatial crosstalk. These results show that VAE-based learning reliably identified similar cell types across tissues, including rare populations.

### VAE analysis of multicellular image patches identifies recurrent tissue motifs

VAE networks can be trained on image patches of arbitrary dimensions. To classify CRC according to multi-cellular architecture, we trained a VAE model on 30x30µm image patches containing an average of 5 nuclei per patch (**Fig. 6a**). Although patches were centered on reference cells for indexing purposes, Gaussian vignetting was omitted to ensure that all pixels were visible to the network at full intensity. At convergence, the model achieved a validation loss of 1.4x10^-3^, with VAE-generated representations closely resembling input image patches (**Fig. 6b**). As in V_9_ analysis, the distribution of concept scores for each V_30_ cluster were bimodal, indicating that clusters with distinct features had been identified (**Extended Data Fig. 10a**). Leiden clustering applied to 30x30µm image patch encodings generated 12 clusters (V_30_0-11) including one microscopy artifact (V_30_11, 0.1% of cells; **Fig. 6c**, **Supplementary Table 6**, **Supplementary Video 6**, **Supplementary Image Gallery 3**). The spatial distributions of V_30_ clusters corresponded to histologic regions in the H&E image, including glandular (V_30_2/8, HR5), mucinous (V_30_4, HR6), and solid AC (V_30_0, HR7); the muscularis propria (V_30_5/6; HR2); normal mucosa (V_30_7, HR1); and serosal mesothelium (V_30_10, HR3). In addition, V_30_9 correctly captured all 17 lymphoid aggregates annotated by our histopathologists in the H&E image plus several smaller ones (V_30_9, HR*; **Fig. 6d**).

**Fig. 6.**
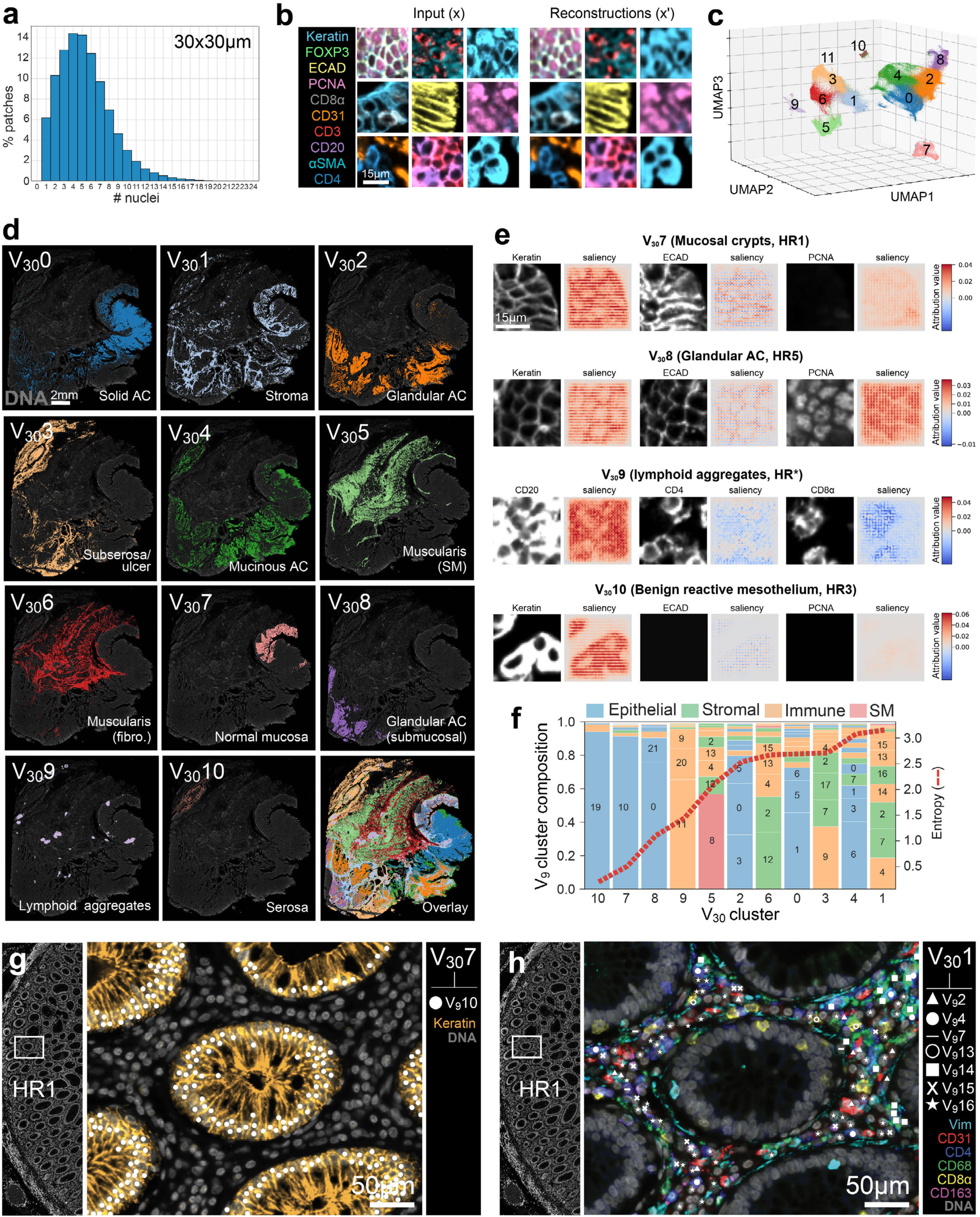
VAE classification of multi-cellular image patches identifies recurrent tissue motifs. **a**, Histogram showing the percentage of 30x30µm image patches containing different numbers of cell nuclei (median = 5, Q1 = 3, Q3 = 7, IQR = 4). **b**, Left: Example 30x30µm multi-channel image patches extracted from CyCIF-1A and used as input into the VAE encoding network. Right: Learned reconstructions of input images shown at left. **c**, 3D UMAP embedding of 30x30µm VAE image patch encodings from CyCIF-1A colored by Leiden cluster; see **Supplementary Video 6** for 360° plot rotation. **d**, Locations of V_30_ clusters in CyCIF-1A corresponding to histological regions observed in the H&E image (Fig. 1b**,c**); DNA channel (gray) shown for reference. **e**, Grayscale channels (left images) and corresponding concept saliency maps (right images) for four V_30_ clusters showing characteristic channels. **f**, Stacked bar chart showing the composition of reference cells at the center of V_30_ cluster image patches (x-axis) according to their V_9_ cluster labels (left y-axis) and their cell type heterogeneity as measured by information entropy (right y-axis, dashed red curve). **g**, Location of reference cells in V_30_7 (a low-entropy cluster) from CyCIF-1A composed of V_9_10 cells as seen in the keratin (orange) channel, indicating that this V_30_ cluster represents healthy mucosal crypts consisting of normal epithelial cells (V_9_10). **h**, Location of reference cells in V_30_1 (a high-entropy cluster) from CyCIF-1A composed of multiple V_9_ cell types, including fibroblasts (vimentin, cyan), endothelial cells (CD31, red), helper T cells (CD4, blue), M1-like macrophages (CD68, green), cytotoxic T cells (CD8⍺, yellow), and M2-like macrophages (CD163, magenta) indicating that this cluster captures cellularly heterogeneous stromal tissue as a recurrent tissue motif.

Because V_30_ clusters contained more pixels and were not vignetted, concept saliency maps had higher resolution, with high-attribution pixels forming clear multi-cellular spatial patterns (**Fig. 6e**, **Extended Data Fig. 10b**, and **Supplementary Image Gallery 4**). For example, keratin saliency maps reproduced the characteristic morphologies of epithelial cells in normal mucosal crypts (V_30_7) and glandular AC (V_30_8), while maps for lymphoid aggregates (V_30_9) highlighted strong contributions from CD20^+^ B cells and CD4^+^ helper T cells, suggesting their importance in defining this cluster concept. In contrast, CD8^+^ T cells contributed negatively to the lymphoid aggregate cluster concept, consistent with the observation that CD8^+^ T cells were less abundant in these structures than B and helper T cells.

To better understand the cell type composition of V_30_ clusters, we used information entropy computed against V_9_ cluster labels. Entropy scores (which had possible values between 0 to 4.6 bits based on 24 possible V_9_ labels) ranged from 0.38 bits for V_30_10 to 3.18 bits for V_30_1 (**Fig. 6f**). At least 75% of cells in low-entropy clusters V_30_7, 8 and 10 corresponded to a single V_9_ cluster label, showing that these structures were primarily composed of a single cell type. For example, 92% of cells in V_30_7 were normal crypt-forming epithelial cells (V_9_10; **Fig. 6g**). However, cells in V_30_1 (a high-entropy cluster) comprised ten different V_9_ clusters, including CD4^+^ T cells (V_9_4, 19.0%), CD68^+^ CD163^+^ macrophages (V_9_7, 17.3%), CD31^+^ endothelial cells (V_9_2, 15.8%), CD68^+^ PD-L1^+^ macrophages (V_9_14, 10.9%), vimentin^+^ fibroblasts (V_9_16, 10.7%), CD8^+^ T cells (V_9_13, 9.5%), and CD163^+^ CD4^+^ macrophages (V_9_15, 6.3%; **Fig. 6h**). This diversity of cell types reflects the role of the colonic stroma in structural support, nutrient exchange, and wound healing mediated by intermixed fibroblasts, immune, and other cell types. Notably, both the lamina propria of healthy mucosal crypts (HR1) and the peritumoral stroma surrounding cancer cells (HR9) were both part of V_30_1, suggesting that these compartments share a common set of constituent cell types. We conclude that a VAE trained on multicellular image patches can correctly identify morphologically distinct arrangements of single cell types as well as complex cellular communities comprising multiple lineages in normal and desmoplastic stromal tissue.

## DISCUSSION

We have described a framework (MORPHAEUS, Morphology Oriented Research in Pathology Harnessing Autoencoder Unsupervised Strategies) that combines the strengths of unsupervised pixel-level ML and traditional image segmentation to support tissue classification across spatial scales, from single cells to tissue neighborhoods. The resulting data fit into current single-cell analysis pipelines where adjacent cells that cannot be separated are treated as a community. We show that VAEs can extract information from images corresponding to marker intensity, cell morphology, and local neighborhood in a manner analogous to human experts, providing deeper insight into tissue composition and organization than segmentation-based analysis alone while addressing spatial crosstalk.

Generative models such as VAEs, adversarial autoencoders, deep convolutional generative adversarial networks, and combinations thereof require no ground truth annotations, making them well-suited to discovery of unanticipated features. In our work VAEs detected (i) an unexpected population of CD163^+^ CD4^+^ tissue-resident macrophages^37^ (V_9_15) missed by segmentation-based clustering, (ii) mitotic tumor cells (V_9M_30) with distinctive DNA morphology, and (iii) cells adjacent to acellular structures such as the arterial elastic lamina (V_9_23). Because VAEs exploit spatial patterns in marker levels to learn neighborhood features, they represent a novel approach to overcoming spatial crosstalk in segmentation-based methods. VAEs accomplish this by identifying multi-cellular cellular niches such as CD8^+^ T cells in stromal (V_9_13) versus tumor tissue (V_9_18).

Moreover, whereas segmentation assumes all cells are separable and spatial relationships are inferred using spatial statistics, training VAEs on patches of different sizes enables learning of both cell-level and neighborhood features simultaneously. This represents a conceptually different approach to spatial profiling in which we accept that it is not possible, or even desirable, to singulate every cell and then attempt to reassemble neighborhoods from single cell data. Instead, VAE-based approaches leave neighborhoods intact depending on patch size: 9μm patches captured single cells or closely associated pairs (e.g. T cells and tumor cells), while 30μm patches captured multi-cellular neighborhoods.

Despite the strengths of unsupervised VAE-based learning, it can miss subtle nuclear features such as the difference between Tcons and Tregs based on expression of the nuclear transcription factor FOXP3.^39^ Segmentation-based methods may be superior in this case because they implicitly increase the saliency of nuclear signals due to the use of nuclear segmentation masks. However, the same effect may be achievable via feature engineering during VAE image patch encoding.^40^ Both segmentation and VAE models also fail to separate B and T cells in dense lymphoid aggregates. A potential solution is a multi-pass learning in which a smaller patch size is used for cell-dense regions of tissue such as lymphoid aggregates with larger patches used to infer cells in the remainder of tissue. However, such approaches will likely be most effective using higher resolution data^41^ that can more effectively discern between cell types. In some cases it may be reasonable to view T and B cell pairs in densely-packed lymphoid aggregates as a recurrent tissue motif rather than sets of fully discriminable cells. Thus, we suggest performing traditional segmentation and patch-based learning in parallel, and using the comparison to optimize patch size, adjust vignetting, and tune clustering parameters.

## SUPPLEMENTARY INFORMATION

### Supplementary Note 1 – Additional information on segmentation methods and cell type calling

Highly multiplex tissue imaging is a rapidly evolving technology whose analysis often involves defining boundaries around individual cells through the process of image segmentation. In the case of whole-slide tissue imaging, this involves the acquisition of tens to hundreds of image tiles stitched and registered together using algorithms such as ASHLAR^42^ that form multi-channel mosaic images stored in metadata-rich open file formats like OME-TIFF^43^ (level 2 data according to MITI guidelines^2^, **Supplementary Fig. 1a**).

Semantic segmentation algorithms, such as the popular U-Net^44,45^ architecture and its derivatives such as UnMICST^46^, TissueNet^47^, and Cellpose^7^ are then applied to images of cell nuclei labeled with DNA intercalating dyes (e.g., Hoechst or DAPI) to identify boundaries between cells and generate cell segmentation masks—reference images in which pixels are labeled according to the cell they belong to (MITI level 3 data, **Supplementary Fig. 1b**).

Mean immunomarker signals for individual cells are then programmatically computed and organized into a tabular data format like CSV (level 4 data according to MITI guidelines, **Supplementary Fig. 1c**). The derived tables are often augmented by measurements of the segmentation instances themselves, which serve as proxies for nuclear morphology (e.g., area, eccentricity, etc.). However, the accuracy of these measurements is conditional on segmentation quality. While standard pipelines such as MCMICRO^28^ have been developed to automate the aforementioned image processing tasks, no consensus exists on how best to classify cell phenotypes in tissue-derived, single-cell data.

A common method for segmentation-based cell state inference involves manual thresholding per-cell signal intensities (i.e., hard gating) performed using univariate histograms or bivariate scatter plots (**Supplementary Fig. 1d**). However, as positive and negative cell populations often overlap, such methods can be sensitive to even small changes in gate placement, leading to ambiguous annotation and potentially changing biological interpretation. Methods based on Gaussian mixture models (GMMs) and naïve Bayes classifiers such as those implemented by the marker-scoring functions in CELESTA^48^ and MCMICRO^28^, respectively, represent “soft” gating methods. Such methods have the advantage of assigning class-conditional probability weights to individual cells as an alternative to strict thresholding (**Supplementary Fig. 1e**). Nevertheless, these methods also require manual parameter tuning and are sensitive to the same limitations imposed by segmentation-based analysis.

Unsupervised clustering methods (e.g., Louvain^49^ or Leiden^31^ community detection, k-means^50^, SOMs^51^ (self-organizing maps), and HDBSCAN^52^ (hierarchical density-based spatial clustering of applications with noise)) are an alternative to gate-based approaches for cell state inference. In these methods, high-dimensional single-cell data are projected onto lower dimensional embeddings (typically 2D or 3D) using data dimensionality reduction algorithms such as t-SNE (t-distributed stochastic neighbor embedding)^53^ or UMAP (uniform manifold approximation and projection)^33^ before or after clustering (**Supplementary Fig. 1f**). Regardless of the method used for cell phenotyping, protein expression profiles for inferred cell populations are often visualized as clustered heatmaps that can be interpreted by domain experts according to prior knowledge of cellular protein expression profiles (**Supplementary Fig. 1g**).

### Supplementary Note 2 – Analysis of Errors in Segmentation Cluster S5

Based on a heatmap of marker expression, segmentation cluster S5 appeared to express markers associated with multiple cell types, including smooth muscle cells (αSMA, collagen IV), macrophages (CD68, CD163), endothelial cells (CD31), mesenchymal cells (vimentin), and CD8^+^ T cells (CD45, CD3ε, CD8α, CD45RO; **Fig. 1e**). Unexpectedly, cells in this cluster gave rise to multiple subclusters in UMAP space, which we termed S5.1, S5.2, and S5.3 (**Supplementary Fig. 5a**). Viewing these cells in CyCIF-1A revealed that they corresponded to phenotypically similar CD8^+^ T cells occupying distinct histological regions that also differed with respect to their protein expression profiles (**Supplementary Fig. 5d,c**). Closer inspection showed cells in S5.1 were influenced by spatial crosstalk with CD68^+^ PD-L1^+^ macrophages located in the peritumoral stroma (HR9) and stromal compartments throughout the muscularis propria (HR2; **Supplementary Fig. 5d**), while cells in S5.2 were affected by crosstalk with Desmin^+^ smooth muscle cells in the muscularis (**Supplementary Fig. 5e**). S5.3 cells, which were located in the serosa (HR3) and subserosa (HR4), did not appear to be impacted by spatial crosstalk. Instead, they were distinguished by high CD45 expression, a feature we had previously observed in immune cells occupying these tissue compartments (**Supplementary Fig. 5f**).

Although all S5 cells were grouped under the same label by the Leiden clustering algorithm, they formed three distinct subpopulations in 3D UMAP space. This observation led us to hypothesize that the effects of spatial crosstalk are more apparent in low-dimensional embeddings of high-plex data due to data compression. This idea aligns with previous findings that distances and relationships between data points are not reliably preserved during dimensionality reduction^54^. To test this hypothesis, we performed density-based clustering using HDBSCAN^52^ on both 2D and 3D UMAP embeddings of cells from CyCIF-1A. Supporting our hypothesis, we found that cells grouped into a single cluster in 3D UMAP space were often redistributed into multiple clusters in a 2D embedding. For example, a subset of tumor cells originally grouped within cluster 17 in a 3D embedding (**Supplementary Fig. 6a-c**) were delocalized in a 2D embedding of the same data (**Supplementary Fig. 6d**). These delocalized cells were found to reside at the tumor-stromal interface and were influenced by spatial crosstalk from collagen IV signals originating in neighboring stromal cells (**Supplementary Fig. 6e,f**). Examples like this help explain the delocalization of three CD8^+^ T cell subsets comprising S5 (S5.1, S5.2, and S5.3) in UMAP feature space. They also underscore the creation of artifactual clusters by spatial crosstalk.

### Supplementary Note 3 – Testing VAE-based analysis on different types of high-plex tissue images

To assess the ability of the method to generalize across different imaging platforms and tissue types, independent VAE models were trained on Lunaphore COMET and Akoya PhenoCycler (CODEX) data. The Lunaphore image (referred to as Lunaphore-1) comprised a section of human non-small cell lung cancer (NSCLC) stained for 27 markers and acquired at a nominal resolution of 0.28 µm/pixel (compared to 0.65 µm/pixel for CyCIF-1A). We found that image patches in this dataset cropped at 5x5µm (compared to 9x9µm for CyCIF-1A) best approximated single cells. Leiden community detection identified 19 VAE image patch clusters (V_5_0-V_5_18) well separated in UMAP space (**Extended Data Fig. 6a**). Clustering protein expression profiles revealed biologically recognizable populations, including IDO-1⁺ and IDO-1⁻ lung cancer cells (V_5_0 and V_5_11, respectively), ⍺SMA^lo^ and ⍺SMA^hi^ vasculature (V_5_1 and V_5_2, respectively), FOXP3⁺ Tregs (V_5_12), CD66b⁺ granulocytes (V_5_16), and podoplanin⁺ lymphatic vessels (V_5_17; **Extended Data Fig. 6b**). Mapping clusters onto the original image confirmed their identities based on characteristic morphologies and spatial distributions (**Extended Data Fig. 6c,d** and **Supplementary Fig. 9**). For example, clusters V_5_1 and V_5_2 corresponded to well-structured, pre-existing (V_5_1) and disorganized, neo-angiogeneic (V_5_2) blood vessels.

The PhenoCycler image (referred to as CODEX-1) comprised a tissue microarray (TMA) core of human tongue squamous cell carcinoma stained for 29 markers and imaged at a nominal resolution of 0.5 µm/pixel. Unlike Lunaphore and CyCIF images, which were stored at 16-bit depth, these data were stored at 8-bit depth. Single cells in this image were approximated using 11x11µm image patches. Despite the lower bit depth and relatively small number of cells in this image (∼10K), the Leiden algorithm identified 19 VAE image patch clusters (V_11_0-V_11_18) that were also well separated in UMAP space (**Extended Data Fig. 6e**). Multiple tumor cell phenotypes were identified in this sample based on differences in marker intensity and morphology, including CD44⁺ stem-like (V_11_0) and CK17⁺ basal-like (V_11_5) carcinoma cells. Other identified cell types included cancer-associated fibroblasts (CAFs, V_11_4), normoxic and hypoxic macrophages (V_11_11 and V_11_18, respectively), and memory CD4⁺ T cells (V_11_16; **Extended Data Fig. 6f**). These clusters were also validated by referencing the primary image (**Extended Data Fig. 6g,h** and **Supplementary Fig. 10**). Taken together, these additional analyses demonstrated that, despite differences in image bit depth, nominal resolution, and cell number, VAE-based cell state inference generalizes effectively across diverse spatial proteomics platforms.

## FIGURES and LEGENDS

**Supplementary Table 1.**
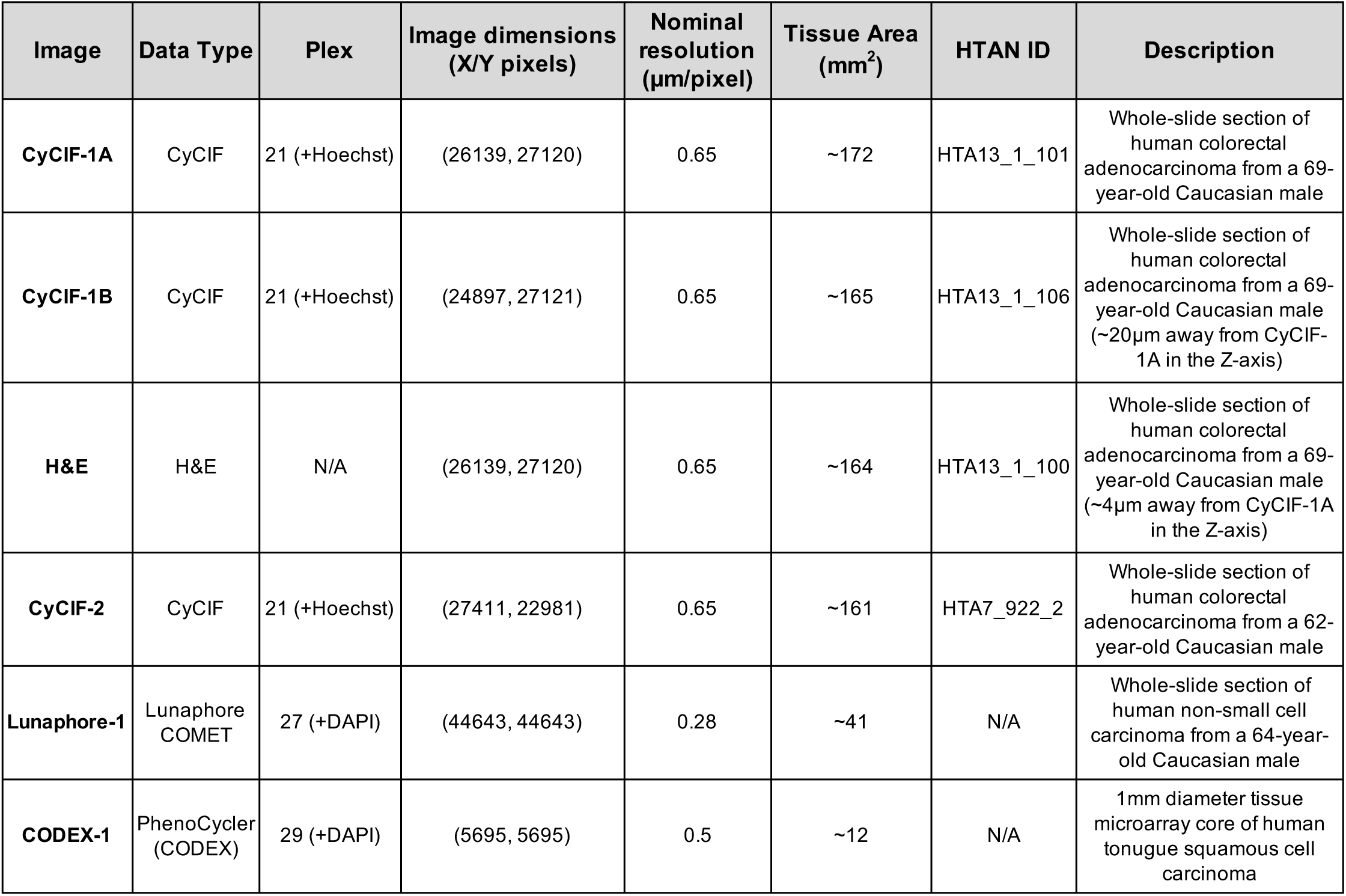
Metadata for biospecimens used in this study. Primary imaging data for the first four samples can be found at the HTAN Data Portal (https://data.humantumoratlas.org/explore) by referencing their corresponding HTAN IDs. Lunaphore and CODEX images are specific to this study and can be accessed upon request from the authors. Additional information on CyCIF antibodies used to label samples CyCIF-1A, CyCIF-1B, and CyCIF-2 can be found in the STAR Methods section of Lin et al. *Cell* 2023 (DOI: 10.1016/j.cell.2022.12.028).

| Image | Data Type | Plex | Image dimensions (X/Y pixels) | Nominal resolution ($\mu\text{m}/\text{pixel}$) | Tissue Area ($\text{mm}^2$) | HTAN ID | Description |
| --- | --- | --- | --- | --- | --- | --- | --- |
| **CyCIF-1A** | CyCIF | 21 (+Hoechst) | (26139, 27120) | 0.65 | ~172 | HTA13_1_101 | Whole-slide section of human colorectal adenocarcinoma from a 69-year-old Caucasian male |
| **CyCIF-1B** | CyCIF | 21 (+Hoechst) | (24897, 27121) | 0.65 | ~165 | HTA13_1_106 | Whole-slide section of human colorectal adenocarcinoma from a 69-year-old Caucasian male (~20$\mu\text{m}$ away from CyCIF-1A in the Z-axis) |
| **H&E** | H&E | N/A | (26139, 27120) | 0.65 | ~164 | HTA13_1_100 | Whole-slide section of human colorectal adenocarcinoma from a 69-year-old Caucasian male (~4$\mu\text{m}$ away from CyCIF-1A in the Z-axis) |
| **CyCIF-2** | CyCIF | 21 (+Hoechst) | (27411, 22981) | 0.65 | ~161 | HTA7_922_2 | Whole-slide section of human colorectal adenocarcinoma from a 62-year-old Caucasian male |
| **Lunaphore-1** | Lunaphore COMET | 27 (+DAPI) | (44643, 44643) | 0.28 | ~41 | N/A | Whole-slide section of human non-small cell carcinoma from a 64-year-old Caucasian male |
| **CODEX-1** | PhenoCycler (CODEX) | 29 (+DAPI) | (5695, 5695) | 0.5 | ~12 | N/A | 1 mm diameter tissue microarray core of human tongue squamous cell carcinoma |

**Supplementary Table 2.** Metadata for CyCIF antibodies. The “Segmentation Object” column indicates the cellular compartment used to extract marker signals for that channel. The column labeled “Analyzed” indicates whether a channel was evaluated in the current study. Unused channels represent secondary antibodies (e.g., channel 2, AF488) used to block non-specific antibody binding in downstream CyCIF cycles, repeated DNA images used for cross-cycle image registration, or markers that failed initial quality control.

| Channel | Marker | Segmentation Object | Analyzed |
| --- | --- | --- | --- |
| 1 | Hoechst-cycle1 | nucleus | yes |
| 2 | AF488 | N/A | no |
| 3 | AF555 | N/A | no |
| 4 | AF647 | N/A | no |
| 5 | Hoechst-cycle2 | nucleus | no |
| 6 | A488 | N/A | no |
| 7 | A555 | N/A | no |
| 8 | A647 | N/A | no |
| 9 | Hoechst-cycle3 | nucleus | no |
| 10 | CD3 | membrane | yes |
| 11 | NaKATPase | membrane | no |
| 12 | CD45RO | membrane | yes |
| 13 | Hoechst-cycle4 | nucleus | no |
| 14 | Ki67 | nucleus | no |
| 15 | Keratin | cytoplasm | yes |
| 16 | αSMA | cytoplasm | yes |
| 17 | Hoechst-cycle5 | nucleus | no |
| 18 | CD4 | membrane | yes |
| 19 | CD45 | membrane | yes |
| 20 | PD1 | membrane | yes |
| 21 | Hoechst-cycle6 | nucleus | no |
| 22 | CD20 | membrane | yes |
| 23 | CD68 | cytoplasm | yes |
| 24 | CD8a | membrane | yes |
| 25 | Hoechst-cycle7 | nucleus | no |
| 26 | CD163 | cytoplasm | yes |
| 27 | FOXP3 | nucleus | yes |
| 28 | PD-L1 | membrane | yes |
| 29 | Hoechst-cycle8 | nucleus | no |
| 30 | ECAD | cytoplasm | yes |
| 31 | Vimentin | cytoplasm | yes |
| 32 | CDX2 | cytoplasm | yes |
| 33 | Hoechst-cycle9 | nucleus | no |
| 34 | LaminABC | nucleus | yes |
| 35 | Desmin | cytoplasm | yes |
| 36 | CD31 | nucleus | yes |
| 37 | Hoechst-cycle10 | nucleus | no |
| 38 | PCNA | nucleus | yes |
| 39 | Ki67 | nucleus | no |
| 40 | CollagenIV | cytoplasm | yes |

**Extended Data Fig. 1.**
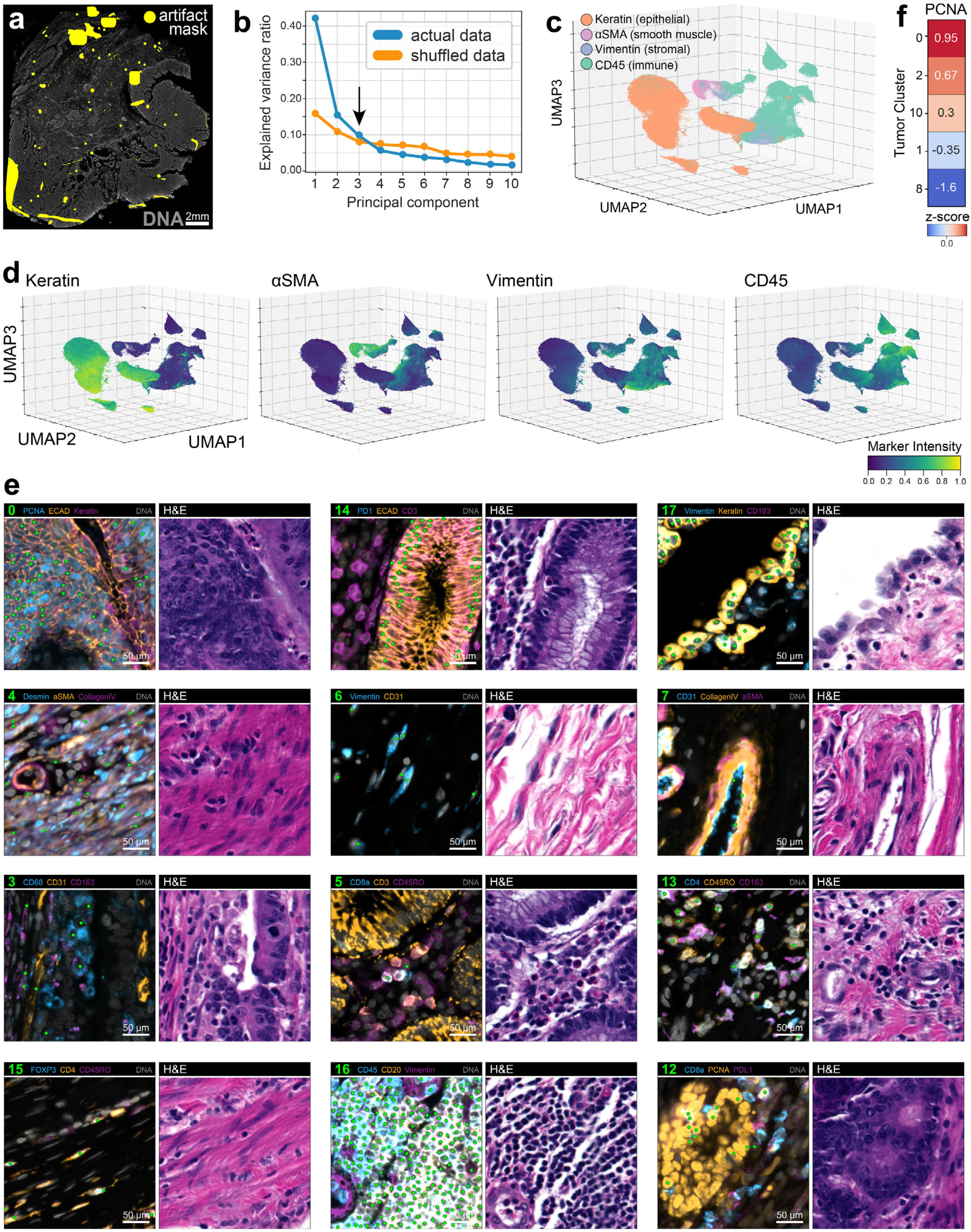
Segmentation-based cell phenotyping of human CRC. **a**, Artifact masks (yellow ROIs) highlighting regions in CyCIF-1A affected by microscopy artifacts (e.g., tissue folds, antibody aggregates, illumination aberrations, etc.) censored from further analysis. **b**, Horn’s parallel analysis showing the fraction of variance in post-QC single-cell data from CyCIF-1A explained by the first 10 principal components (blue curve) relative to the same data after randomization (orange curve). Black arrow highlights the point of intersection between the two curves, indicating that the first three principal components capture non-random variation in the dataset. **c**, UMAP embedding of segmented cells from CyCIF-1A colored by major cell lineage markers: keratin (epithelial, orange), αSMA (smooth muscle, pink), vimentin (stromal, blue), and CD45 (immune, green). **d**, UMAP embedding of segmented cells from CyCIF-1A colored by mean marker intensity for major cell lineage markers keratin, αSMA, vimentin, and CD45. See **Supplementary Fig. 2** for colormaps of all 21 immunomarkers. **e**, Paired CyCIF (left) and H&E (right) fields-of-view from CyCIF-1A showing the location of random examples of cells from various clusters (green scatter points in CyCIF images). See **Supplementary Fig. 3** for views of all 18 clusters. **f**, Heatmap of z-score-normalized mean PCNA intensities among tumor-associated clusters.

**Supplementary Fig. 1.**
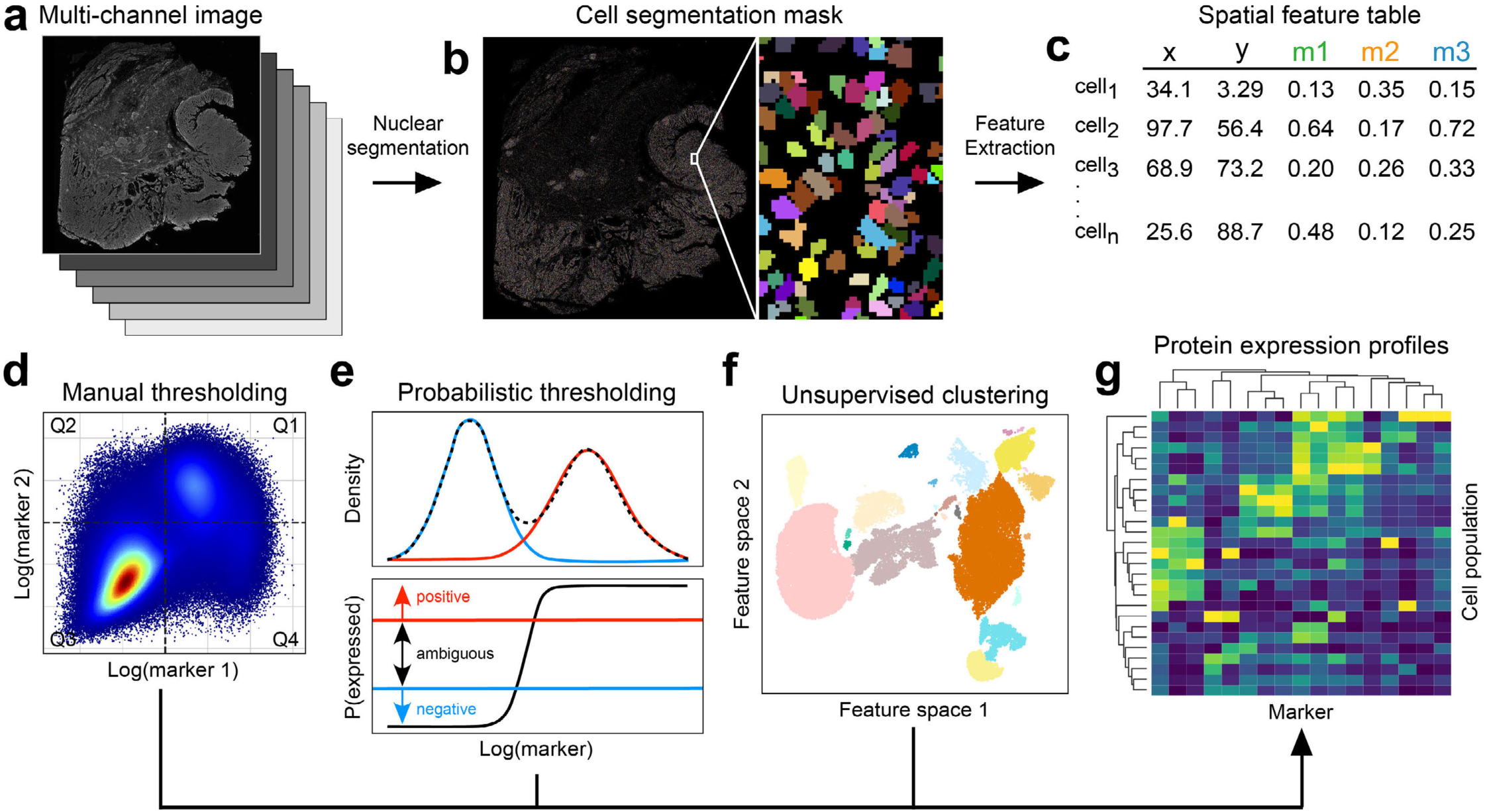
Workflow for segmentation-based cell phenotyping of multiplex tissue images. **a**, Multi-channel OME-TIFF/TIFF file. **b**, Segmentation mask generated by semantic segmentation of counterstained cell nuclei. **c**, Spatial feature table specifying X/Y spatial coordinates and mean marker intensities (e.g., m1-m3) for segmented cells (rows). **d**, Quadrant gating of single-cell data according to two markers colored by cell density. **e**, Two-mode Gaussian mixture model for assigning continuous probability weights for single marker expression values of single-cell data. Probability of immunoreactivity is computed as the Bayesian posterior probability (y-axis, bottom plot) at a given channel intensity (x-axis). Cells with posterior probabilities above the horizontal red line are positive for the marker, those beneath the horizontal blue line are negative. Those with posterior probabilities between the red and blue lines are deemed ambiguous for the marker in this example. **f**, Unsupervised clustering of high-dimensional single-cell data embedded in a 2D feature space (e.g., UMAP or t-SNE). **g**, Clustered heatmap of mean marker intensities (columns) for different cell populations (rows) identified by any of the methods shown in panels d-f.

**Supplementary Table 3.**
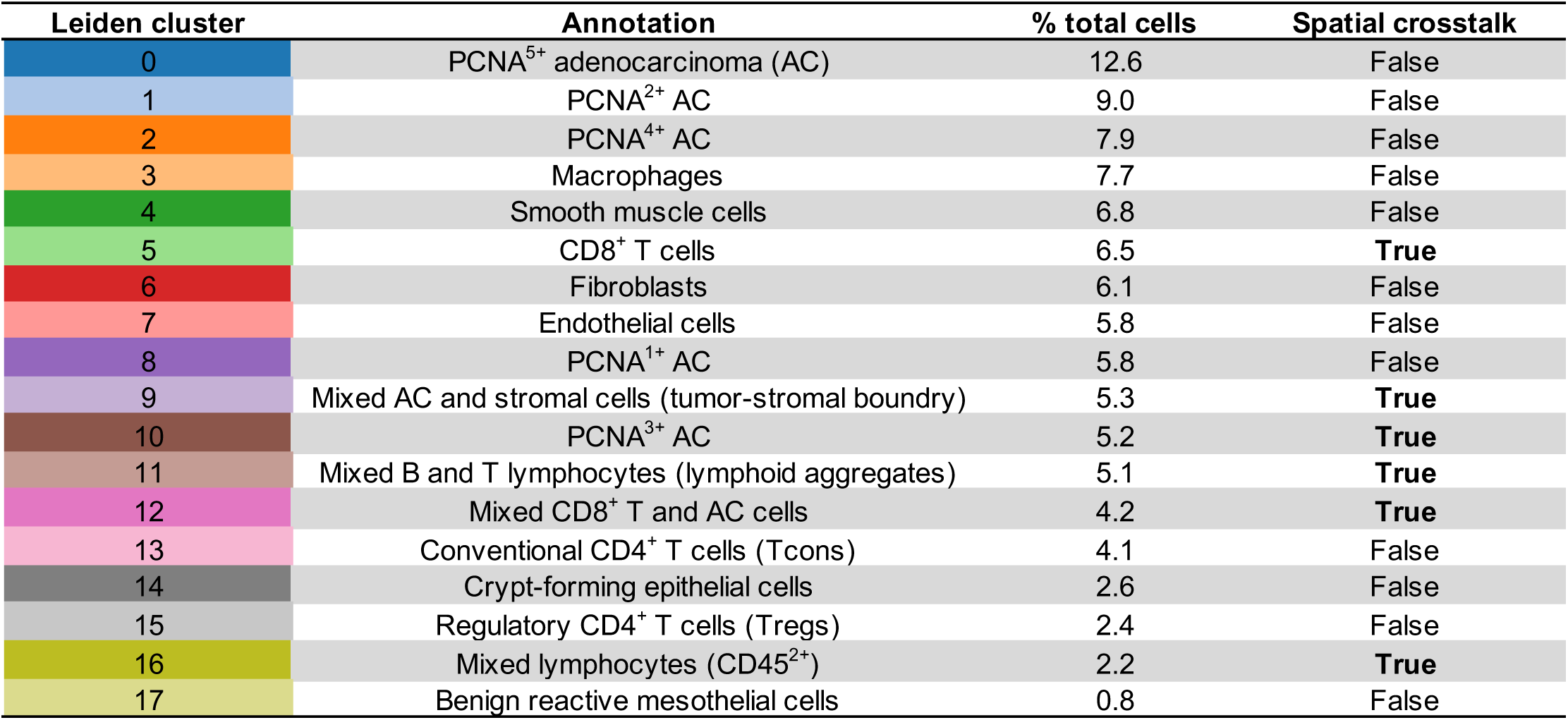
Segmentation-based Leiden cluster annotations. The column labeled “% total cells” indicates the percentage of total cells in CyCIF-1A accounted for by that cluster. Column labeled “Spatial crosstalk” indicates whether signal spillover across segmentation boundaries of neighboring cells was observed in CyCIF-1A.

**Supplementary Video 1.**
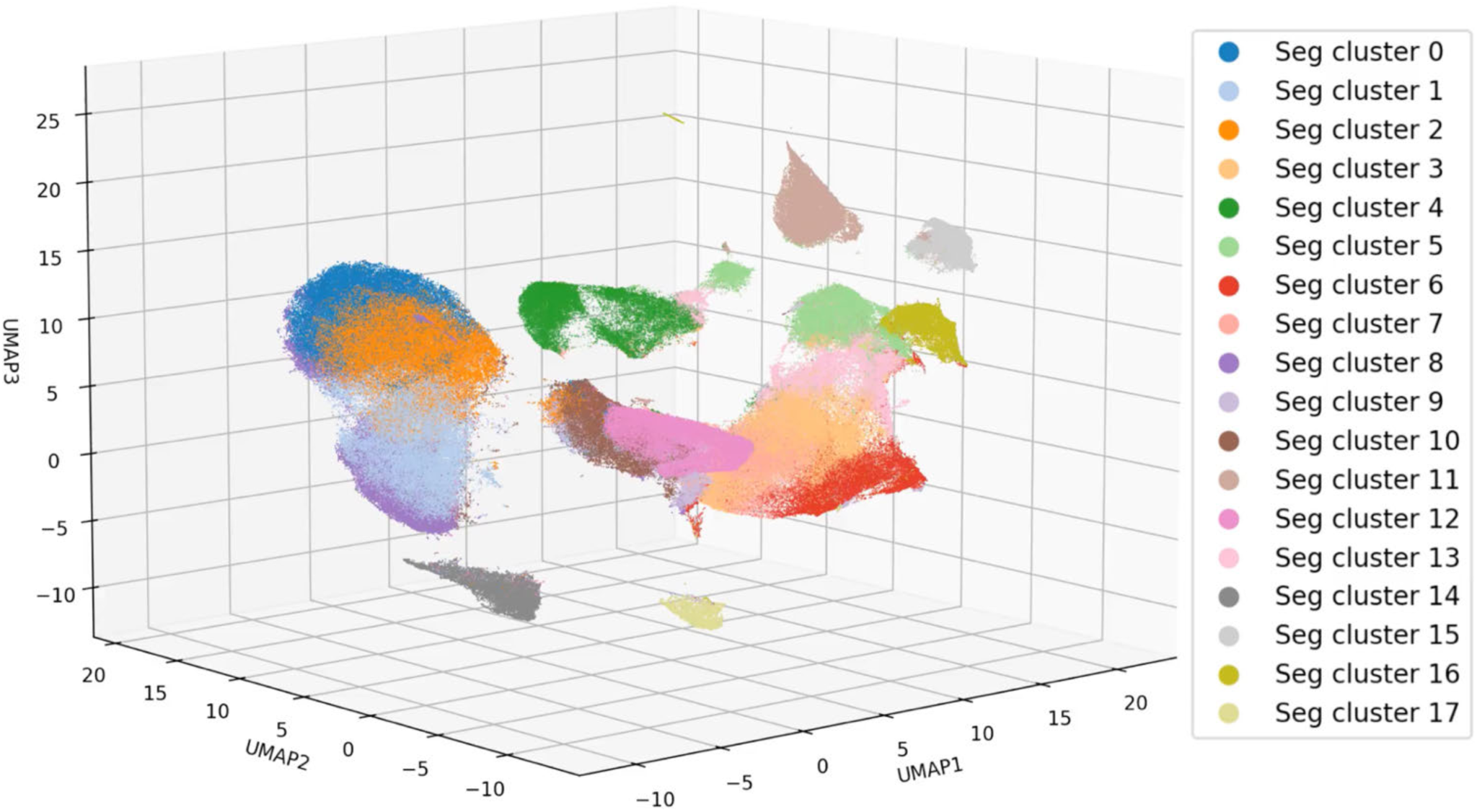
360° UMAP plot rotation showing segmented cells colored by Leiden cluster.

**Supplementary Fig. 2.**
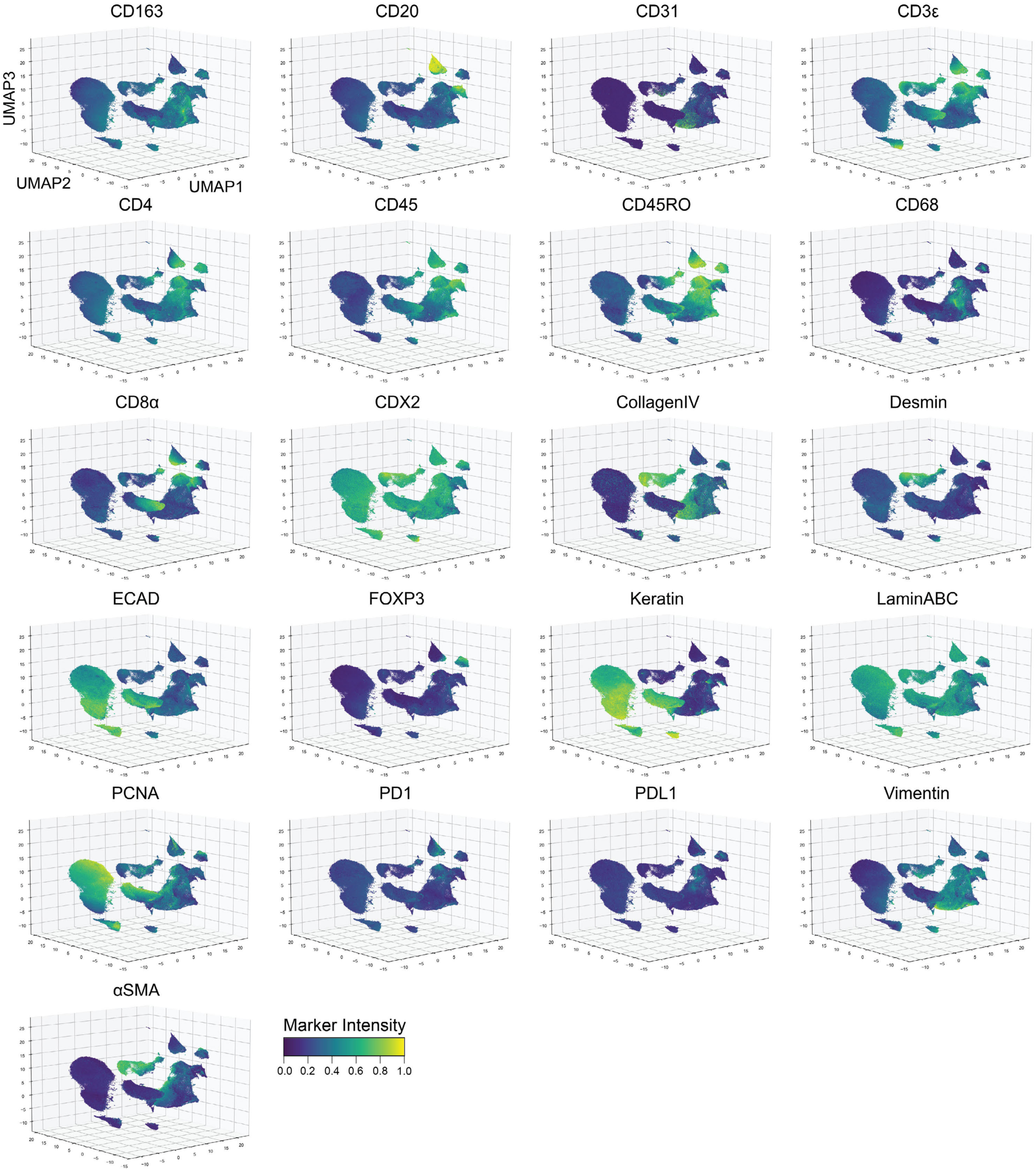
Channel colormaps for segmented cells in CyCIF-1A. UMAP embeddings of segmented cells colored by mean marker intensity for 21 immunomarkers.

**Extended Data Fig. 2.**
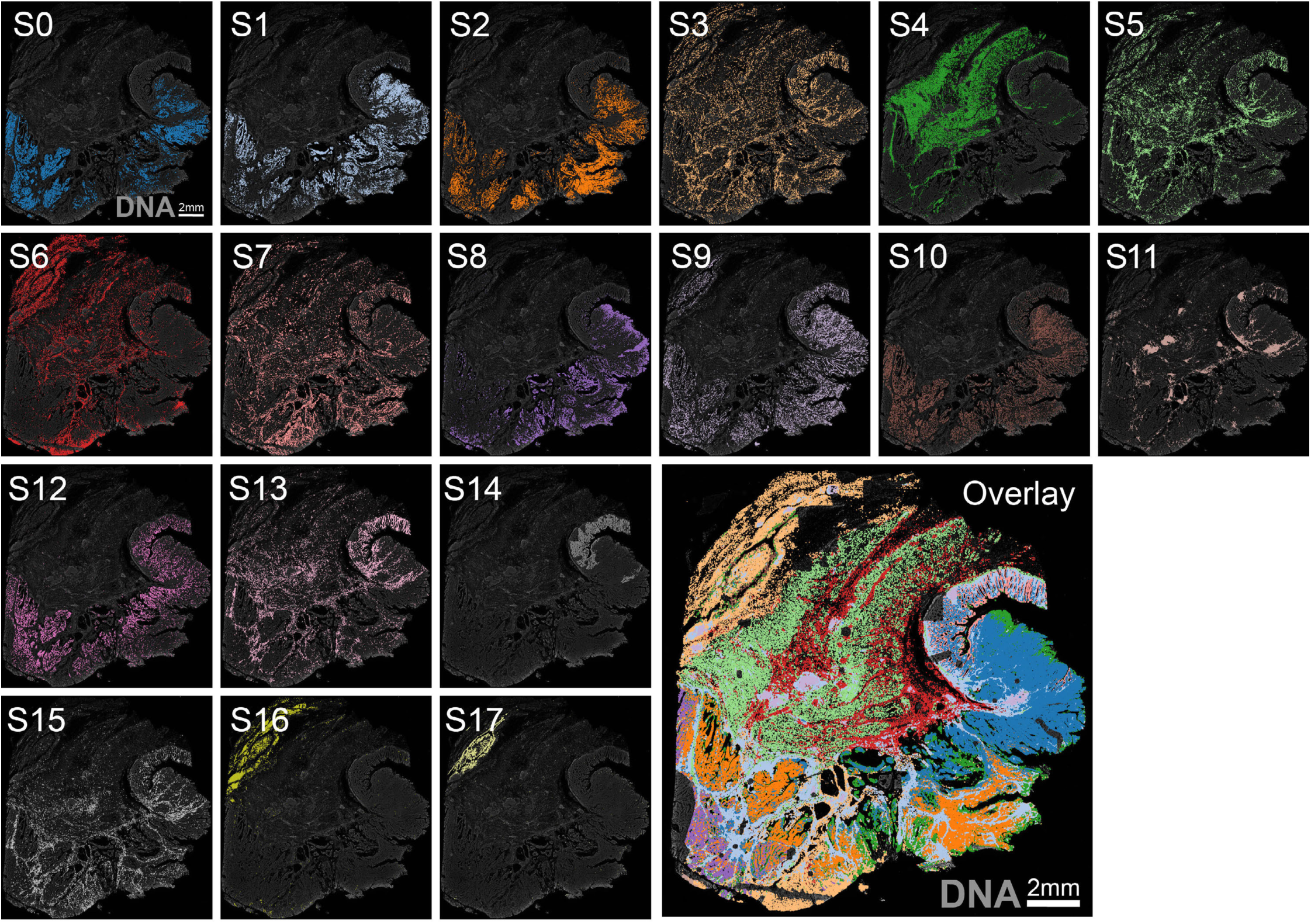
Location of segmentation-based Leiden clusters in CyCIF-1A. DNA (gray background) shown for reference.

**Supplementary Fig. 3.**
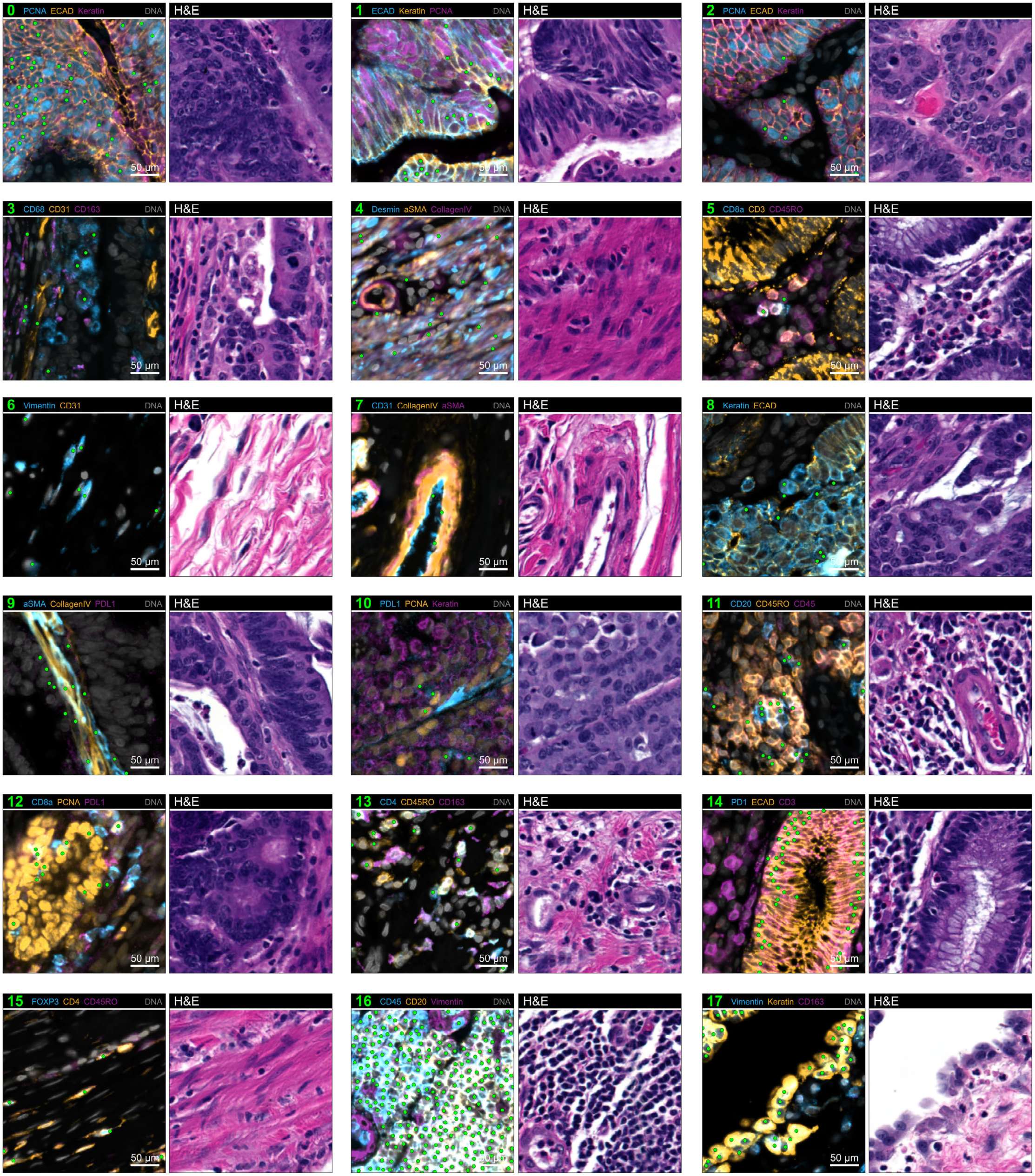
Fields-of-view showing the location of segmentation-based Leiden clusters in CyCIF-1A. Paired CyCIF (left) and H&E (right) fields-of-view in CyCIF-1A showing the location of random examples of cells from all 18 Leiden clusters (green scatter points in CyCIF images).

**Supplementary Fig. 4.**
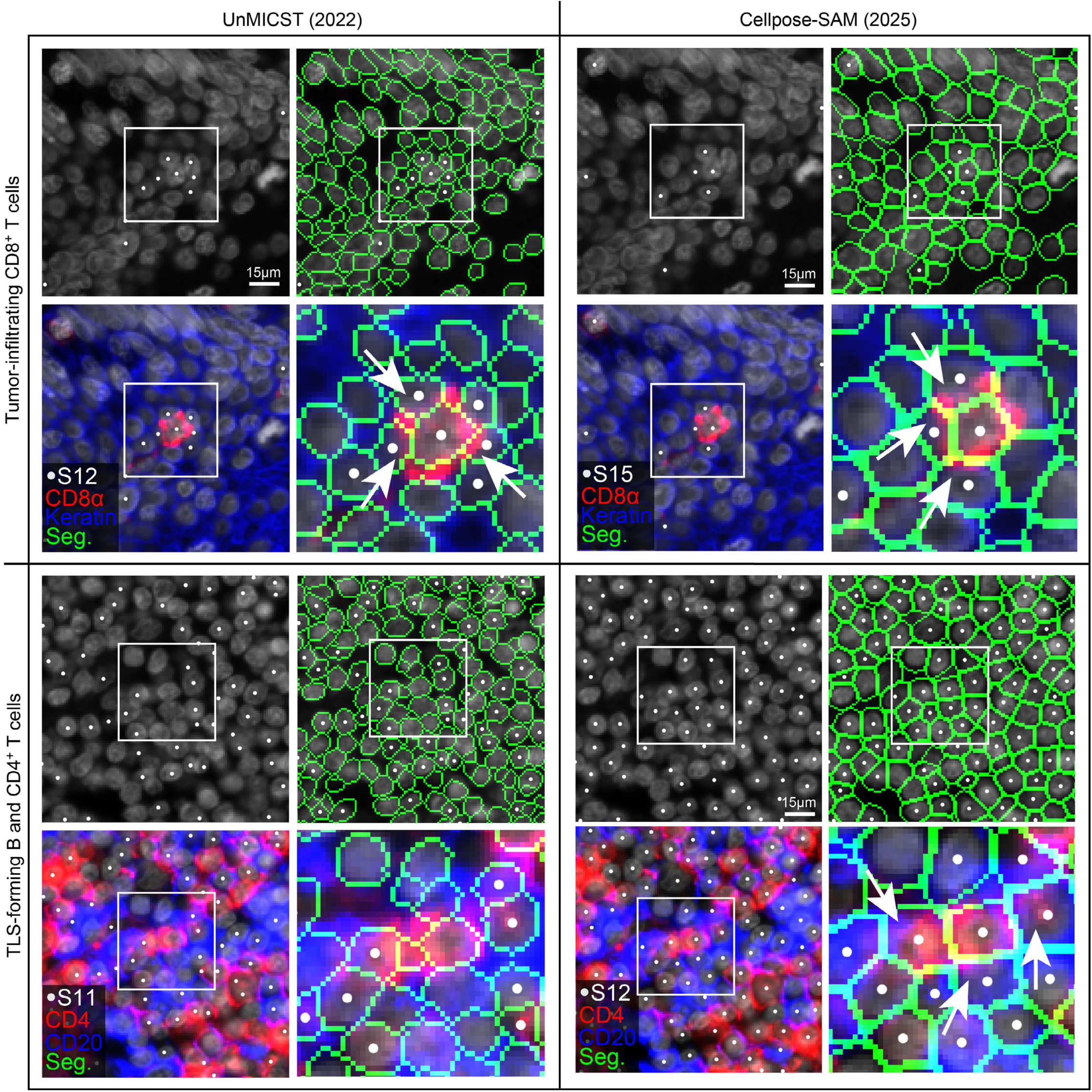
CNN-based (UnMICST) and transformer-based (Cellpose-SAM) segmentation algorithms are affected by spatial crosstalk, independent of model accuracy. Since neighboring cells can have overlapping membrane extensions (e.g. lamellipodia) or overlapping cell bodies along the Z-axis, segmentation masks often include portions of adjacent cells of different types referred to as “spatial crosstalk”. As a result, even state-of-the-art transformer-based segmentation methods like Cellpose-SAM^55^ introduce unavoidable crosstalk errors when cell boundaries are imposed on an image. Top two rows: UnMICST (left column) and Cellpose-SAM (right column) segmentation outlines in the same field-of-view in CyCIF-1A showing that, in both cases, tumor-infiltrating CD8^+^ T cells (red) and neighboring keratin^+^ tumor cells (blue) fall into the same cluster due to the sharing of signals across segmentation boundaries (white arrows) independent of segmentation quality. Bottom two rows: UnMICST (left column) and Cellpose-SAM (right column) segmentation outlines in the same field-of-view in CyCIF-1A showing that, in both cases, lymphoid aggregates consisting of B cells (blue) and CD4^+^ T cells (red) also co-cluster due to sharing of signals across segmentation boundaries (white arrows); an unavoidable consequence of spatial crosstalk.

**Supplementary Fig. 5.**
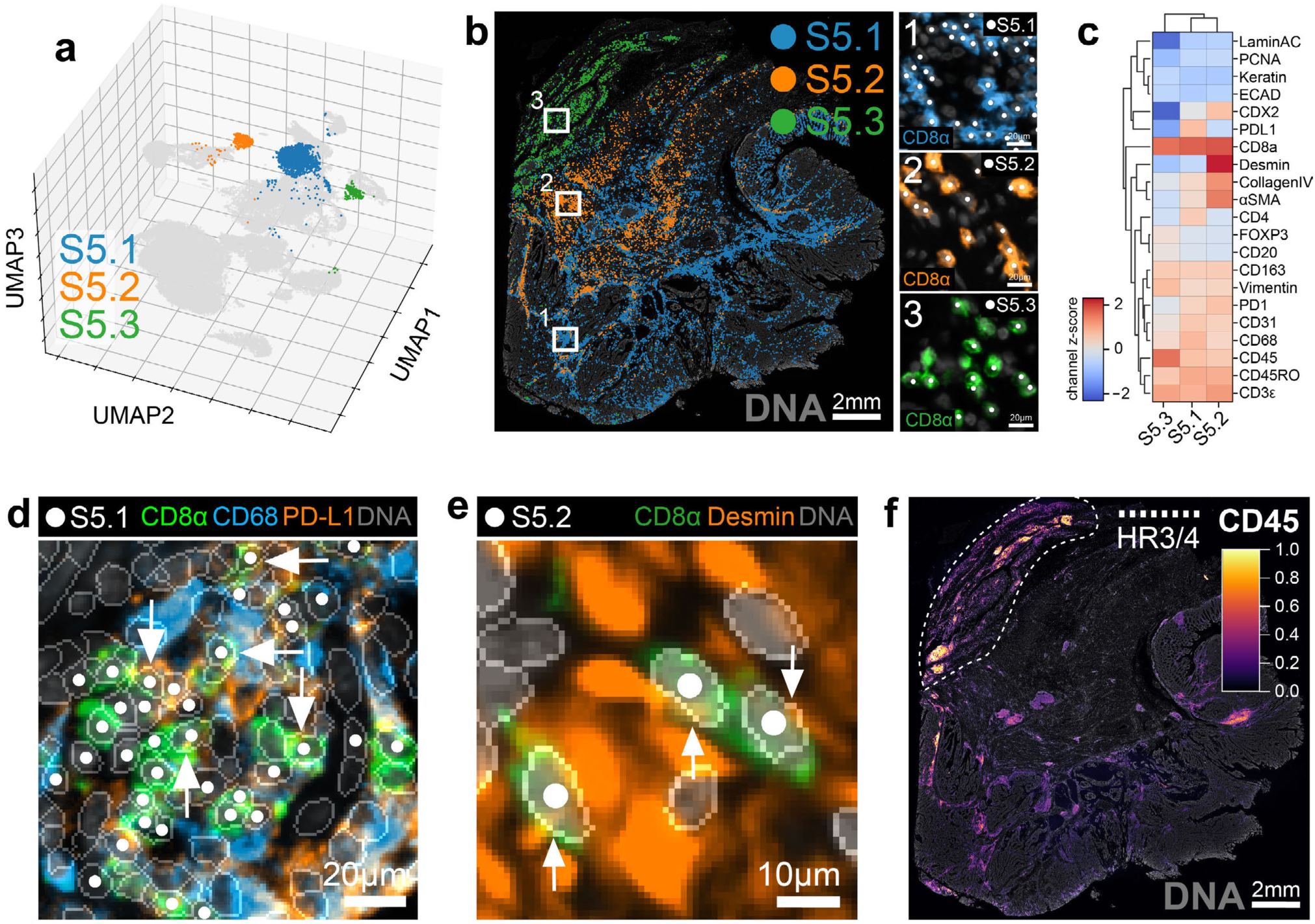
CD8^+^ T cells in segmentation cluster S5 are delocalized in UMAP space and affected by spatial crosstalk with cells in different tissue niches. **a**, 3D UMAP embedding of segmented cells from CyCIF-1A highlighting three subclusters of S5 cells: S5.1 (blue), S5.2 (orange), and S5.3 (green). **b**, Location of S5 subclusters in CyCIF-1A showing that they localize to different regions of tissue: S5.1 (peritumoral stroma, HR9), S5.2 (muscularis propria, HR2), S5.3 (sersosa/subserosa, HR3/4). Insets at right show that each subcluster represents a population of CD8^+^ T cells. **c**, Clustered heatmap of z-score-normalized mean marker intensities for S5 subclusters computed relative to all 18 segmentation-based Leiden clusters (not only S5 subclusters) revealing differences in their protein expression profiles. **d**, Enlarged view of the region highlighted by box 1 in panel b revealing spatial crosstalk between CD8α^+^ T cells from S5.1 (white scatter points) and neighboring CD68^+^ PD-L1^+^ macrophages. White arrows point to instances of signal spillover across segmentation boundaries. **e**, Enlarged view of the region highlighted by box 2 in panel b showing spatial crosstalk between CD8α^+^ T cells from S5.2 cells (white scatter points) and neighboring Desmin^+^ smooth muscle cells. White arrows point to instances of signal spillover across segmentation boundaries. **f**, CyCIF-1A as seen in the CD45 channel (colormap) showing that immune cells in the serosa/subserosa (HR3/4), in which CD8^+^ T cells in S5.3 reside, exhibit higher CD45 intensity compared to other areas of the tissue, helping to explain why these cells form their own cluster.

**Supplementary Fig. 6.**
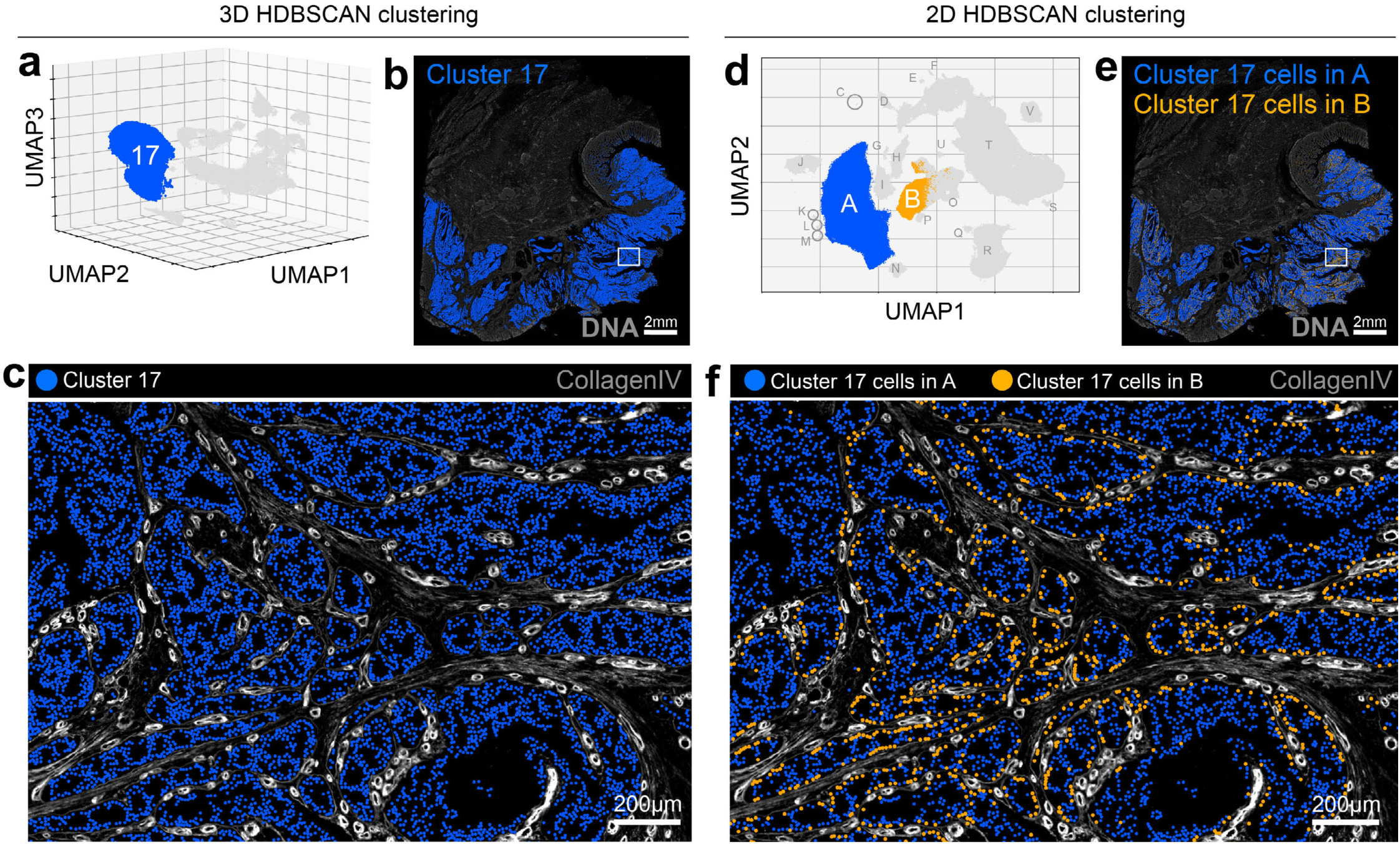
Cells affected by spatial crosstalk form discrete clusters on embedding in low-dimensional feature space. **a**, 3D UMAP embedding of segmented cells from CyCIF-1A with HDBSCAN cluster 17 highlighted in blue. (HDBSCAN is a density-based clustering algorithm that was applied directly to the embedded data points.) **b**, Location of cluster 17 cells in CyCIF-1A. **c**, Enlarged view of the region highlighted by the white box in panel b showing that cluster 17 cells do not include collagen IV^+^ (gray) stromal cells at the tumor-stromal interface. **d**, 2D UMAP embedding of the same segmented cells from CyCIF-1A showing that cluster 17 cells from panel now form two discrete clusters (A, blue and B, orange) on embedding in 2D feature space. **e**, Location of cells from clusters A and B in CyCIF-1A. **f**, Enlarged view of the region highlighted by the white box in panel e showing that cluster B represents tumor cells at the tumor-stromal boundary affected by spatial crosstalk with collagen IV^+^ (gray) stromal cells.

**Extended Data Fig. 3.**
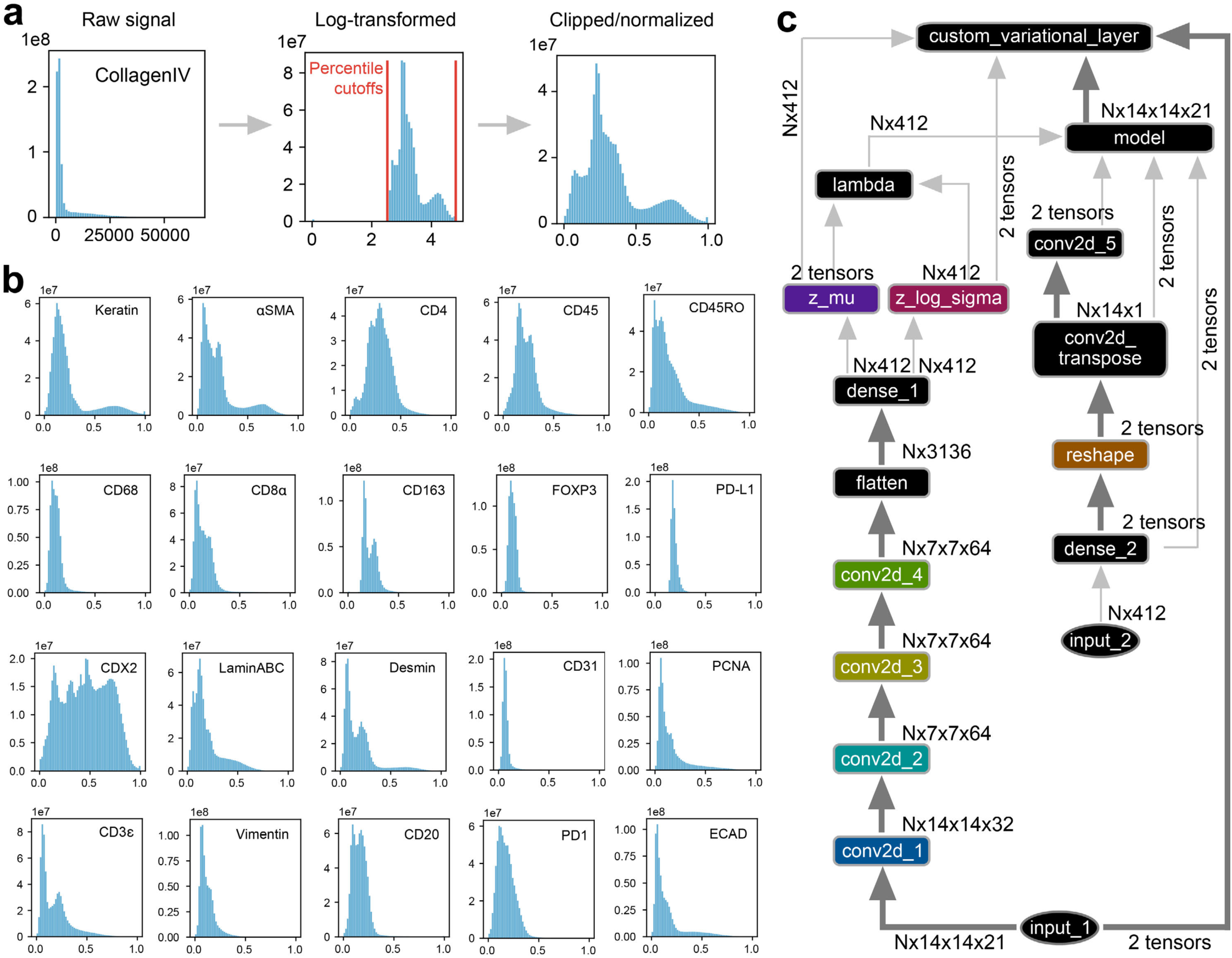
VAE network architecture and image patch preprocessing. **a**, Preprocessing steps undergone by image patches prior to entering the VAE encoding network using the collagen IV channel as an example showing the raw pixel intensity distribution for the full CyCIF-1A image channel (right histogram), the same data after log-transformation (center histogram) and log-transformed data after clipping and 0-1 normalization (right histogram). Lower (0.17^th^ percentile) and upper (99.99^th^ percentile) clipping cutoffs are highlighted by vertical red bars in the center histogram. **b**, Clipped and normalized channel intensity distributions for the other 20 immunomarkers analyzed in this study. **c**, Schematic of the VAE network architecture used in this study. Conv2d = 2D convolutional layer, N = image patch batch size.

**Supplementary Table 4.**
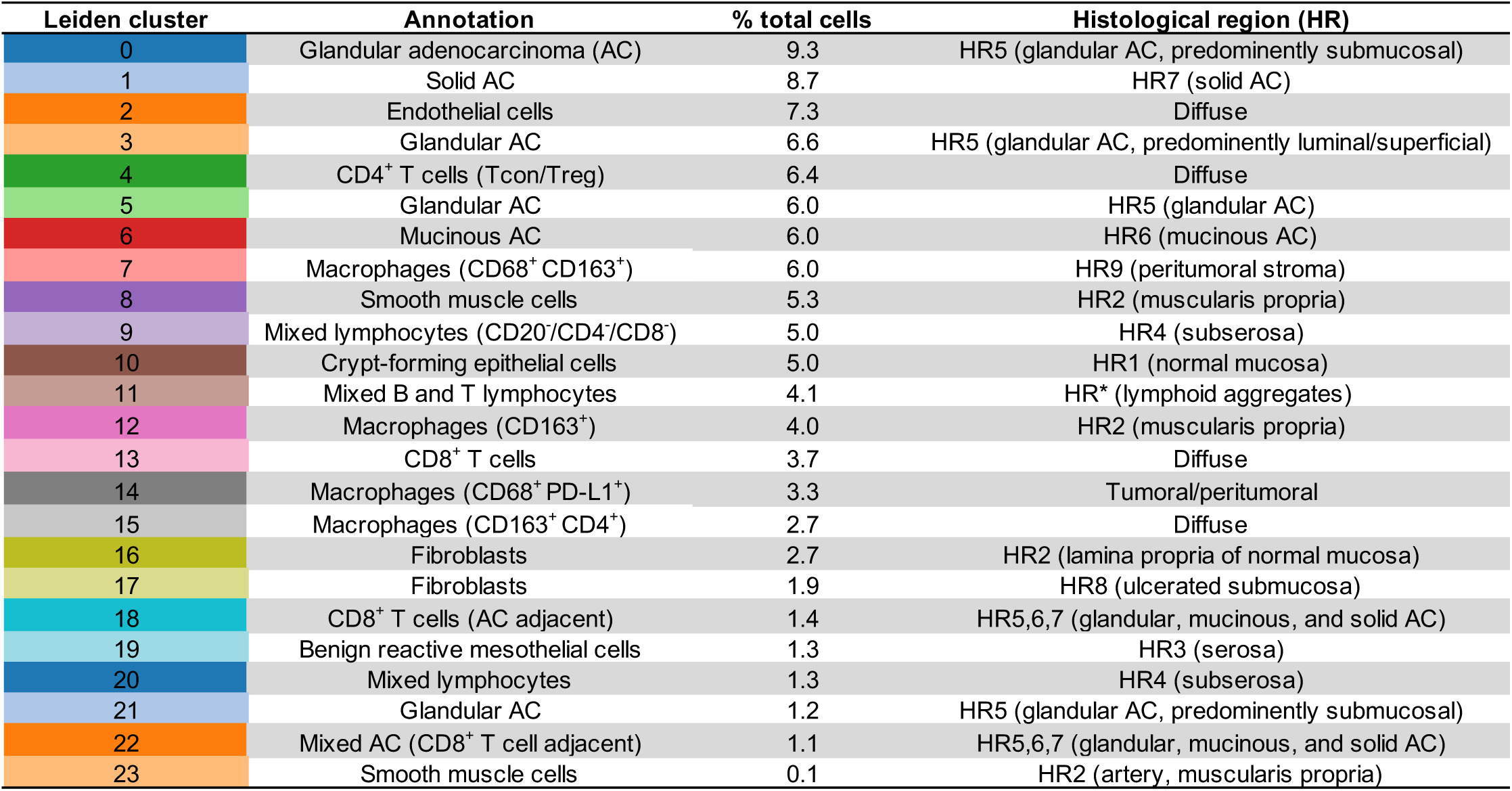
9x9µm VAE image patch Leiden cluster annotations.

**Supplementary Video 2.**
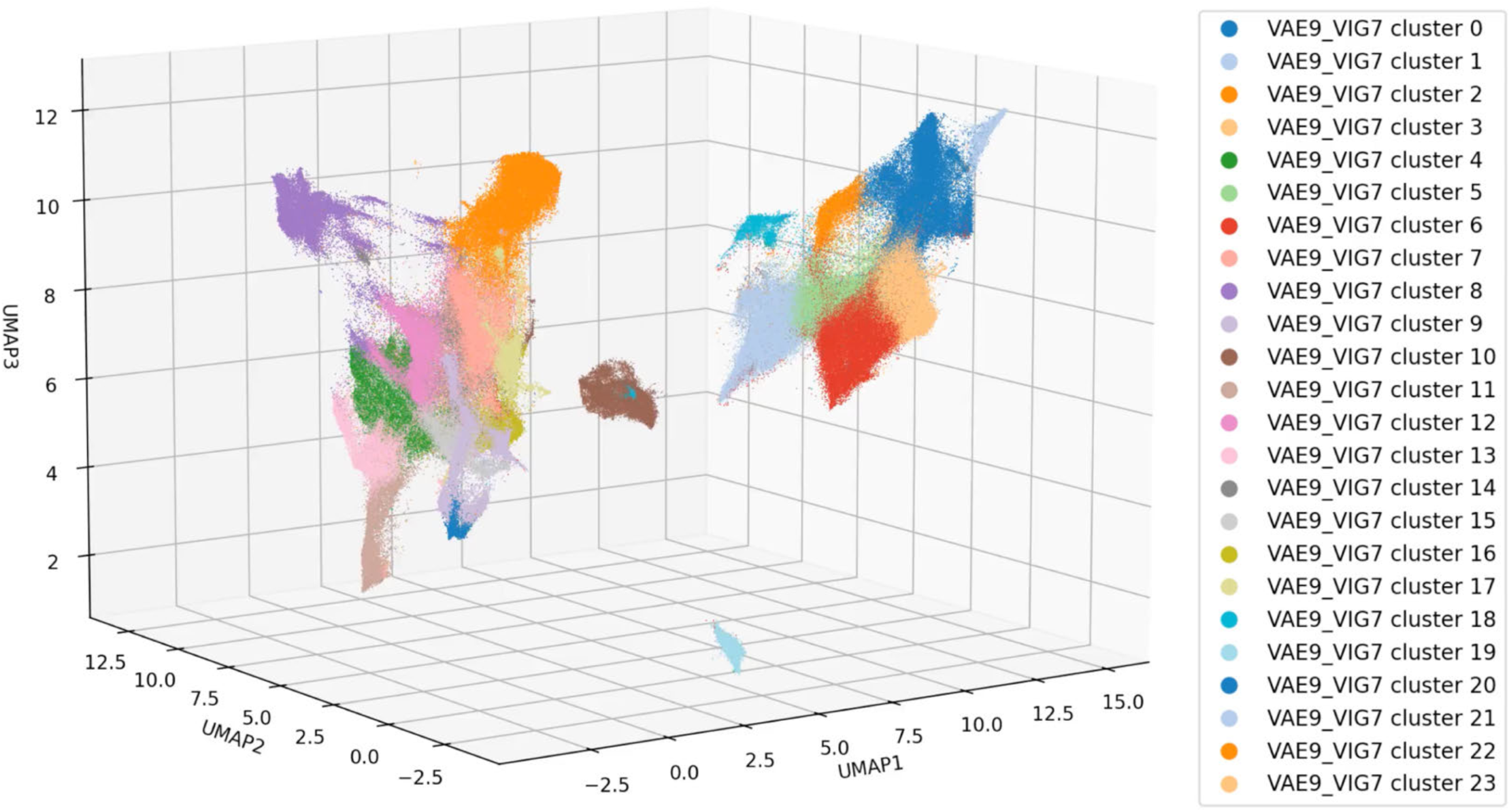
360° UMAP plot rotation showing 9x9µm VAE image patch encodings colored by Leiden cluster label.

**Extended Data Fig. 4.**
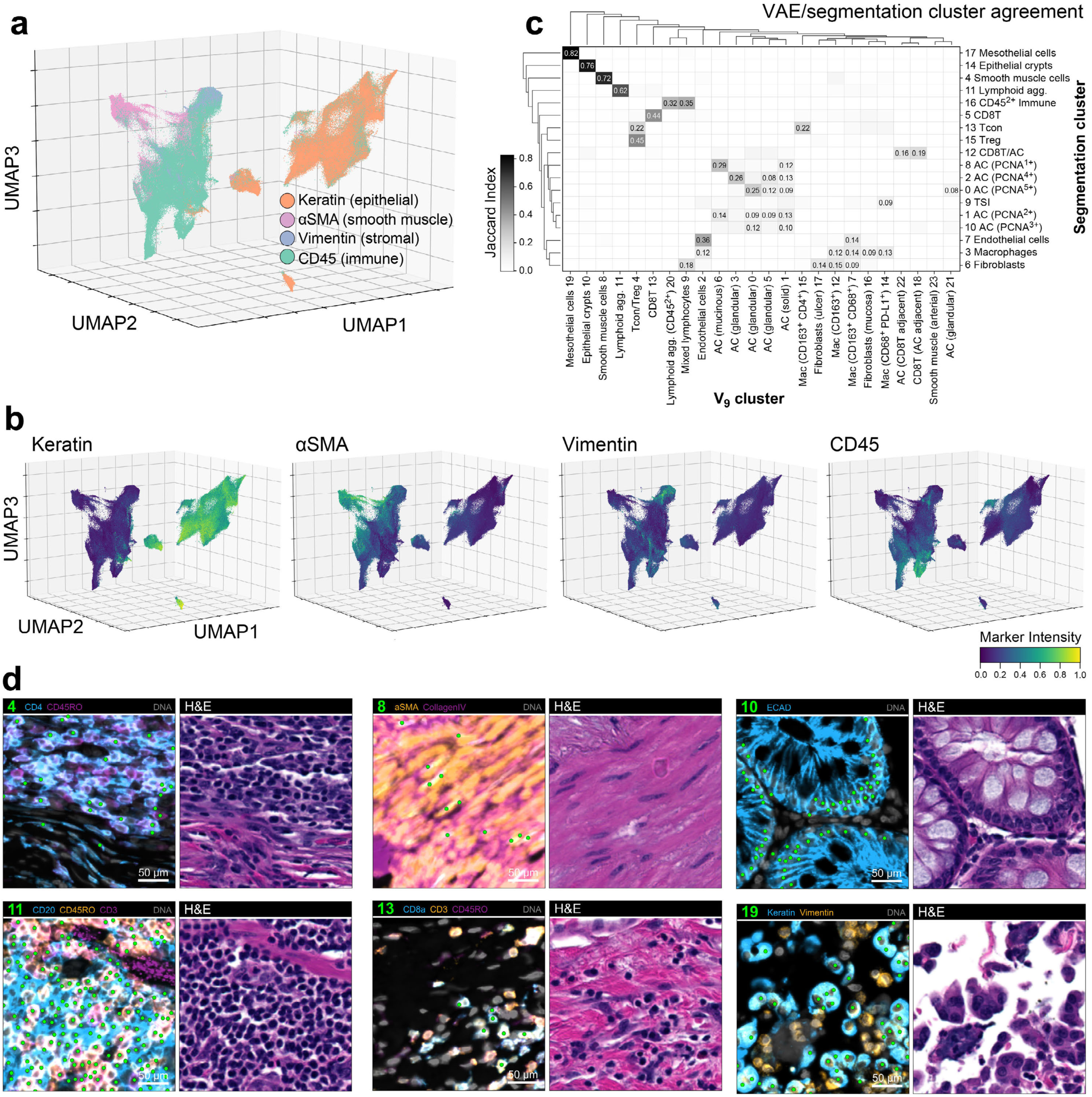
VAE-based classification of human CRC at single-cell resolution. **a**, 3D UMAP embedding of 9x9µm VAE image patch encodings from CyCIF-1A colored by major cell lineage markers: keratin (epithelial, orange), αSMA (smooth muscle, pink), vimentin (stromal, blue), and CD45 (immune, green). **b**. 3D UMAP embedding of VAE image patch encodings from CyCIF-1A colored by pixel-level median marker intensity of major cell lineage markers keratin, αSMA, vimentin, and CD45. See **Supplementary Fig. 7** for colormaps of all 21 immunomarkers. **c**, Jaccard similarity indices (defined as the ratio of the intersection over the union of two sets) for cells in pairs of VAE and segmentation-based clusters. Possible values range from 0 (no overlap) to 1 (perfect overlap). **d**, Paired CyCIF (left) and H&E (right) fields-of-view from CyCIF-1A showing the location of random examples of cells from various Leiden clusters (green scatter points in CyCIF images). See **Supplementary Fig. 8** for views of all 24 clusters.

**Supplementary Fig. 7.**
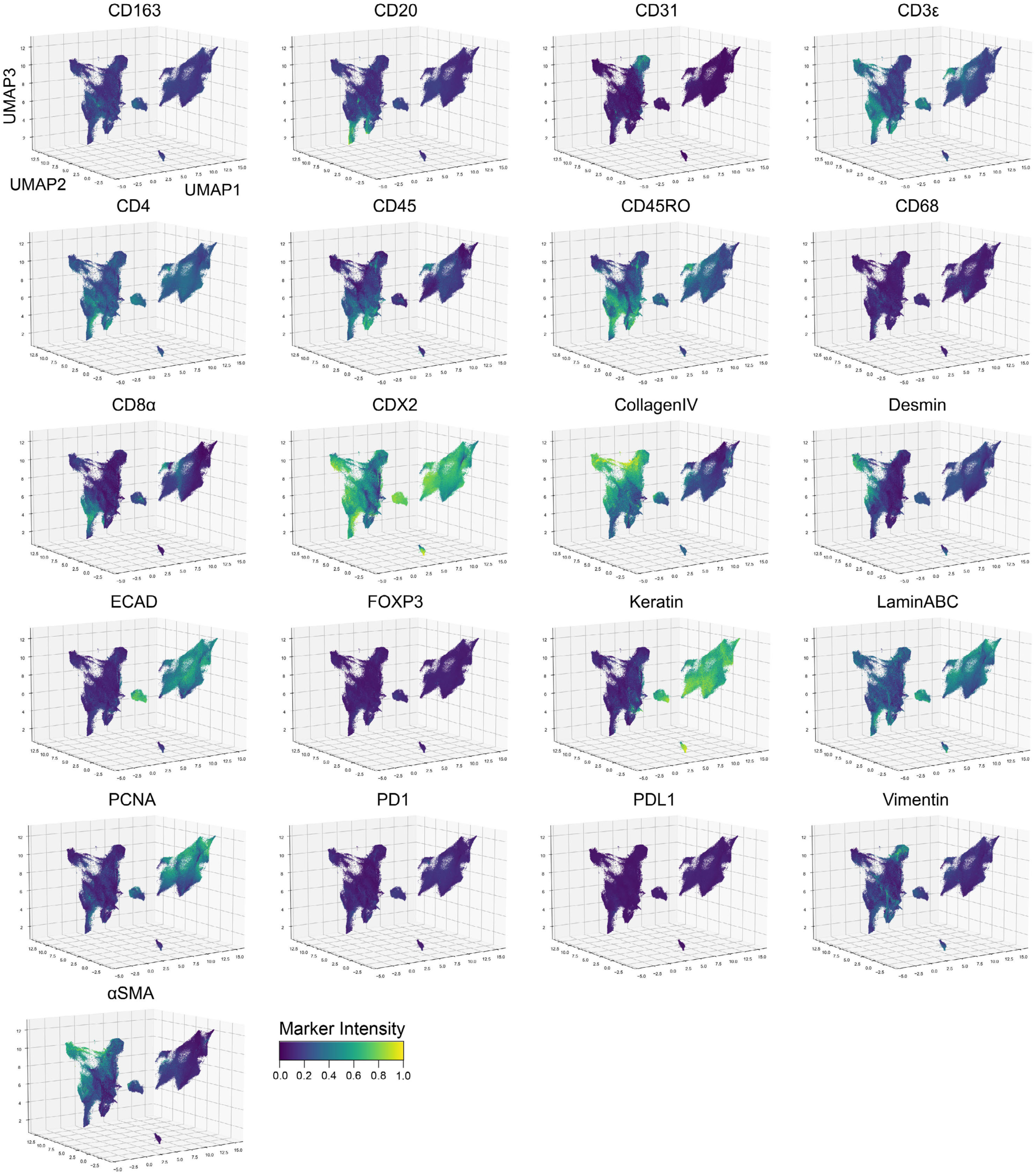
Channel colormaps for 9x9µm VAE image patch encodings from CyCIF-1A. UMAP embeddings of VAE image patch encodings colored by pixel-level median marker intensity for 21 immunomarkers.

**Extended Data Fig. 5.**
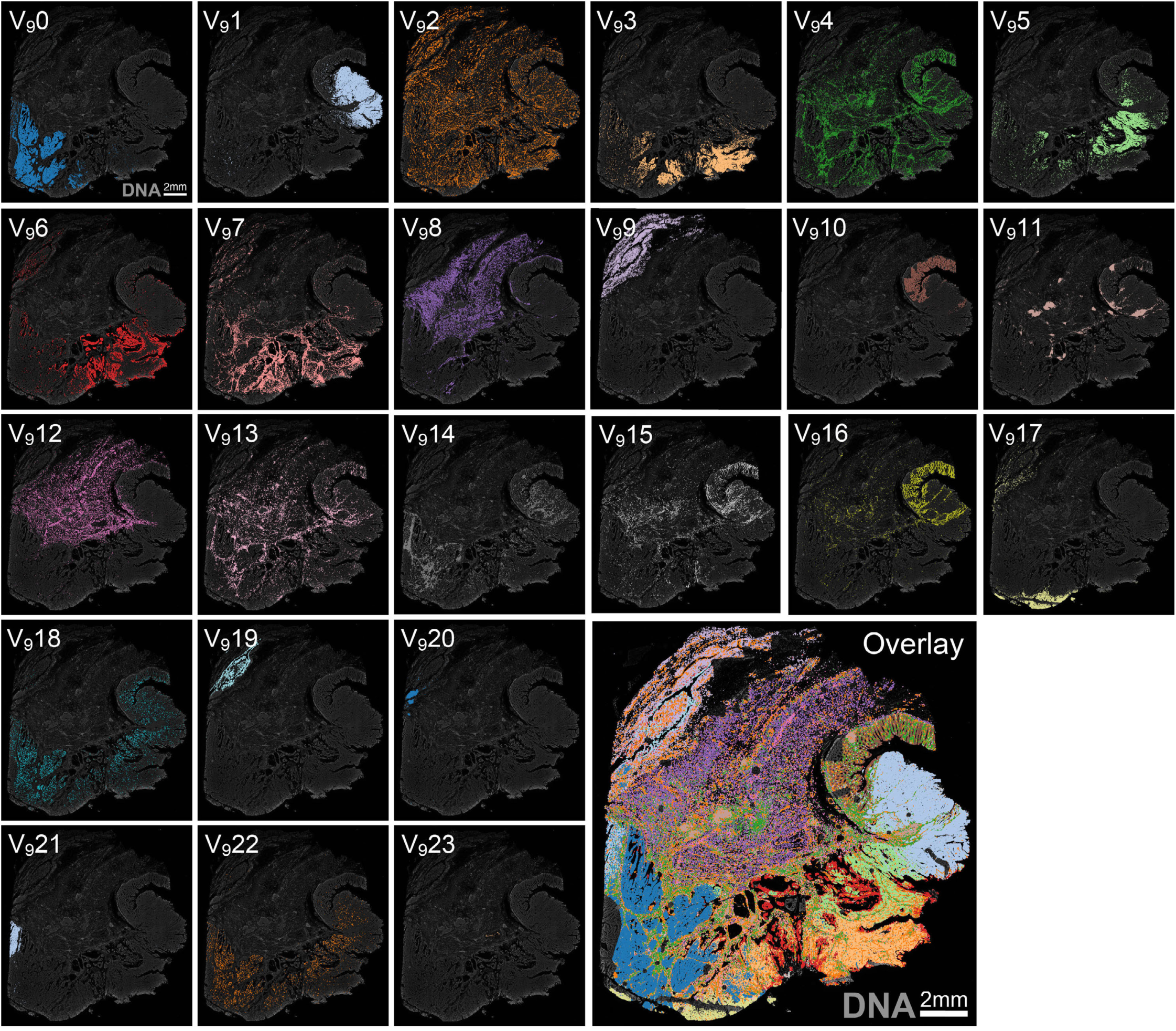
Location of 9x9µm VAE image patch Leiden clusters in CyCIF-1A. DNA (gray background) shown for reference.

**Supplementary Fig. 8.**
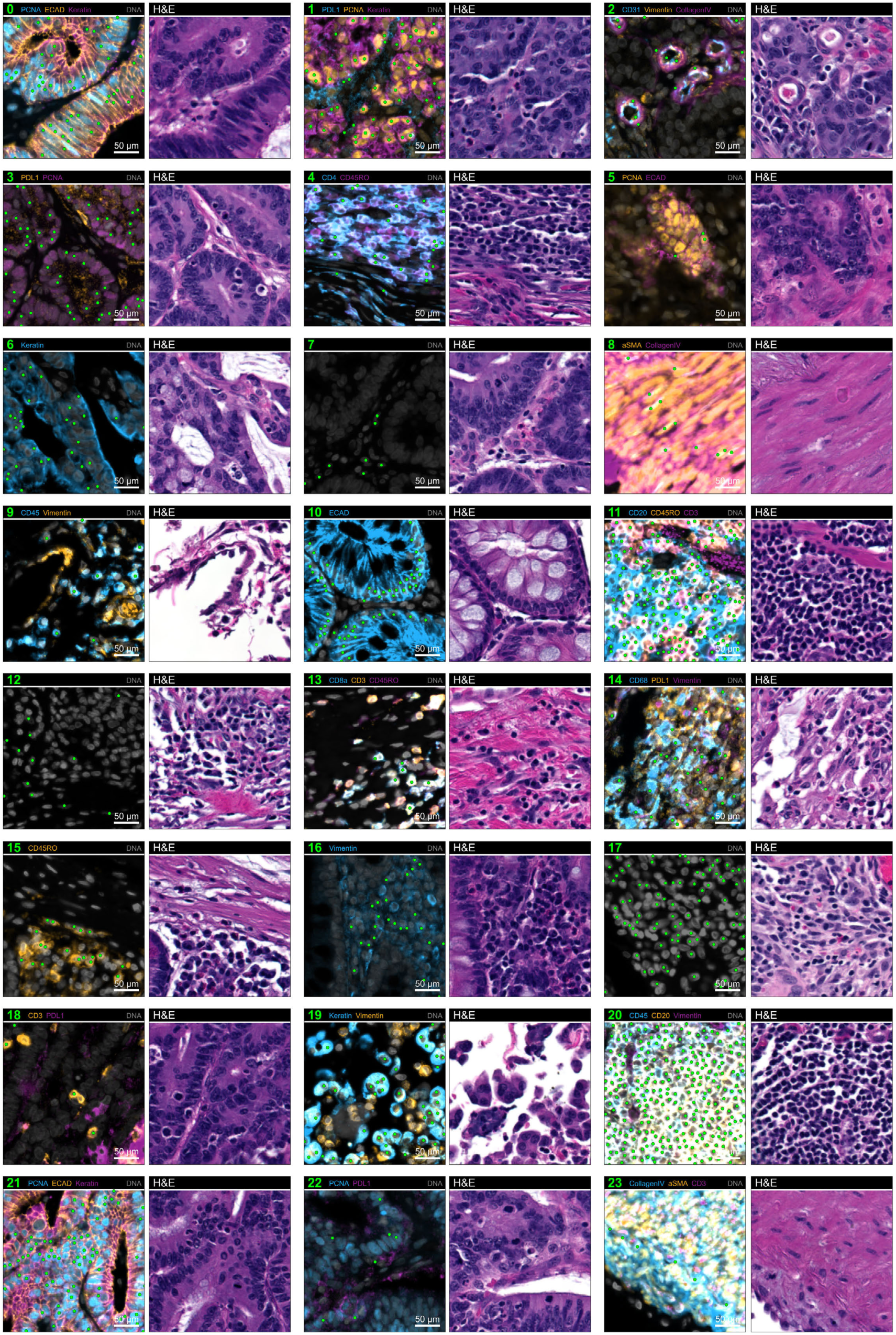
Fields-of-view showing the location of 9x9µm VAE image patch Leiden clusters in CyCIF-1A. Paired CyCIF (left) and H&E (right) fields-of-view in CyCIF-1A showing the location of random examples of cells from all 24 Leiden clusters (green scatter points in CyCIF images).

**Extended Data Fig. 6.**
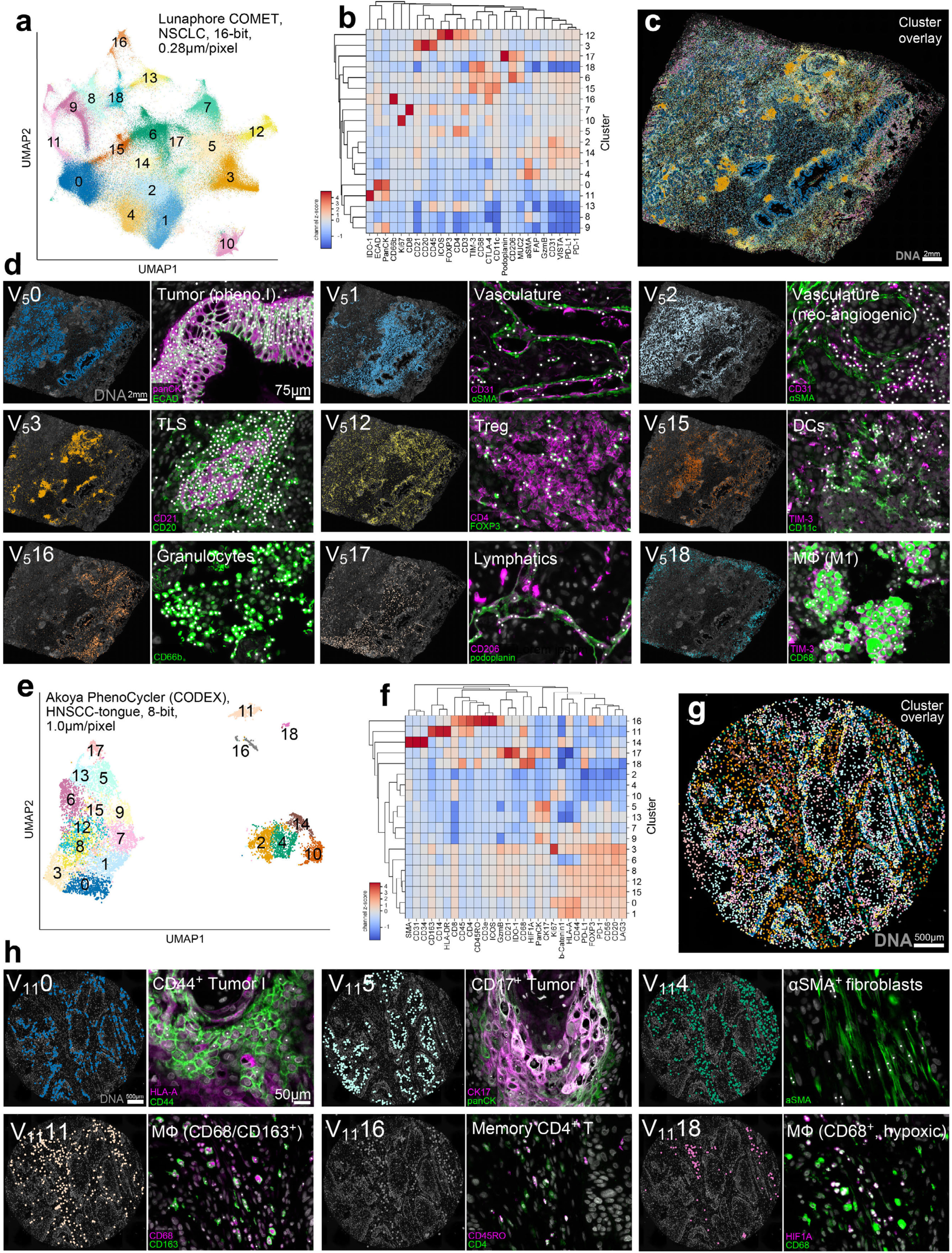
VAE-based classification of Lunaphore and PhenoCycler data. **a-d**, 27-plex Lunaphore COMET imaging of human NSCLC: **a**, 2D UMAP embedding of 5x5µm VAE image patch encodings colored by Leiden cluster. **b**, Hierarchical clustering of z-score-normalized mean pixel-level marker intensities for Leiden clusters shown in panel a. **c**, Location of Leiden clusters in Lunaphore-1; DNA (gray) shown for reference. **d**, Examples of individual clusters showing their locations (left panels) and phenotypes (right panels) as seen in characteristic image channels. See **Supplementary Fig. 9** for views of all 19 clusters. **e-h**, 29-plex Akoya PhenoCycler imaging of human tongue squamous cell carcinoma: **e**, 2D UMAP embedding of 11x11µm VAE image patch encodings colored by Leiden cluster. **f**, Hierarchical clustering of z-score-normalized mean pixel-level marker intensities for Leiden clusters shown in panel e. **g**, Location of Leiden clusters in CODEX-1; DNA (gray) shown for reference. **h**, Examples of individual clusters showing their locations (left panels) and phenotypes (right panels) as seen in characteristic image channels. See **Supplementary Fig. 10** for views of all 19 clusters.

**Supplementary Fig. 9.**
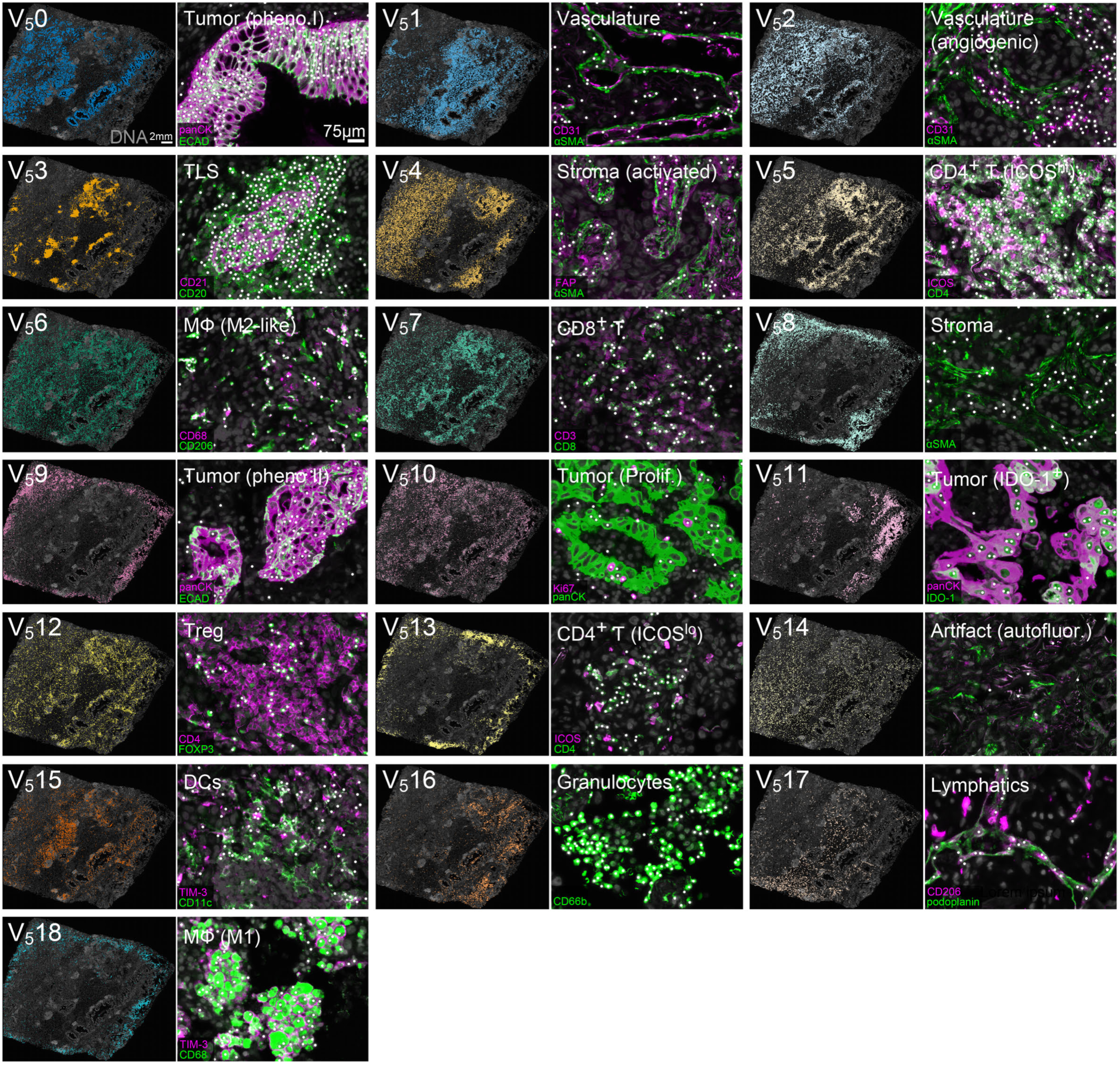
Locations and phenotypes of 5x5µm VAE image patch Leiden clusters in Lunaphore-1. Left panels show the overall spatial distribution of clustering cells with DNA (gray) for reference. Right panels show examples of clustering cells (white points) as seen in their characteristic marker channels.

**Supplementary Fig. 10.**
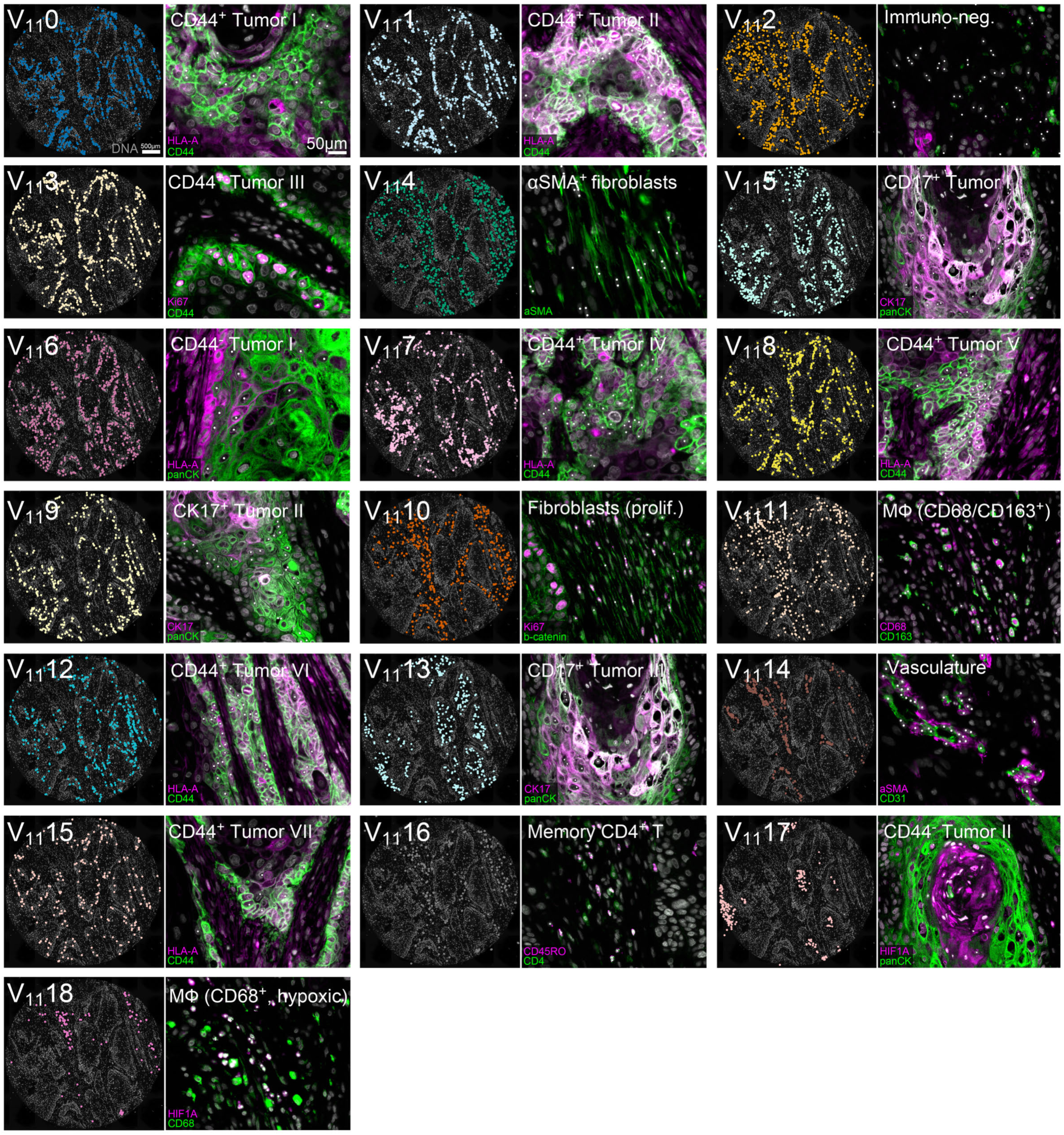
Locations and phenotypes of 11x11µm VAE image patch Leiden clusters in CODEX-1. Left panels show the overall spatial distribution of clustering cells with DNA (gray) for reference. Right panels show examples of clustering cells (white points) as seen in their characteristic marker channels.

**Extended Data Fig. 7.**
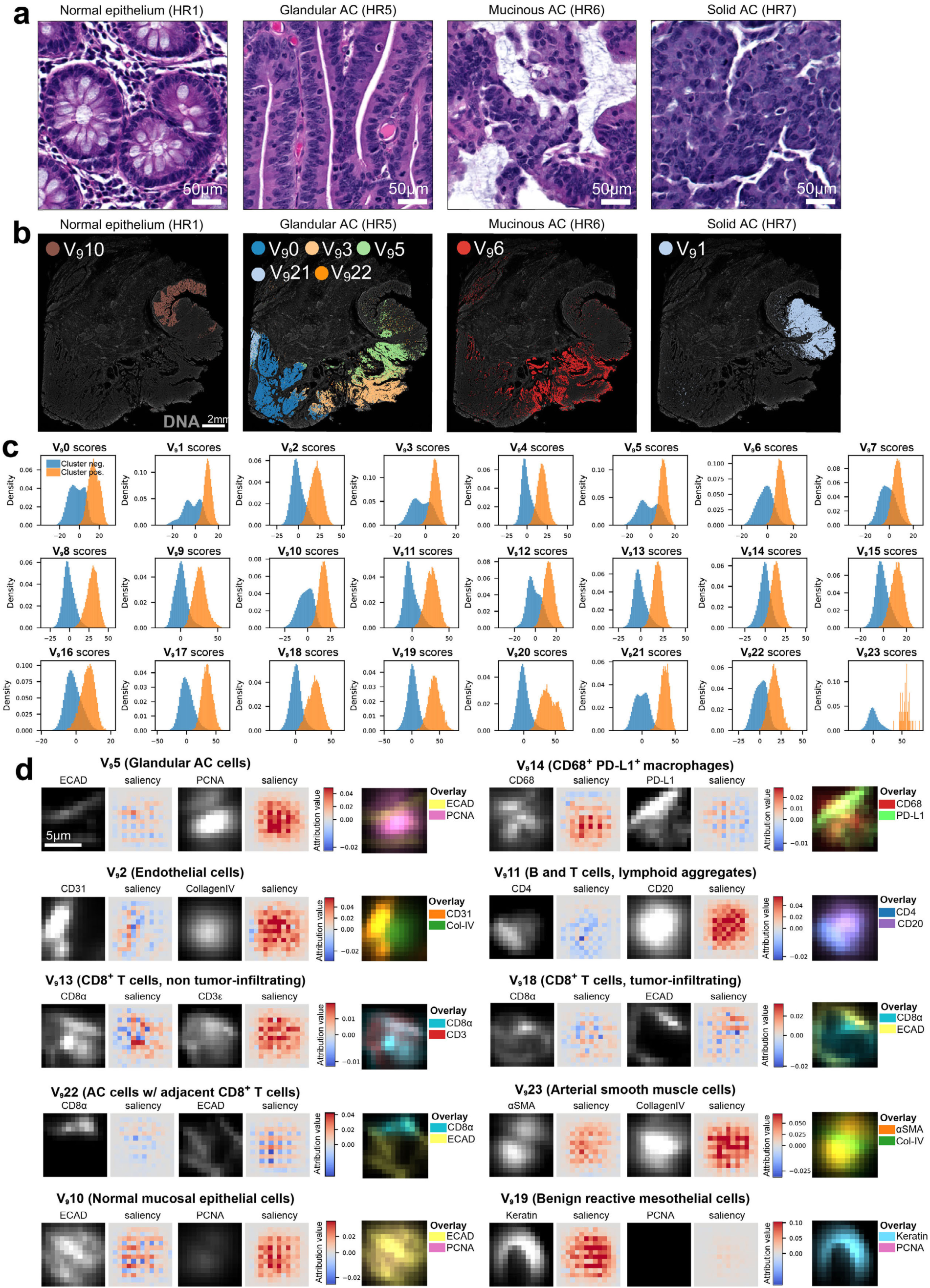
Morphology and concept saliency analysis for 9x9µm VAE image patch Leiden clusters. **a**, Fields-of-view from the H&E image (**Fig. 1b**) showing examples of various normal and cancerous epithelial histologies. **b**, Location of epithelial V_9_ clusters in CyCIF-1A grouped by histological region. See (**Fig. 1c**) for histological annotations. **c**, Distributions of concept scores (Sc), defined as the dot product between the i^th^ VAE image patch encoding (*Zi*) and the corresponding cluster concept vector (*Zc*) such that *Sc = Zi ⋅ Zc* for patches with (orange) and without (blue) a given cluster label. **d**, Channel intensities (left images) and corresponding concept saliency maps (right images) for 10 different V_9_ clusters. Overlay color images shown for reference; see **Supplementary Image Gallery 2** for fifteen examples of saliency maps across all 21 channels for all 24 V_9_ clusters.

**Extended Data Fig. 8.**
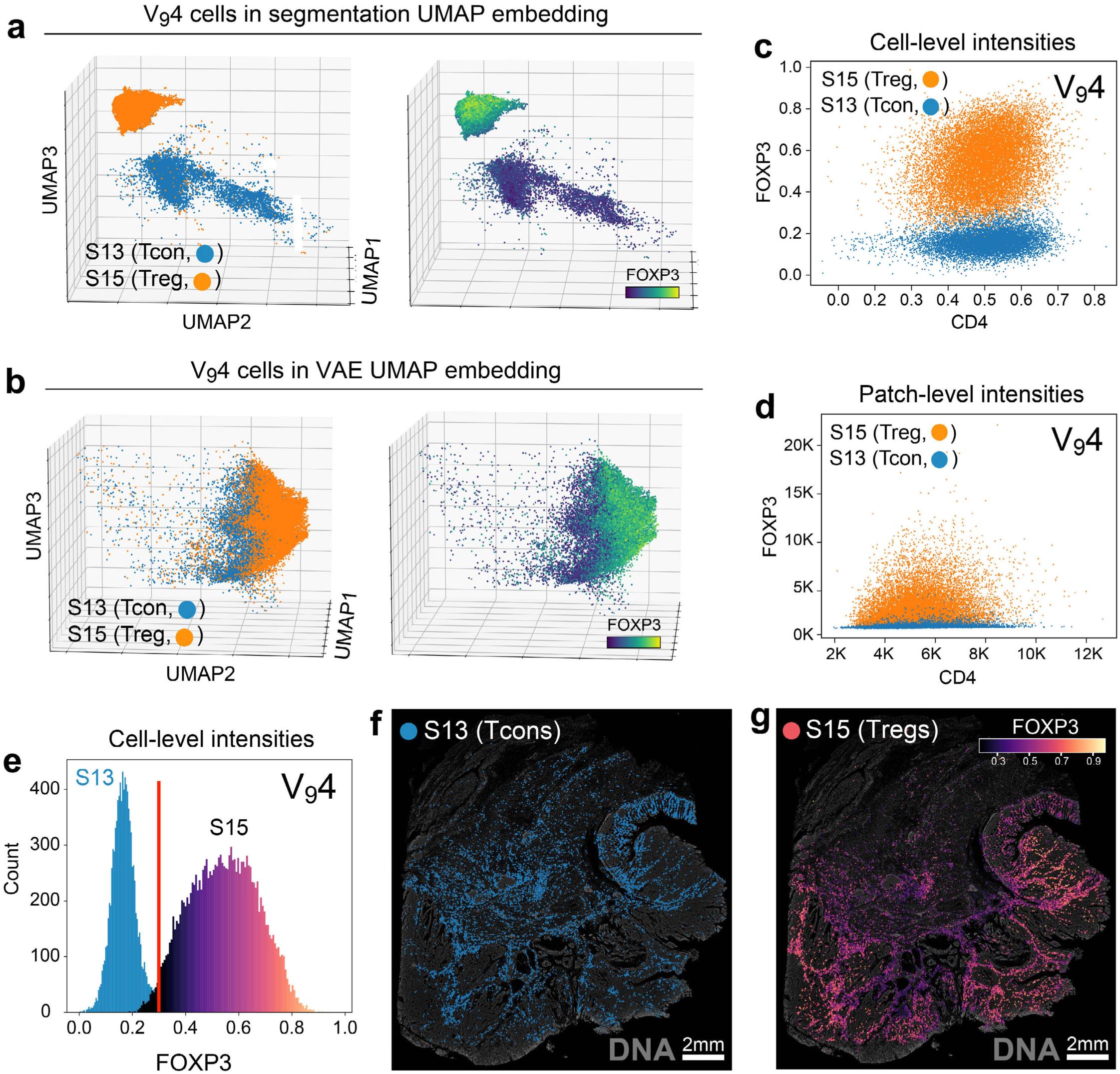
Tregs and Tcons exhibit greater similarity at the image patch level compared to the single-cell level. **a**, Distribution of V_9_4 cells in the segmentation-based UMAP embedding colored by S13 (Tcons, blue) and S15 (Tregs, orange) in the left plot and FOXP3 intensity in the right plot. **b**, Distribution of V_9_4 cells in the VAE-based UMAP embedding colored by S13 (Tcons, blue) and S15 (Treg, orange) in the left plot and FOXP3 intensity in the right plot. **c**, Scatter plot showing the distribution of V_9_4 cells according to CD4 (x-axis) and FOXP3 (y-axis) intensity at the level of segmented cells colored by S13 (Tcons, blue) and S15 (Tregs, orange) showing that the two cell types form discrete populations along the FOXP3 axis. **d**, Scatter plot showing the distribution of V_9_4 cells according to CD4 (x-axis) and FOXP3 (y-axis) intensity in 9x9µm image patches colored by S13 (Tcons, blue) and S15 (Tregs, orange) showing that the FOXP3 axis no longer exhibits a bimodal distribution. **e**, Distribution of V_9_4 cells according to FOXP3 intensity colored by S13 (Tcons, blue) and S15 (Tregs, colormap) again revealing a bimodal distribution for the two cell populations according to FOXP3 at the level of segmented cells. Red vertical line indicates a manually defined gate distinguishing positive (right of gate) and negative (left of gate) cell populations. **f**, Location of gated S13 cells (Tcons, FOXP3^-^ cells in panel e) in CyCIF-1A. **g**, Location of gated S15 cells (Tregs, FOXP3^+^ cells in panel e) in CyCIF-1A revealing that Tregs occupy similar spatial niches as Tcons in CyCIF-1A and tend to express higher FOXP3 levels in tumor regions (HR5, 6, and 7)

**Supplementary Video 3.**
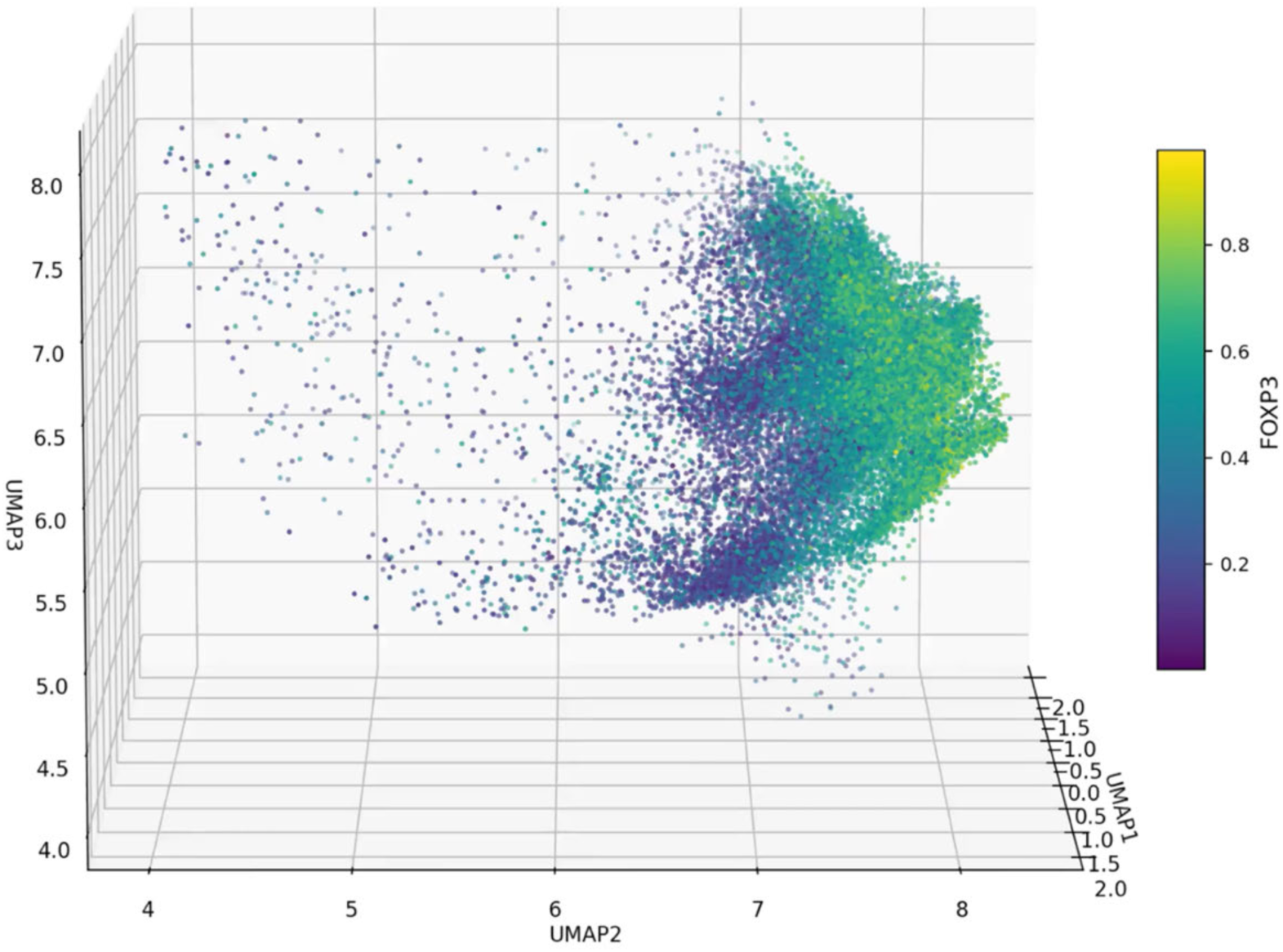
360° UMAP plot rotation showing 9x9µm VAE image patch encodings for VAE cluster 4 (V_9_4) colored by FOXP3 expression.

**Supplementary Table 5.**
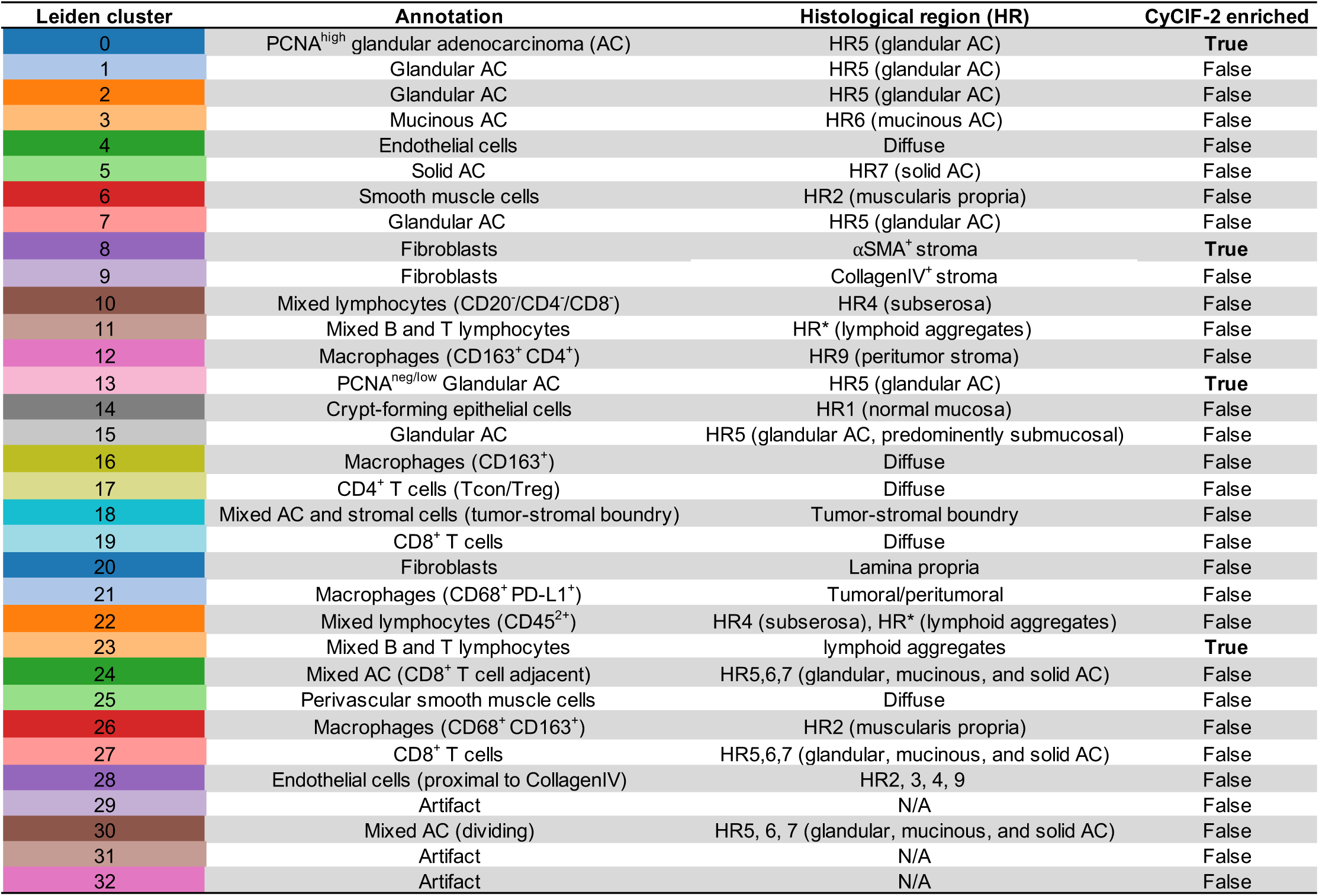
9x9µm VAE image patch Leiden cluster annotations (multi-tissue analysis). Column labeled “CyCIF-2 enriched” indicates whether the given cluster is predominantly found in that specimen.

**Supplementary Video 4.**
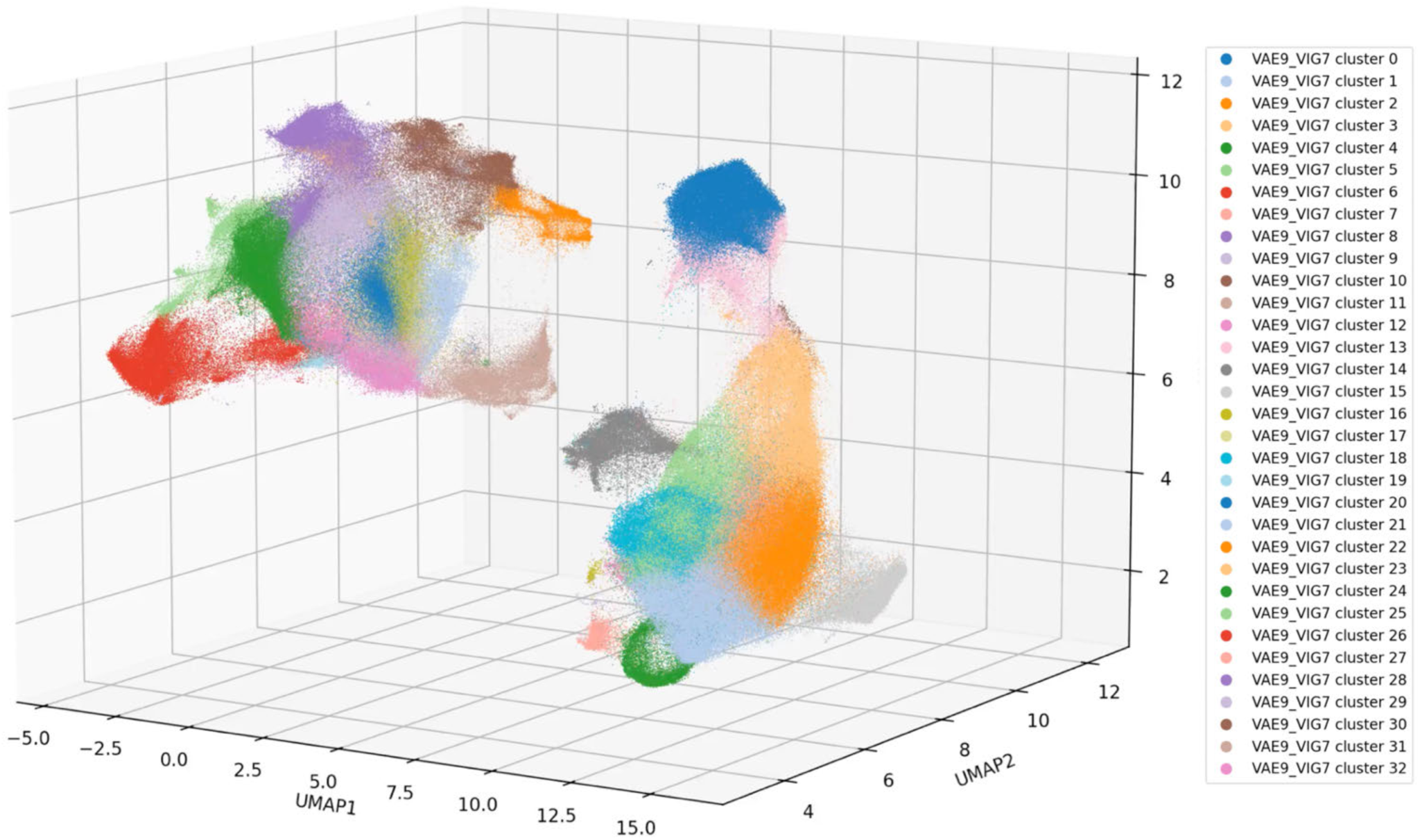
360° UMAP plot rotation showing 9x9µm VAE image patch encodings from CyCIF-1A, CyCIF-1B, and CyCIF-2 colored by Leiden cluster label.

**Extended Data Fig. 9.**
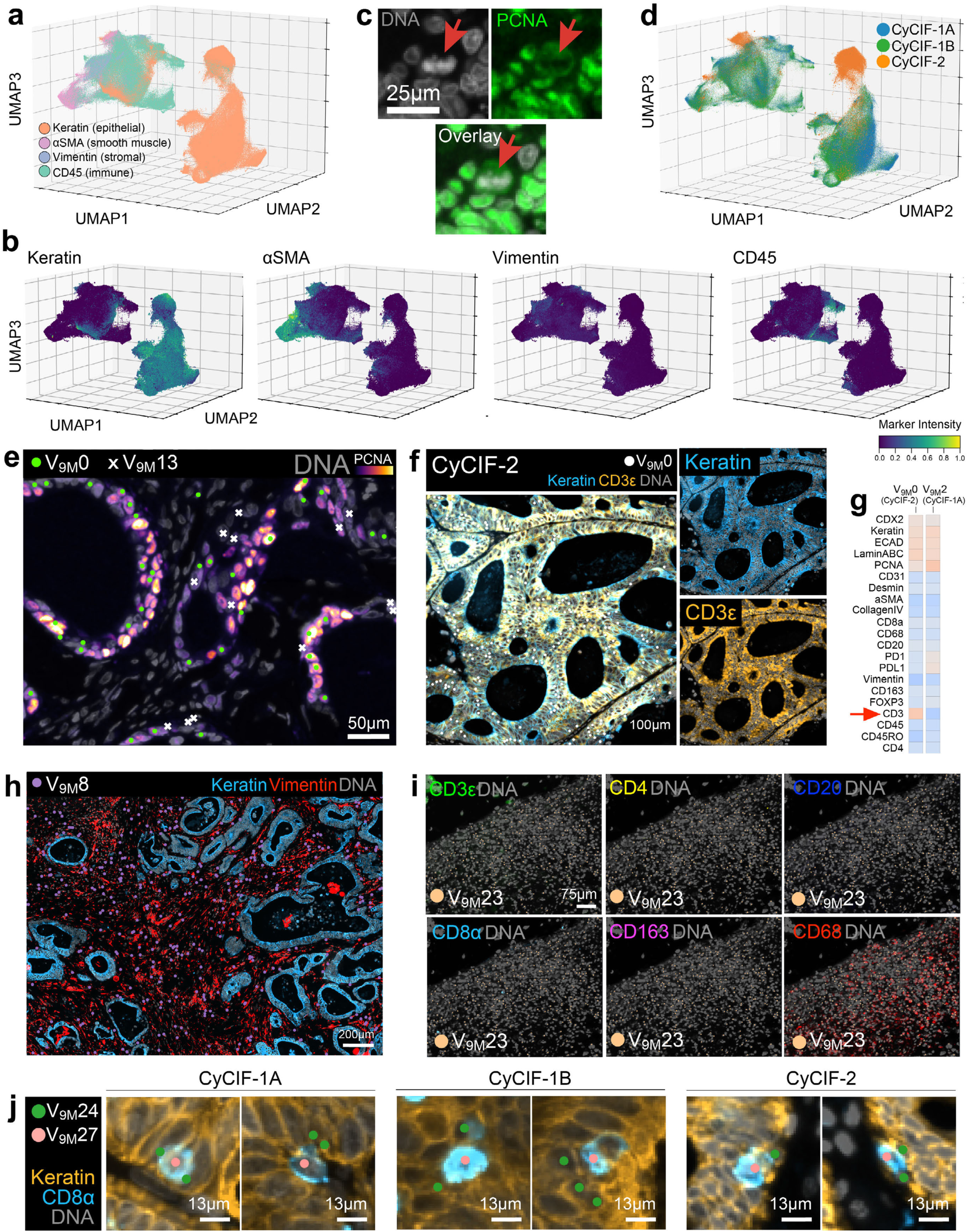
VAE analysis identifies matched cell states within and across patient specimens. **a**, 3D UMAP embedding of 9x9µm VAE image patch encodings from CyCIF-1A, CyCIF-1B, and CyCIF-2 colored by major cell lineage markers: keratin (epithelial, orange), αSMA (smooth muscle, pink), vimentin (stromal, blue), and CD45 (immune, green). **b**. 3D UMAP embedding of VAE image patch encodings from CyCIF-1A, CyCIF-1B, and CyCIF-2 colored by pixel-level median marker intensity for major cell lineage markers keratin, αSMA, vimentin, and CD45. See **Supplementary Fig. 11** for colormaps of all 21 immunomarkers. **c**, Field-of-view in CyCIF-1A showing a tumor cell (red arrow) undergoing cell division as evidenced by its condensed chromatin seen in the DNA channel (gray) and in silhouette in the PCNA channel (green). **d**, 3D UMAP embedding of 9x9µm VAE image patch encodings from CyCIF-1A, CyCIF-1B, and CyCIF-2 colored by specimen; see **Supplementary Video 5** for 360° plot rotation. **e**, Field-of-view in CyCIF-2 showing that V_9M_0 (green circles) and V_9M_13 (white X symbols) both represent glandular AC cells but V_9M_0 cells are brighter for the cell proliferation marker PCNA. **f**, Field-of-view in CyCIF-2 showing glandular AC cells in V_9M_0 (white scatter points). Note that these tumor cells are unexpectedly reactive to anti-CD3ε antibodies. **g**, Channel z-score-normalized protein expression signatures for glandular AC cells in V_9M_0 in CyCIF-2 and analogous cells in V_9M_2 in CyCIF-1A showing that tumor cells in CyCIF-2 are brighter for CD3ε (red arrow) due to non-specific anti-CD3ε antibody binding. **h**, Field-of-view in CyCIF-2 showing fibroblasts in V_9M_8 (purple scatter points) and revealing a large ratio of stromal (vimentin, red)-to-tumor (keratin, blue) tissue. **i**, Field-of-view in CyCIF-2 showing a region of tissue in which CD45^+^ cells are negative for other, more specific, immune lineage markers. **j**, Examples of CD8^+^ T cells and adjacent keratin^+^ tumor cells in CyCIF-1A (left), CyCIF-1B (center), and CyCIF-2 (right) showing that a VAE correctly and reproducibly identifies these neighboring cell types that co-cluster in segmentation-based analysis due to spatial crosstalk.

**Supplementary Fig. 11.**
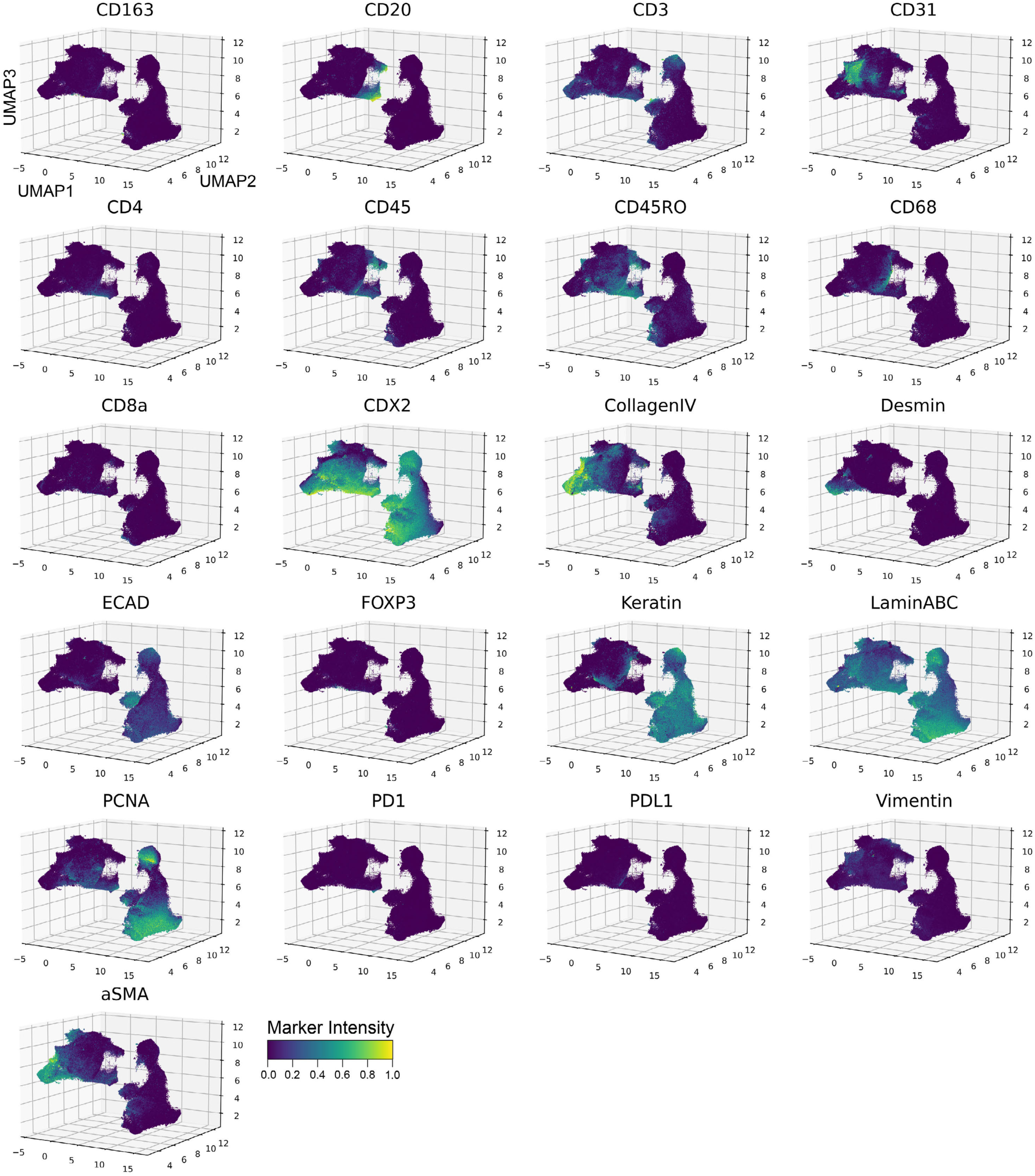
Channel colormaps for 9x9µm VAE image patch encodings from multi-tissue analysis. UMAP embeddings of VAE image patch encodings from CyCIF-1A, CyCIF-1B, and CyCIF-2 colored by pixel-level median marker intensity for 21 immunomarkers.

**Supplementary Fig. 12.**
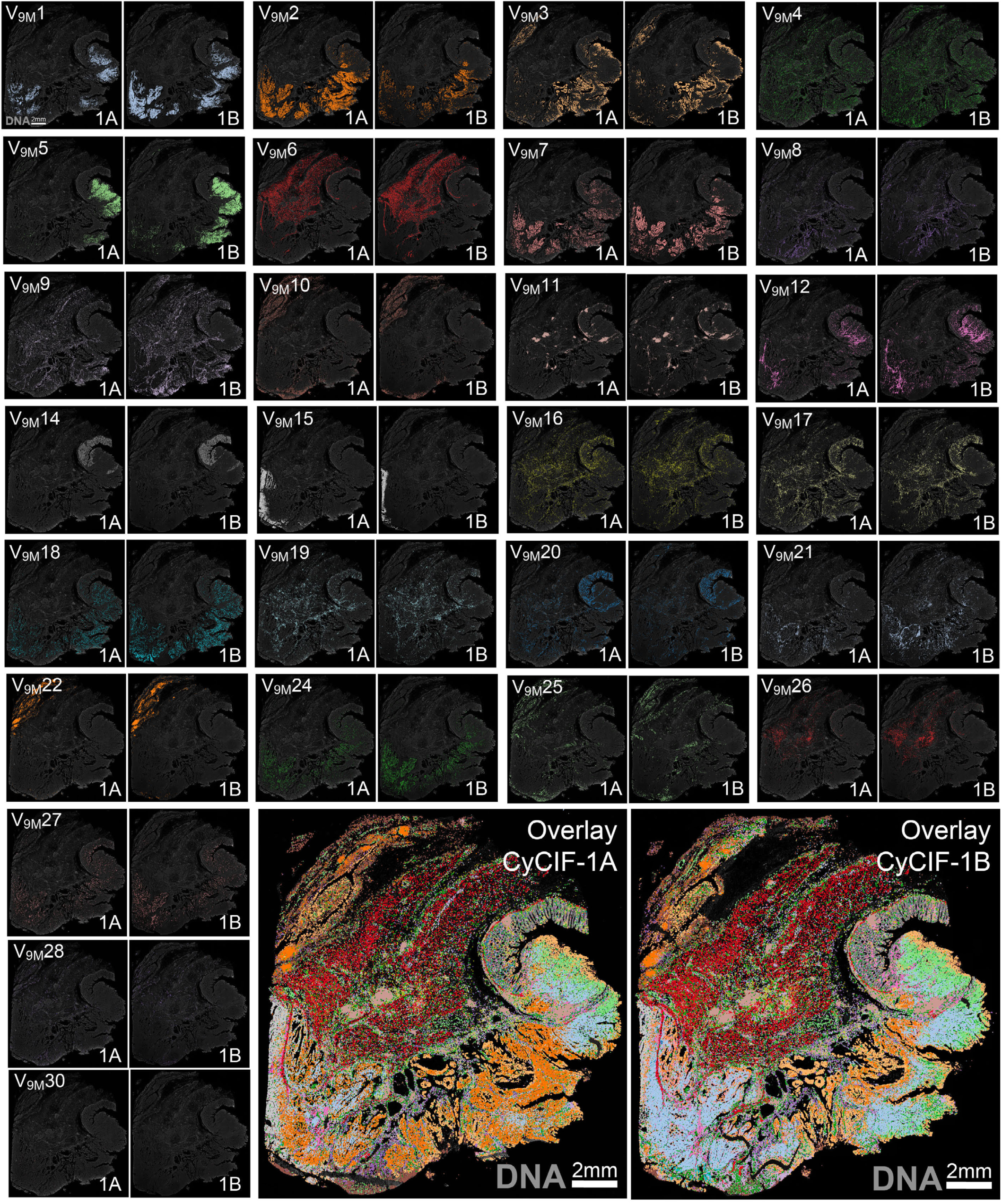
9x9µm VAE image patch Leiden clusters exhibit concordance between serial sections CyCIF-1A and CyCIF-1B. Location of V_9M_ clusters in CyCIF-1A and CyCIF-1B revealing overlap in their histological regions; DNA (gray background) shown for reference.

**Supplementary Video 5.**
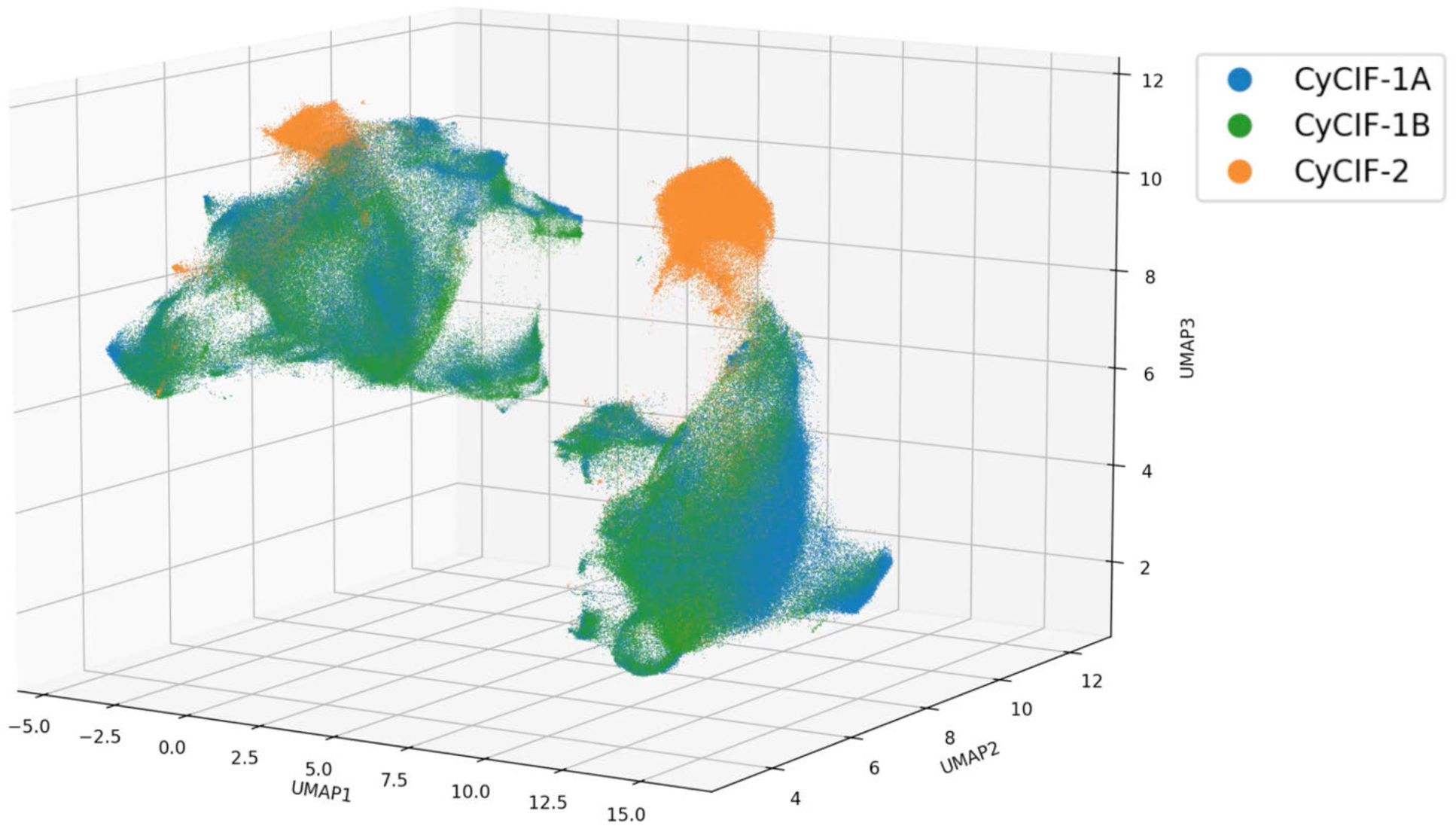
360° UMAP plot rotation showing 9x9µm VAE image patch encodings from CyCIF-1A, CyCIF-1B, and CyCIF-2 colored by tissue specimen.

**Extended Data Fig. 10.**
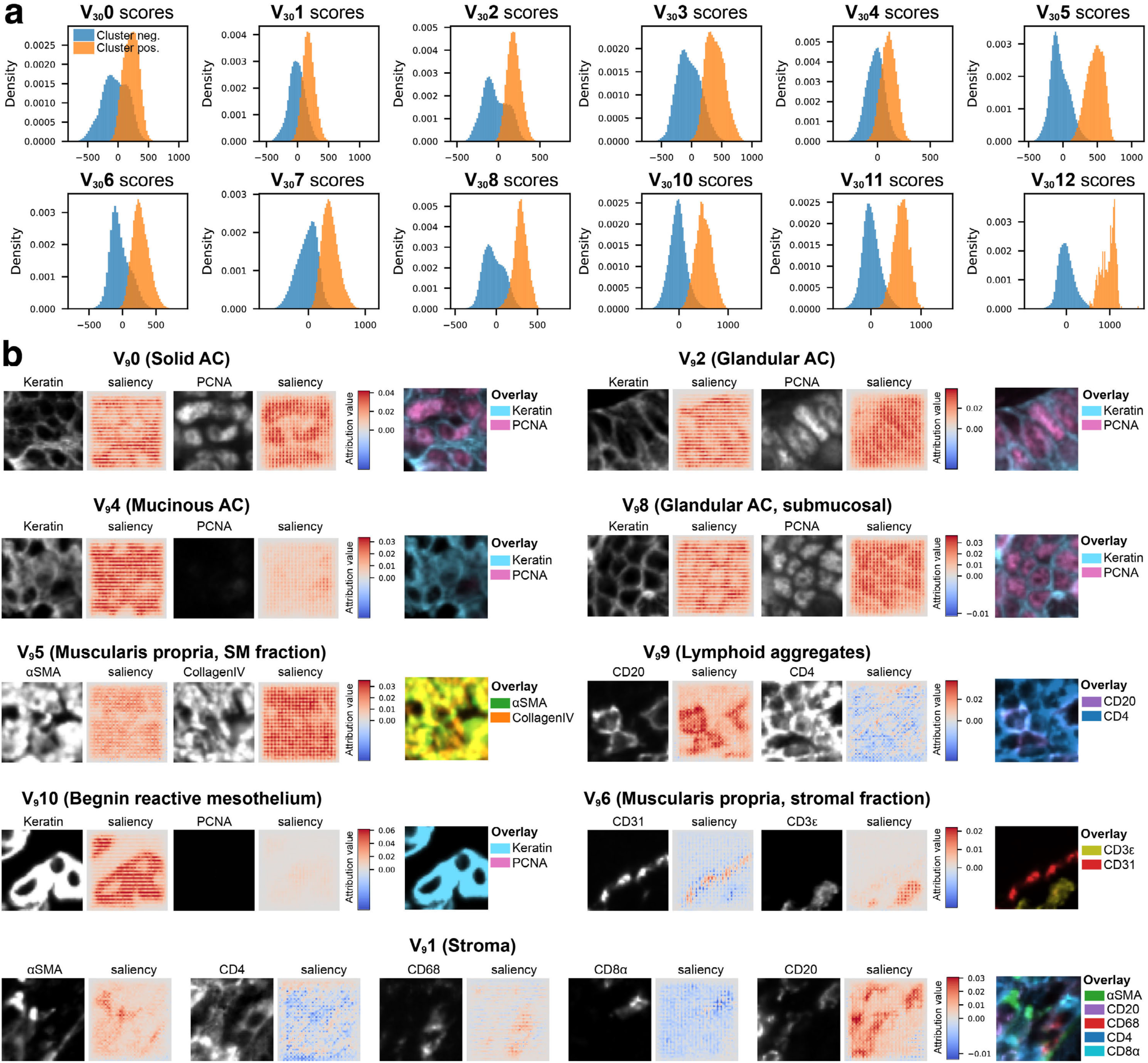
Concept saliency analysis for 30x30µm VAE clusters. **a**, Distributions of concept scores (Sc), defined as the dot product between the i^th^ VAE image patch encoding (*Zi*) and the corresponding cluster concept vector (*Zc*) such that *Sc = Zi ⋅ Zc*, for patches with (orange) and without (blue) a given cluster label. **b**, Channel intensities (left images) and corresponding concept saliency maps (right images) for nine different V_30_ clusters. Overlay color images at far right shown for reference; see **Supplementary Image Gallery 4** for fifteen examples of saliency maps across all 21 channels for each of the twelve V_30_ clusters.

**Supplementary Table 6.** 30x30µm VAE image patch Leiden cluster annotations.

| Leiden cluster | % total cells | Histological region (HR) |
| --- | --- | --- |
| 0 | 16.8 | HR7 (solid AC) |
| 1 | 15.1 | HR9 (peritumoral stroma) |
| 2 | 14.7 | HR5 (glandular AC) |
| 3 | 10.7 | HR4 (subserosa), HR8 (ulcerated submucosa) |
| 4 | 10.5 | HR6 (mucinous AC) |
| 5 | 8.8 | HR2 (muscularis propria, smooth muscle) |
| 6 | 7.8 | HR2 (muscularis propria, stroma) |
| 7 | 5.1 | HR1 (normal mucosa) |
| 8 | 5.0 | HR 5 (glandular AC, predominantly submucosal) |
| 9 | 4.5 | HR* (lymphoid aggregates) |
| 10 | 1.0 | HR3 (serosa) |
| 11 | 0.1 | Microscopy artifact |

**Supplementary Video 6.**
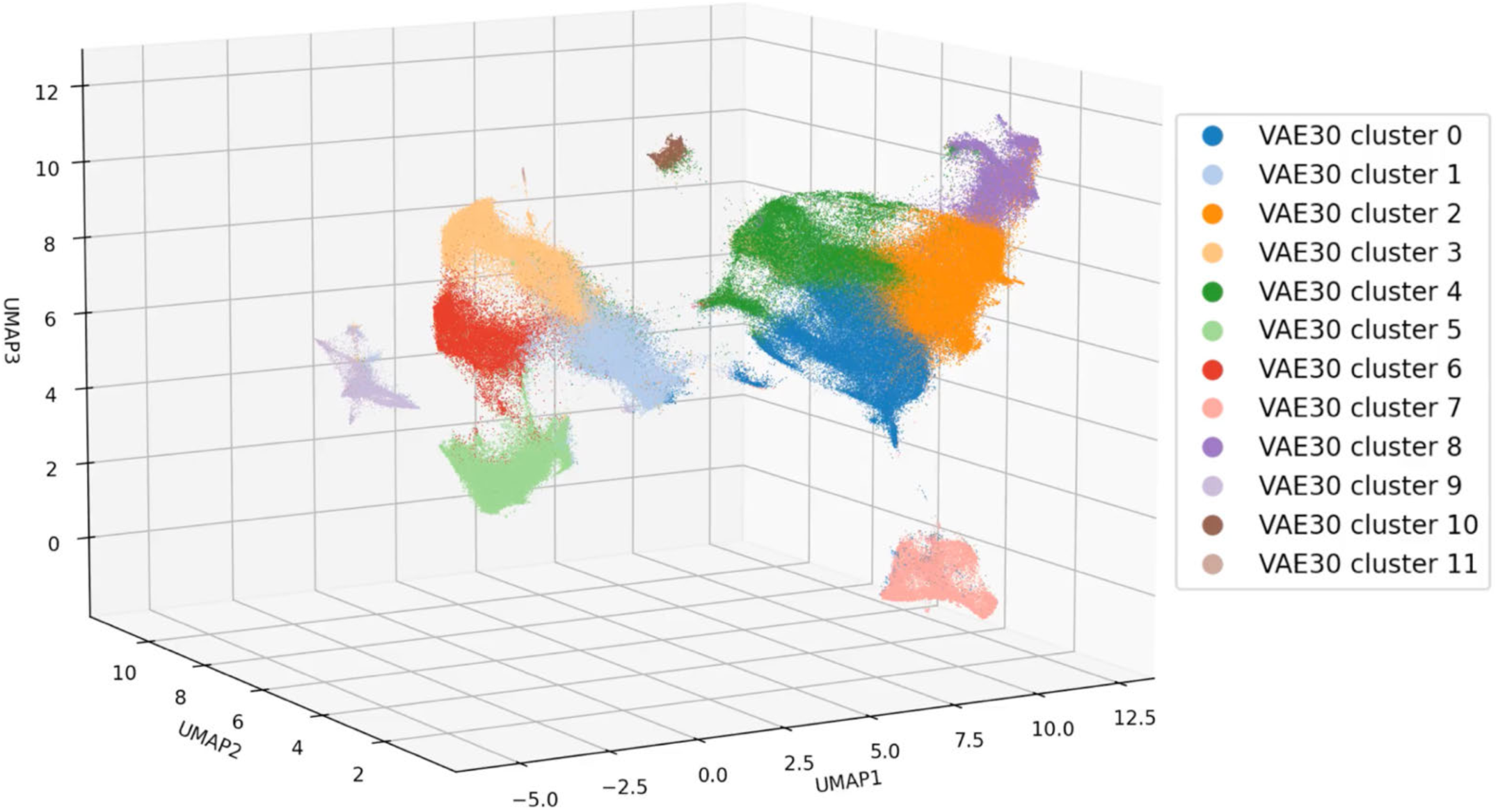
360° UMAP plot rotation showing 30x30µm VAE image patch encodings from CyCIF-1A colored by Leiden cluster labels.

## MATERIALS AND METHODS

### Provenance and Characteristics of Tissue used in this Study

The colorectal cancer (CRC) specimens referred to in this paper as “CyCIF-1A” (HTAN biospecimen ID: HTA13_1_101), “CyCIF-1B” (HTAN biospecimen ID: HTA13_1_106), and “H&E” (HTAN biospecimen ID: HTA13_1_100) consist of 5µm-thick, formalin-fixed paraffin-embedded (FFPE) tissue sections, derived from the same tumor block (referred to as CRC1). These specimens were acquired through the Cooperative Human Tissue Network (CHTN) and were first described in a prior study supported by the Human Tumor Atlas Network (HTAN)^35^. The tumor originated from a right-sided, poorly differentiated stage IIIB colorectal adenocarcinoma (pT3N1bM0) of the cecum, taken from a 69-year-old Caucasian male with microsatellite instability (MSI-H) and a BRAFV600E (c.1799T>A) mutation. The CRC specimen referred to as “CyCIF-2” (HTAN ID: HTA7_922_2) was obtained from Brigham and Women’s Hospital (BWH) in compliance with all relevant ethical regulations established by the BWH Institutional Review Board (IRB). This sample, also analyzed as part of the same HTAN study^35^, represents a stage IVA adenocarcinoma (pT4aN1bcM1a) from a 62-year-old Caucasian male with p53 and APC mutations. See (**Supplementary Table 1**) for an overview of biospecimen metadata. The Lunaphore COMET and Akoya PhenoCycler images are specific to this study and can be accessed upon request from the authors.

### Image Processing

CRC tissue used in this study underwent 10 rounds of four-color multiplex immunofluorescence imaging by CyCIF^56^. DNA plus 3 immunomarkers were imaged at 20x magnification (0.65µm/pixel) with 2x2 pixel binning at each cycle, for a total of 40 channels (**Supplementary Table 2**). Nineteen (19) channels were withheld from analysis in the current study, as they represented either cycle-specific Hoechst DNA stain (channels 1, 5, 9, 13, 17, 21, 25, 29, 33, 37), secondary antibodies alone (channels 2, 3, 4, 6, 7, 8), or channels determined to be of insufficient quality. Raw microscopy tiles (in .rcpnl format) were stitched, registered, segmented, and quantified using the MCMICRO image processing pipeline^28^. Cells in the image were quantified according to three different cell segmentation mask objects: nucleus, cytoplasm, and plasma membrane. To ensure the most accurate segmentation-based signal quantification, the choice of mask selection was determined on a per immunomarker basis according to prior knowledge of the immunolabeling pattern for that marker. The raw single-cell spatial feature table data consisted of 1,218,752 total segmentation instances, which were reduced to 962,343 after QC with CyLinter software^30^ to remove cells affected by visual and image-processing artifacts (e.g., tissue folds, slide debris, illumination aberrations, mis-segmentation, etc.).

### Detailed Experimental Protocols

The following two protocols were used in the processing of tissue for this study: 1) FFPE Tissue Pre-treatment before t-CyCIF on Leica Bond RX V.2 dx.doi.org/10.17504/protocols.io.bji2kkge); 2) Tissue Cyclic Immunofluorescence (t-CyCIF) V.2 dx.doi.org/10.17504/protocols.io.bjiukkew)

### Cellcutter

Cellcutter (https://github.com/labsyspharm/cellcutter) is a Python module developed in our lab for extracting square image patches of arbitrary pixel size centered on individual cells from larger multi-channel TIFF/OME-TIFF images by referring to their corresponding cell ID masks. Patches are stored in the efficient Zarr file format. In this study, Cellcutter was instantiated as a subprocess in the MORPHAEUS data analysis pipeline (https://github.com/labsyspharm/vae) by passing the following inputs to the tool’s *cut_cells* command (see Cellcutter’s README.md file at https://github.com/labsyspharm/cellcutter#readme for further implementation details):

- -window-size 14 (for 9x9µm patches), 46 (for 30x30µm patches)
- -cells-per-chunk 1
- -cache-size 57711
<path to CRC OME-TIFF>
<path to CRC cell ID mask>
<path to CRC spatial feature table>
<path to Zarr output file>
- -channels ["anti_CD3", "anti_CD45RO", "Keratin_570", "aSMA_660", "CD4_488", "CD45_PE", "PD1_647", "CD20_488", "CD68_555", "CD8a_660", "CD163_488", "FOXP3_570",
"PDL1_647", "Ecad_488", "Vimentin_555", "CDX2_647", "LaminABC_488", "Desmin_555", "CD31_647", "PCNA_488", "CollagenIV_647"]

### Image Patch Preprocessing

Cellcutter image patches were centered on the nucleus of individual cells identified during image processing with MCMICRO^28^ and stored as Zarr arrays (https://zarr.readthedocs.io/en/stable/index.html) with dimensions [channels, cells, x_dim, y_dim]. Image patches contained the following channels (in order): CD3ε, CD45RO, keratin, αSMA, CD4, CD45, PD1, CD20, CD68, CD8α, CD163, FOXP3, PD-L1, ECAD, vimentin, CDX2, laminABC, desmin, CD31, PCNA, collagenIV. Pixel intensity values for each channel were log_10_-transformed. Pixels dimmer than the 0.17^th^ and brighter than the 99.99^th^ percentiles of signal intensity for full CRC image channels were clipped and normalized to a range between 0-1 prior to model training to mitigate the impact of excessively dim and bright outlier pixels.

### VAE Network Implementation

Model training was performed using Tensorflow 2.15.0. Batches of 64 image patches were converted from 16-bit unsigned integer to 32-bit floating point format prior to entering to the VAE network for both the 9x9µm and 30x30µm analyses. The VAE architecture was composed of four 2D convolutional layers. The filter size for the first layer was 32 pixels and the subsequent three were 64 pixels. All filters used the ReLU activation function with a kernel size of 3 pixels. Convolutional layers were followed by a 412-dimensional dense layer (in 9x9µm analysis) from which latent means and (log) variances were computed per image patch. A custom variational layer was used to calculate the loss between input image patches and their corresponding learned reconstructions defined as the mean squared error (MSE) between multi-channel input image patches and their learned reconstructions, plus the latent space structure loss (i.e., deviation from a multivariate Gaussian distribution) computed as the Kullback-Leibler (KL) divergence, the coefficient of which was -5.0x10^-4^. The root mean squared propagation (RMSprop) algorithm was used for model optimization and the model learning rate was set to 1.0x10^-6^. Training image patches were shuffled prior to each epoch of model training. Model weights were saved only when validation loss decreased relative to the preceding epoch. To make the VAE encoding and decoding networks end-to-end differentiable, the “reparameterization trick” was employed to enable the standard Gaussian (with a mean of zero and a standard deviation of one) to be the reference distribution in the KL divergence calculation.

### Reporting Summary

Further information on research design is available in the Nature Portfolio Reporting Summary linked to this article.

### Ethics and IRB Statement

HTAN tissue specimens and metadata used in this study complied with all relevant ethical regulations. The research described in this study is considered Non-Human Subjects Research.

### Data Availability Statement

Imaging data analyzed in this paper are stored in a public Amazon S3 bucket (<u>s3://lsp-public-data/baker-2025-vae/</u>) and can be accessed by following instructions under “Downloading input data files” at the dedicated GitHub repository for this paper (https://github.com/labsyspharm/vae-paper). Primary imaging data for all four tissues is also available through the HTAN Data Portal (https://data.humantumoratlas.org/explore) and can be accessed by searching for their respective HTAN biospecimen IDs: CyCIF-1A (HTA13_1_101), CyCIF-1B (HTA13_1_106), H&E (HTA13_1_100), CyCIF-2 (HTA7_922_2). With a free account, access to supplementary videos and image galleries is provided at the dedicated Sage Synapse Wiki page for this paper: https://www.synapse.org/Synapse:syn53216852/files/.

### Code Availability Statement

Source code for the MORPHAEUS data analysis pipeline is available for academic re-use under the MIT open-source license agreement at Github (https://github.com/labsyspharm/vae) and is archived on Zenodo at XXX (to be indexed post-review). Python code used to generate the figures shown in this paper is available at (https://github.com/labsyspharm/vae-paper) and archived on Zenodo at XXX (to be indexed post-review).

### Author Contribution Statements

G.J.B., A.S., and E.N. conceived and designed the study. H.P., and P.K.S. supervised and funded the work. G.J.B. developed and implemented the MORPHAEUS data analysis pipeline. E.N., Y.-A.C., Z.A., S.A.C., S.H., and C.Y. assisted with model development, data analysis, and visualization. G.J.B. and C.H. conceived of and developed the Cellcutter software. G.N.J and FY provided the Lunaphore COMET dataset. S.C. and S.S. provided histopathological annotations. G.J.B. performed data interpretation and generated the figures. G.J.B., E.N., S.S. and P.K.S. wrote the manuscript with input from all authors.

## Acknowledgements

This work was supported by NCI grant U01-CA284207, the Harvard Ludwig Center (P.K.S., S.S.), an ASPIRE Award from The Mark Foundation for Cancer Research, and the David Liposarcoma Research Initiative, and was initiated as part of the computational toolbox for the Human Tissue Atlas Network (HTAN). Histopathology support was provided by P30-CA06516. S.S. is supported by the BWH President’s Scholars Award. We thank J. Tefft for superb editorial support. We also thank Paul Lambert and Taja Lozar from the McArdle Laboratory for Cancer Research at the University of Wisconsin-Madison for kindly providing the Akoya PhenoCycler dataset. Results shown in this study are in part based upon data generated by HTAN (https://humantumoratlas.org/).

## Prefix definitions

S: denotes a segmentation-based cluster
V_9_: denotes a 9x9µm image patch cluster (single-tissue analysis)
V_9M_: denotes a 9x9µm image patch cluster (multi-tissue analysis)
V_30_: denoted a 30x30µm image patch cluster (single-tissue analysis)

